# Glucose stress causes mRNA retention in nuclear Nab2 condensates

**DOI:** 10.1101/2022.01.30.478372

**Authors:** Stephanie Heinrich, Maria Hondele, Désirée Marchand, Carina Patrizia Derrer, Mostafa Zedan, Alexandra Oswald, Liliana Malinovska, Federico Uliana, Sarah Khawaja, Roberta Mancini, David Grunwald, Karsten Weis

## Abstract

Nuclear mRNA export via nuclear pore complexes is an essential step in eukaryotic gene expression. Although factors involved in mRNA transport have been characterized, a comprehensive mechanistic understanding of this process and its regulation is lacking. Here, we use single-RNA imaging in yeast to show that cells use mRNA retention to control mRNA export during stress. We demonstrate that upon glucose withdrawal the essential RNA-binding factor Nab2 forms RNA-dependent condensate-like structures in the nucleus. This coincides with a reduced abundance of the DEAD-box ATPase Dbp5 at the nuclear pore. Depleting Dbp5, and consequently blocking mRNA export, is necessary and sufficient to trigger Nab2 condensation. The state of Nab2 condensation influences the extent of nuclear mRNA accumulation and can be recapitulated *in vitro*, where Nab2 forms RNA-dependent liquid droplets. We hypothesize that cells use condensation to regulate mRNA export and to control gene expression during stress.

**Highlights:**

- The nuclear poly(A)-binding protein Nab2 forms RNA-containing condensate-like structures upon glucose starvation and upon acute cellular depletion of the DEAD-box ATPase Dbp5
- The Nab2 multimerization interface but not the intrinsically disordered regions (IDRs) are essential for condensation *in vitro* and *in vivo*
- Glucose stress leads to poly(A) RNA retention in the nucleus, which is affected by the state of the Nab2 condensate

## Introduction

Gene expression in eukaryotic cells is intricately regulated and aided by subcellular compartmentalization. The nuclear envelope serves as a physical barrier and separates nuclear transcription from cytoplasmic translation processes. Embedded in the nuclear membrane are nuclear pore complexes (NPC) that regulate the passage of macromolecules into and out of the nucleus. This includes the export of messenger RNAs (mRNA) that relay the genetic information to the protein translation machinery in the cytoplasm (reviewed in ^1,2^). mRNAs are extensively processed in the nucleus and undergo a coordinated series of modifications before export, which leads to the addition of a 5′ m7G cap structure, removal of introns through splicing, and 3′ end poly(A) tail synthesis (reviewed in ^3–5)^. To facilitate these processes, a range of RNA-binding proteins (RBPs) form a dynamic mRNA coat on the nascent mRNA, either co- or post-transcriptionally in the nucleoplasm. The orderly assembly of this coat is essential for forming an export-competent messenger ribonucleoprotein (mRNP) particle (reviewed in ^1,6^). Early binding proteins include the cap-binding complex (CBC), comprising Cbc1 (CBP80 in vertebrates) and Cbc2 (CBP20), which binds co-transcriptionally to the 5’-cap of nascent mRNAs^7^. Another early binding complex, the <u>tr</u>an-scription and <u>ex</u>port (TREX) complex (consisting of the multi-subunit THO subcomplex, the DEAD-box ATPase Sub2 (UAP56), and the RNA binding protein Yra1 (ALYREF)) connects 3′ end processing with mRNA export ^8–11^. Similar to TREX, the abundant class of serine-arginine-rich (SR) proteins (e.g., yeast Npl3) has been implicated in a multitude of mRNA processing steps, such as co-transcriptional splicing, 3′ end processing and nuclear export (reviewed in ^12^). Furthermore, the RNA-binding protein Nab2 (ZC3H14) is recruited to the poly(A) tail of the pre-mRNA and is involved in nuclear mRNA surveillance protecting mRNAs against degradation by the nuclear exosome^13–16^. Nab2 has been shown to play a role in proper 3′ end processing of nascent transcripts to terminate polyadenylation and facilitate compaction of the mRNP^17–19^. In addition, it directly interacts with the NPC component Mlp1^20^ and has been shown to shuttle between nucleus and cytoplasm, getting re-imported to the nucleus by the karyopherin Kap104 ^21^. The translocation of mature transcripts through NPCs is mediated by the export factor Mex67:Mtr2 (NXF1:NXT1), which functions akin to a mobile nuclear pore protein facilitating transport by binding to both mRNPs as well as phenylalanine–glycine (FG)-containing NPC components (nucleoporins)^1,22^. On the cytoplasmic face of the NPC, the essential DEAD-box ATPase Dbp5 (DDX19B), aided by its interaction partners Nup159 (NUP214), Gle1 and inositol hexakisphosphate (IP_6_), was proposed to remodel emerging mRNPs by removing factors such as Nab2 from the mRNA, thereby facilitating their release into the cytoplasm^23–25^. However, these models are mainly based on *in vitro* studies^24,26,27^, and there are only a few reports that address the molecular mechanism(s) of mRNA export and its regulation *in vivo* ^28–30^.

In addition to membrane-based compartmentalization, cells also organize their content in membraneless organelles, or ‘biomolecular condensates’, through protein de-mixing, also referred to as phase separation (reviewed in ^31,32^). Formation of these biomolecular condensates depends on weak, multivalent interactions and proteins able to form condensates often contain low-complexity domains with intrinsically disordered regions (IDRs). IDRs are frequently found in RNA-binding proteins and RNA can enhance their propensity to form condensates (reviewed in ^33^). Condensation is an evolutionarily ancient and conserved self-organizing principle ^34,35^. Within a cell, many structures can be defined as membraneless organelles and they have been implicated in various aspects of gene expression regulation (reviewed in ^31^). For example, stress conditions like nutrient depletion or acute temperature shifts have been shown to induce the formation of cytoplasmic condensates such as P-bodies (PB) or stress granules (SG)^36,37^. If condensation plays a role in mRNA export is unknown.

Here, we report that blocking mRNA export through depletion of the DEAD-box ATPase Dbp5 or through glucose withdrawal results in the formation of nuclear condensate-like structures containing the mRNA-binding factor Nab2. Nab2 mutants that affect the material properties of the condensate affect the degree of mRNA retention in the nucleus in an RNA dose-dependent manner. Our results suggest that condensation can modulate mRNA transport during cellular stress and provide mechanistic insights into mRNA export regulation.

## Results

### Acute Dbp5 depletion causes nuclear mRNA accumulation and Nab2 mobility reduction

To investigate mRNA export *in vivo*, we employed an auxin-inducible acute protein depletion system in the budding yeast *Saccharomyces cerevisiae* (Nishimura 2009, Fig. 1A, ‘degron’) and characterized the essential DEAD-box ATPase Dbp5, a proposed mRNP remodeler and ‘motor’ of directional transport, located at the cytoplasmic side of the NPC^23–25^. Addition of auxin led to the efficient and specific depletion of Dbp5 with a half-time of approximately 12 min (Fig. S1A,B), without affecting the overall NPC composition based on the quantification of selected nucleoporins using a previously established method (NuRIM ^38^; Fig. S1C). Dbp5 depletion did not affect protein export as an NLS-NES-2xGFP reporter did not enrich in the nucleus after auxin addition. In contrast, blocking Xpo1-dependent export using Leptomycin B (LMB) in an LMB-sensitive Xpo1 mutant ^39^ strongly impaired nuclear export of the GFP reporter (Fig. S1D-G). Furthermore, addition of LMB to cells that were already Dbp5-depleted shifted the GFP reporter distribution towards the nucleus. This shows that nuclear protein import and export remain functional when Dbp5 is depleted (Fig. S1D-G).

**1).**
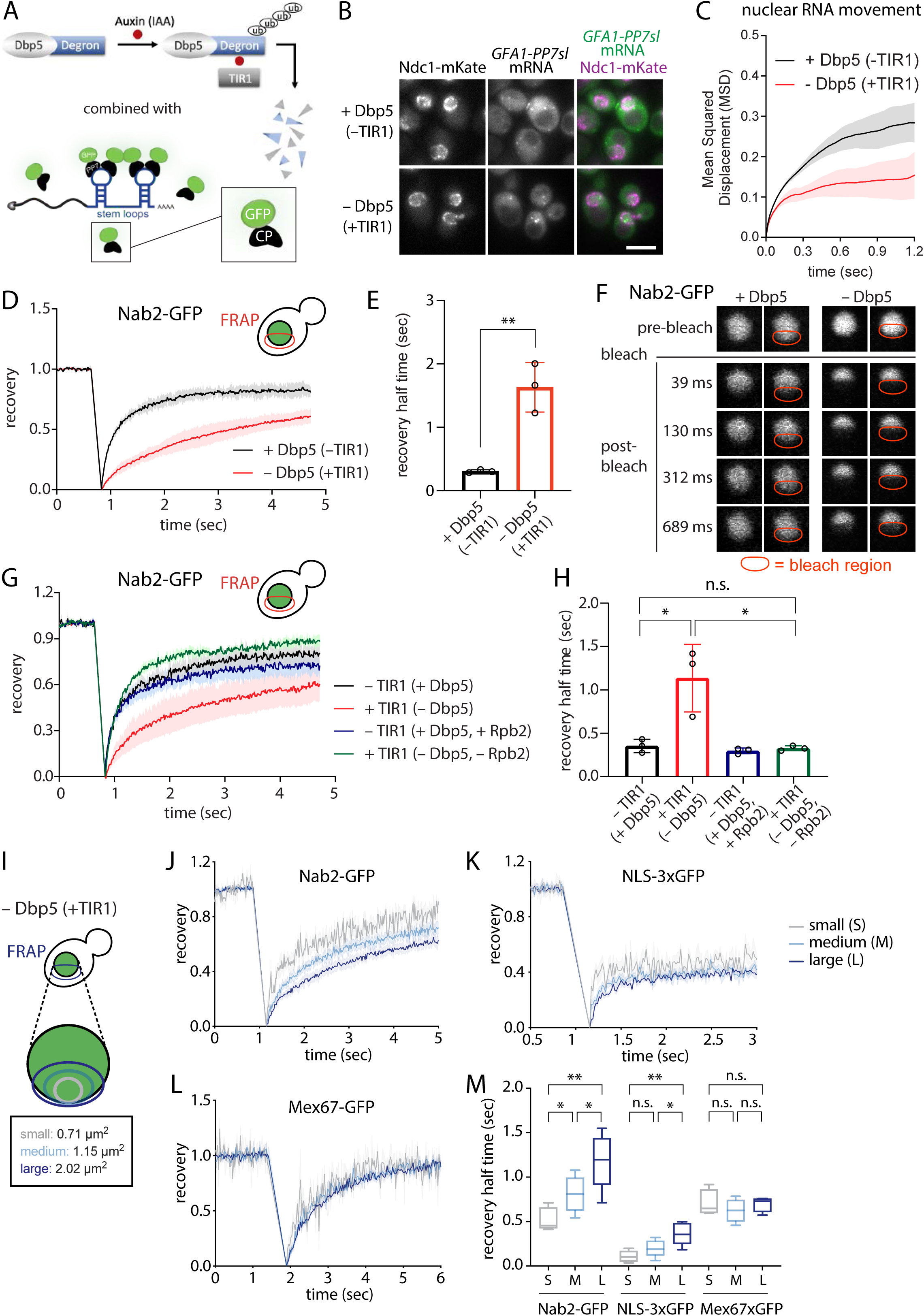
Acute Dbp5 depletion causes nuclear mRNA accumulation and Nab2 condensation. A) Schematic representation of the auxin-inducible protein depletion tool and the single RNA visualization in vivo. (Upper) Upon addition of Auxin, degron-tagged Dbp5 gets recognized by TIR1 and targeted for proteasomal degradation ^85^; (Lower) 24 PP7 stem loops (PP7sl) are integrated after the STOP codon of a low-copy number gene of interest. A second locus drives the expression of GFP-tagged PP7 coat protein (PP7CP-GFP) that can bind to PP7sl with high affinity, thereby allowing single RNA visualization ^42^. B) Cells expressing GFA1-PP7sl, PP7CP-GFP, the nuclear rim marker Ndc1 and the Dbp5-degron with or without TIR1 are treated for 1 hr with auxin (IAA) before imaging using a custom-built microscope ^43^. Scale bar 5 µm. C) Quantification of mean squared displacement (MSD) of PP7-labeled GFA1 mRNA in cells from B). Images were taken every 15 ms. Shown are means ± SD of three biological replicates with ≥265 RNA particles per condition. D) FRAP of Nab2-GFP cells expressing the Dbp5-degron with or without TIR1 treated for 2 hr with auxin (IAA). FRAP curves are normalized to prebleach and postbleach values. Mean curves ± SEM of three biological replicates with ≥16 cells per replicate are shown. E) Half time of recovery retrieved from fitting individual FRAP curves from D) with a single component model. Shown are means ± SEM of three biological replicates. * p=0.028, unpaired t-test. F) Representative FRAP images of a single nucleus from D) G) FRAP of Nab2-GFP cells expressing either only the Dbp5-degron or both the Dbp5-degron and the Rpb2-degron with or without TIR1 treated for 2 hr with auxin (IAA). FRAP curves are normalized to prebleach and postbleach values. Mean curves ± SEM of three biological replicates with ≥15 cells per replicate are shown. H) Half time of recovery retrieved from fitting individual FRAP curves from G) with a single component model. Shown are means ± SEM of three biological replicates. * p=0.0271 (– TIR1(+Dbp5) vs. +TIR1(–Dbp5), * p=0.0232 +TIR1(–Dbp5) vs. +TIR1(–Dbp5, –Rbp2), ns p=0.6109 ((–TIR1(+Dbp5) vs. +TIR1(–Dbp5, –Rbp2)), unpaired t-test. I) Schematic showing sizes of different FRAP bleach areas. J) FRAP of Nab2-GFP cells expressing the Dbp5-degron with TIR1 treated for 2 hr with auxin (IAA). FRAP curves are normalized to prebleach and postbleach values. Mean curves ± SEM of six biological replicates with ≥12 cells per replicate are shown. K) FRAP of NLS-3xGFP cells expressing the Dbp5-degron with TIR1 treated for 2 hr with auxin (IAA). FRAP curves are normalized to prebleach and postbleach values. Mean curves ± SEM of six biological replicates with ≥13 cells per replicate are shown. L) FRAP of Mex67-GFP cells expressing the Dbp5-degron with TIR1 treated for 2 hr with auxin (IAA). FRAP curves are normalized to prebleach and postbleach values. Mean curves ± SEM of three biological replicates with ≥11 cells per replicate are shown. M) Half time of recovery retrieved from fitting individual FRAP curves from F-H) with a single component model. Shown are box-and-whisker plots (min, median, max). * p=0.0124 (Nab2(small) vs. Nab2(medium)), ** p=0.0018 (Nab2(small) vs. Nab2(large)), * p=0.0321 (Nab2(medium) vs. Nab2(large)), ns p=0.0941 (3xGFP(small) vs. 3xGFP(medium)), ns p=0.0941 (3xGFP(small) vs. 3xGFP(medium)), ** p=0.0015 (3xGFP(small) vs. 3xGFP(large)), * p=0.0305 (3xGFP(medium) vs. 3xGFP(large)), ns p=0.4621 (Mex67-GFP(small) vs. Mex67-GFP(medium)), ns p=0.9675 (Mex67-GFP(small) vs. Mex67-GFP(large)), ns p=0.3802 (Mex67-GFP(medium) vs. Mex67-GFP(large)), unpaired t-test.

Upon Dbp5 depletion, we observed a global nuclear retention of poly(A) RNA using fluorescence *in situ* hybridization (FISH) with an oligo(dT)30 probe, similar to previous reports using temperature-sensitive *dbp5* alleles (Fig. S1H)^40,41^. To investigate whether this oligo(dT)30 staining corresponds to the nuclear retention of mRNA, we examined the export of individual mRNAs. To this end, we marked transcripts of an essential gene, glutamine-fructose-6-phosphate amidotransferase (*GFA1)*, with the PP7-PCP stem loop labeling system (^42^, Fig. 1A) in the Dbp5 degron background. In the presence of Dbp5, the GFA1-PP7 transcripts were mostly localized in the cytoplasm (Fig. 1B). Upon Dbp5 depletion, transcripts accumulated in the nucleus close to the nuclear periphery. Interestingly, the mobility of nuclear mRNAs was reduced in the absence of Dbp5 based on single particle tracking results obtained from a custom-built microscope (^43^, Fig. 1C, Video S1, Video S2). To test whether the dynamics of nuclear export factors were also affected, we performed fluorescence recovery after photobleaching (FRAP) measurements after Dbp5 depletion. The nuclear dynamics of the export factors Yra1 and Npl3 as well as an NLS-GFP(3x) control did not change when Dbp5 was depleted (Fig. S2A-F). However, the nuclear mobility of the poly(A)-binding protein Nab2 was dramatically reduced (Fig. 1D-F), with a >5-fold increase in recovery half-time in the absence of Dbp5. Intriguingly, FRAP of approximately half of the nucleus showed very slow recovery of Nab2 in the entire bleached region (Fig. 1F). The reduction in Nab2 mobility is RNA-dependent as additional depletion of a polymerase II subunit (Rpb2-AID) reverted the phenotype (Fig. 1G,H) by reducing the amount of poly(A) RNA in the nucleus (Fig. S2G).

Dbp5-dependent mobility reduction of Nab2 could suggest the formation of a more viscous, higher-order structure, such as a biomolecular condensate, that spans across the nucleus. Assessing material properties *in vivo* is technically challenging. One critical control is to demonstrate that the rate of FRAP recovery is dominated by diffusion rather than binding. This can be shown by testing the dependency of FRAP recovery on the size of the bleach spot^44^ (Fig. 1I). If nuclear Nab2 mobility is diffusion-dominant, increasing the bleach area would lead to a reduction in FRAP recovery half-times. Inversely, if Nab2-GFP mobility is binding-dominant, recovery half-times should be independent of the bleach area size. Indeed, Nab2-GFP, as well as the positive control, freely diffusing nuclear NLS-3xGFP, showed diffusion-dominant behavior (Fig. 1J,K,M). In contrast, Mex67-GFP, which is a mobile nucleoporin^22^, shows a binding-dominant FRAP recovery (Fig. 1L,M).

Our results therefore suggest that Nab2 forms an mRNA-dependent, condensate-like structure in the nucleus *in vivo*, whose dynamics depend on active mRNA export mediated by Dbp5.

### Glucose stress causes reduction of nuclear Nab2 dynamics and bulk nuclear mRNA retention

Having established that nuclear Nab2 mobility is reduced in an acute Dbp5 depletion condition *in vivo*, we next asked if changes in Nab2 dynamics can also be observed in physiological conditions in cells. It has been previously observed that Dbp5 re-localizes from the NPC to the nucleoplasm in specific stress conditions, such as acute ethanol stress^45,46^. We therefore tested Dbp5 localization upon nutrient stress. Interestingly, NPC localization of Dbp5, but not its cytoplasmic NPC-interaction partners Nup159 and Gle1, is reduced both upon acute (30 min –DEX) and prolonged (24 hrs –DEX) glucose starvation (Fig. 2A,B and S3A,B). In contrast, the major export adaptor Mex67 does not change its localization at the nuclear rim upon glucose withdrawal (Fig. S3C). Furthermore, in these starvation conditions, export of poly(A) mRNA is blocked (Fig. 2C,D). Intriguingly, glucose starvation resulted in the formation of predominantly nuclear Nab2 foci, while other nuclear export factors such as Yra1 and Npl3 were unaffected (Fig. 2C,D and S2H,I). Discrete Nab2 foci formed during short-term glucose withdrawal and focus formation was enhanced upon long-term glucose starvation (Fig. 2C,D). Nab2 foci were mostly resistant to 1,6-hexanediol treatment, which dissolves cytoplasmic membraneless organelles such as P-bodies, but not stress granules (Fig. S3D)^36^. Nab2 focus formation was accompanied by nuclear accumulation of poly(A) RNA in a punctate pattern overlapping with Nab2 foci (Fig. 2C,D and S3E,F). The poly(A) RNA that was retained in these foci seemed to correspond to intact transcripts as they also showed co-localization with the cap-binding complex proteins Cbc1 and Cbc2 (^7^, Fig. S4A). Nab2-mRNA foci were reversible, as release from starvation conditions into medium containing glucose led to the full dispersal of Nab2 in the nucleus within an hour, concomitant with a dispersal of poly(A) RNA foci (Fig. S4B).

**2).**
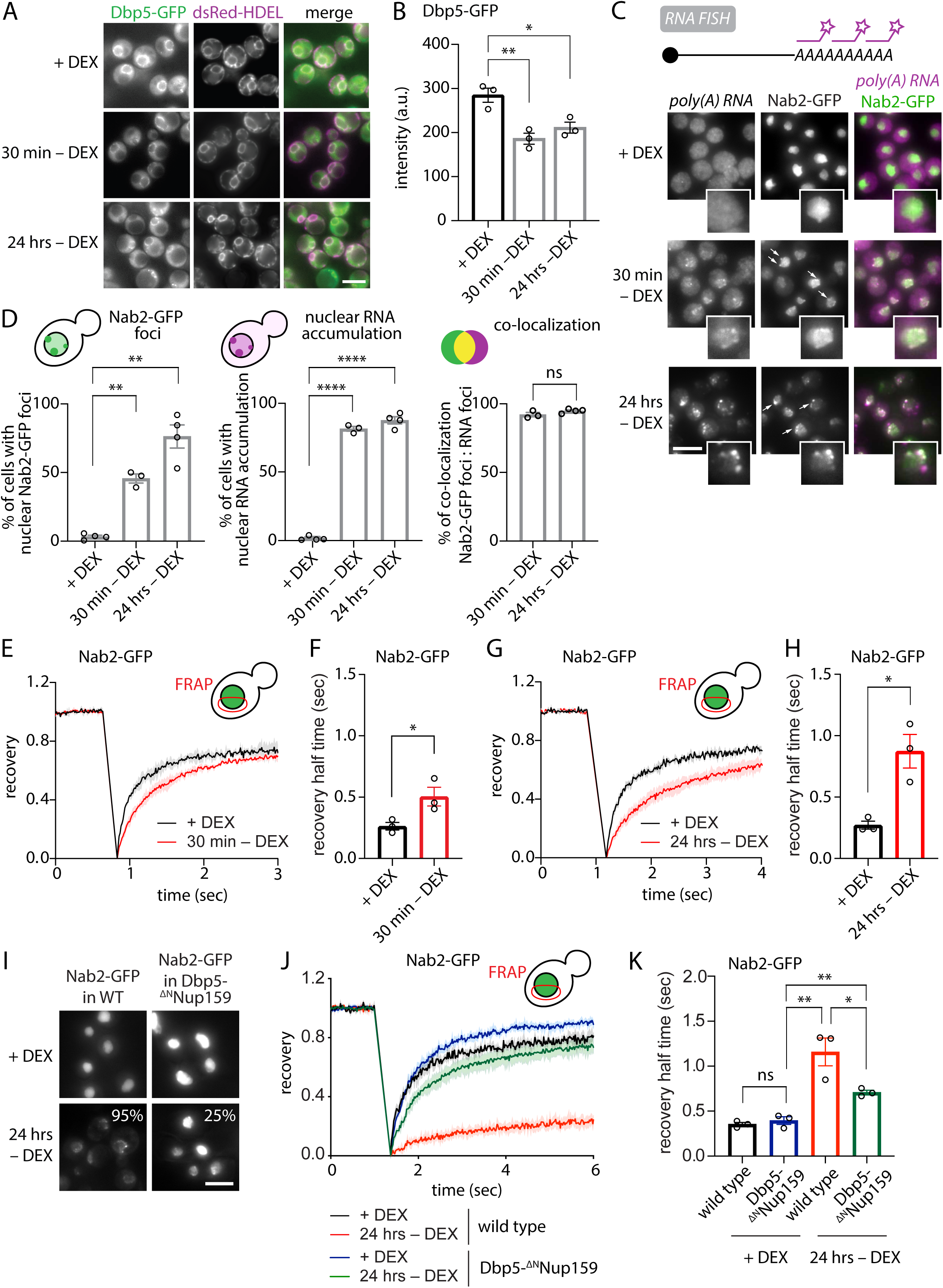
Glucose stress causes reduction of nuclear Nab2 dynamics and nuclear *poly(A)* RNA retention. A) Cells expressing Dbp5-GFP and dsRed-HDEL were grown in medium + DEX or acutely shifted for 30 min or 24 hrs into medium without DEX. Scale bar 5 µm. B) Quantification of Dbp5-GFP intensity at the nuclear envelope by NuRIM in the conditions shown in A). Shown are means ± SEM of three biological replicates. At least 350 cells were analyzed per condition and replicate. Dbp5: ** p=0.0082 (+DEX vs. 30 min –DEX), * p=0.022 (+DEX vs. 24 hrs –DEX), unpaired t-test. C) RNA FISH against poly(A) using an oligo(dT)30 probe was performed in fixed cells expressing Nab2-GFP. Samples were taken in +DEX or after acute starvation in medium without DEX for 30 min or 24 hrs. Insets show single nucleus (2.16x magnification). White arrows indicate Nab2-GFP focus. Scale bar 5 µm. D) Quantification of C). (Left) Nab2-GFP foci, percentage of cells with nuclear GFP foci; (Center) nuclear RNA accumulation, percentage of cells where average poly(A) RNA signal intensity is larger in nucleus than in cytoplasm (includes cells with and without visible RNA foci); (Right) co-localization, percentage of nuclear Nab2-GFP foci co-localizing with poly(A) RNA foci. ** p=0.0032 (Nab2-GFP foci, +DEX vs. 30 min –DEX), ** p=0.0029 (Nab2-GFP foci, +DEX vs. 24 hrs –DEX), **** p<0.0001 (nuclear RNA accumulation, +DEX vs. 30 min –DEX), **** p<0.0001 (nuclear RNA accumulation, +DEX vs. 24 hrs –DEX), ns p=0.2181 (co-localization, 30 min –DEX vs. 24 hrs –DEX), unpaired t-test. E) FRAP of Nab2-GFP cells in +DEX or 30 min –DEX. FRAP curves are normalized to prebleach and postbleach values. Mean curves ± SEM of three biological replicates with ≥15 cells per replicate are shown. F) Half time of recovery retrieved from fitting individual FRAP curves from E) with a single component model. Shown are means ± SEM of three biological replicates. * p=0.04, unpaired t-test. G) FRAP of Nab2-GFP cells in +DEX or 24 hrs –DEX. FRAP curves are normalized to prebleach and postbleach values. Mean curves ± SEM of three biological replicates with ≥15 cells per replicate are shown. H) Half time of recovery retrieved from fitting individual FRAP curves from G) with a single component model. Shown are means ± SEM of three biological replicates. * p=0.01, unpaired t-test. I) Representative images of Nab2-GFP cells in wild type (WT) or Dbp5-^ΔN^Nup159 (‘Dbp5 tether’) background in +DEX or 24 hrs –DEX. Percentages show number of cells with nuclear Nab2-GFP foci (means of two biological replicates). Scale bar 5 µm. J) FRAP of cells in I). FRAP curves are normalized to prebleach and postbleach values. Mean curves ± SEM of three biological replicates with ≥15 cells per replicate are shown. K) Half time of recovery retrieved from fitting individual FRAP curves from J) with a single component model. Shown are means ± SEM of three biological replicates. ns p=0.4425, * p=0.0425, ** p=0.0067 (wild type 24 hrs –DEX vs. Dbp5-^ΔN^Nup159 +DEX), ** p=0.0032 (Dbp5-^ΔN^Nup159 +DEX vs. 24 hrs –DEX), unpaired t-test.

We then tested Nab2 mobility in short-term and long-term starvation. Similar to acute Dbp5 depletion, we observed a reduction in nuclear dynamics after 30 min of acute glucose starvation (Fig. 2E,F). This phenotype was even more prominent upon long-term glucose starvation (24 hrs) (Fig. 2G,H). The reduction in dynamics was specific for Nab2 as the mobility of other export factors such as Npl3 and Yra1 was not affected (Fig. S2J-M). Nab2 mobility reduction was dependent on the presence of mRNA and treatment with the transcription inhibitor thiolutin^47^ restored Nab2 mobility to the levels of non-stressed cells (Fig. S4C,D). We next wanted to examine whether Nab2 focus formation is triggered by the loss of Dbp5 from the NPC during long-term stress. Intriguingly, when Dbp5 was prevented from leaving the NPC by permanently fusing it to its interaction partner Nup159 (‘Dbp5 tether’, Dbp5-^ΔN^Nup159, ^28^ Fig. S4E), Nab2 focus formation was strongly reduced, with the fraction of cells showing observable foci down from 95% to 25% (Fig. 2I). Furthermore, the mobility of Nab2 was only mildly affected by long-term glucose withdrawal in this mutant (Fig. 2J-K). This suggests that reduction of Dbp5 abundance at the NPC in response to glucose starvation is required for Nab2 focus formation.

We conclude that, upon glucose starvation, RNA-dependent Nab2 focus formation coincides with reduction of Dbp5 abundance at the NPC, which can be reversed by permanently tethering Dbp5 to the NPC.

### Nab2 forms condensates *in vitro*

The sequence of Nab2 shows many hallmarks of RNA-binding proteins that can undergo condensation^31,32^. In addition to an N-terminal domain required for Mlp1 binding and mRNA export, Nab2 contains a Q-rich domain of unknown function (QQQP), an RGG domain with an internal nuclear localization signal (NLS), and 7 zinc fingers (ZnF), of which ZnF5-7 are essential for RNA binding (Fig. S4F)^13,48,49^. Furthermore, according to PONDR, a predictor of naturally disordered regions^50,51^, the Q-rich domain and the RGG domain are intrinsically disordered (Fig. S4F). We therefore wanted to examine whether Nab2 can form condensates *in vitro* and purified Nab2 recombinantly from bacteria, in a monomeric eGFP-tagged, soluble and RNA-free form (Fig. S4G). Intriguingly, in the presence of poly(A) RNA recombinant Nab2 formed liquid droplets at close to endogenous concentrations (Fig. 3A and S4H,I). These droplets were dynamic as they had the ability to fuse and grow over time (Video S3), and were mildly sensitive to the aliphatic alcohol 1,6-hexanediol, consistent with them being formed via condensation (Fig. S4J).

**3).**
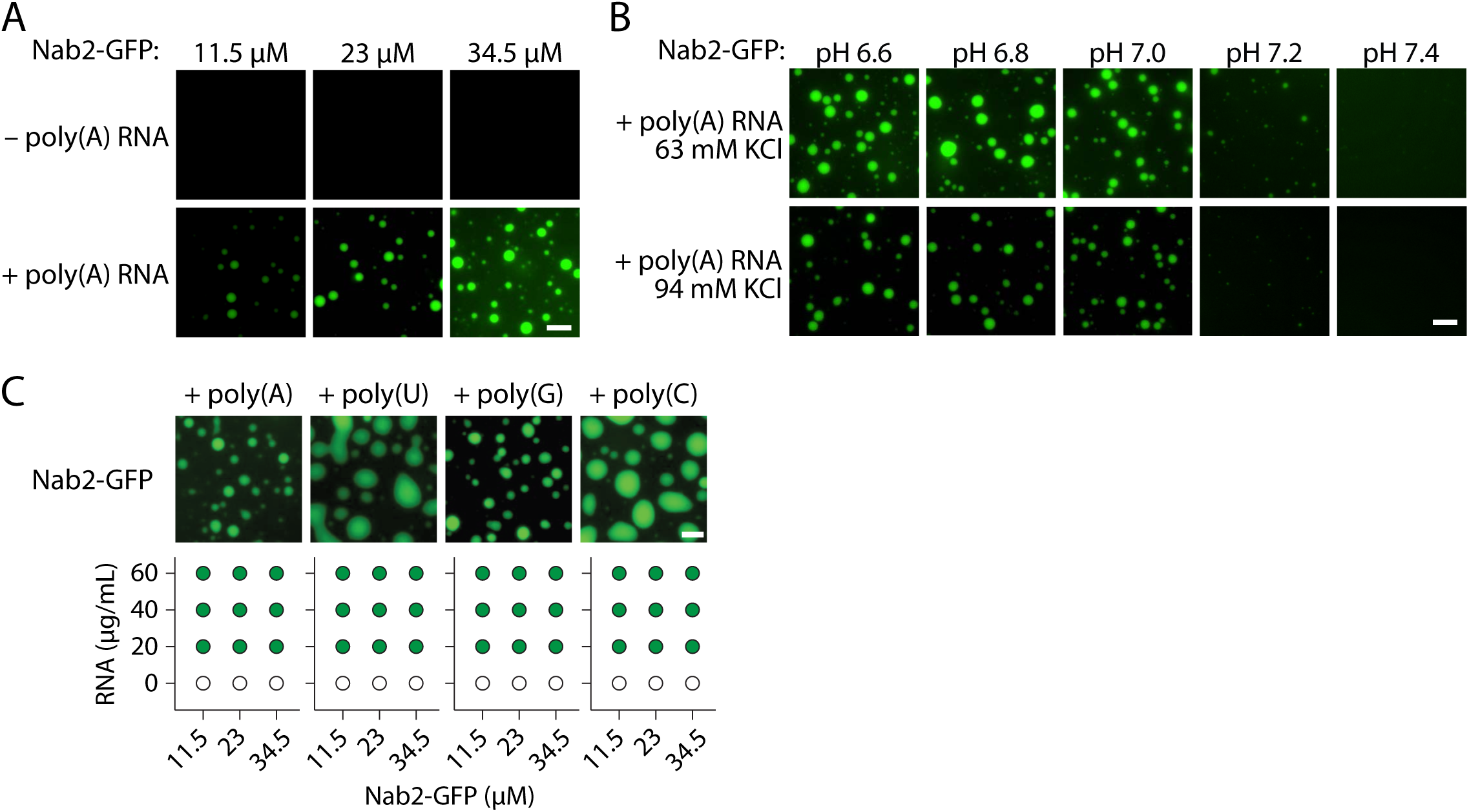
Nab2 phase-separates *in vitro*. A) Recombinant Nab2-GFP droplets were formed in LSB150 buffer to a final KCl concentration of 94 mM in the presence or absence of 0.04 mg/mL poly(A) RNA and imaged at 25 °C 30 min after setup. Scale bar 10 µm. B) Recombinant Nab2-GFP droplets were formed in LSB100 or LSB150 buffers to a final KCl concentration of 63 or 94 mM, respectively, at the indicated pH in the presence of 0.04 mg/mL poly(A) RNA and imaged at 25 °C 30 min after setup. Scale bar 10 µm. C) Representative images and phase diagram of recombinant Nab2-GFP droplets, which were formed in LSB150 buffer (to a final KCl concentration of 94 mM) containing the indicated single-nucleoside polymers and imaged at 25 °C 20 min after setup. Representative images show droplets formed with 0.04 mg/mL single-nucleoside RNA. Scale bar 5 µm.

Nab2 condensates form at physiological salt concentrations and are pH-dependent (Fig. 3B). In addition to poly(A), Nab2 can form condensates with other single-nucleoside polymers (poly(U)/poly(G)/poly(C), Fig. 3C), suggesting that Nab2 does not exclusively bind to the poly(A) tail of mRNAs. Indeed, previous reports have shown that Nab2 can bind also non-poly(A) RNA *in vitro* ^13^ and associates with mRNA regions upstream of the poly(A) tail *in vivo* ^52^.

Our experiments demonstrate that Nab2 forms RNA-dependent condensates *in vitro* at physiological conditions.

### The multimerization interface, but not the predicted IDRs, drives Nab2 condensation *in vitro*

Nab2 is predicted to contain several IDRs including an RGG and a polyQ domain (Fig. S4F). To identify the domains in Nab2 that are responsible for the alterations in nuclear dynamics in glucose stress, we tested the effect of IDR domain removal on condensation behavior *in vitro*. Deleting the Q-rich domain did not abolish condensate formation, but slightly shifted the pH-dependency to higher pH values (Fig. S5A). Deleting the RGG domain and replacing it with an exogenous nuclear import signal (ΔRGG^SV40NLS^, with the SV40NLS replacing the endogenous NLS found within the RGG domain) also affected the pH sensitivity of Nab2, which was even more pronounced than for Nab2^ΔQ^-GFP (Fig. S5A). Surprisingly, when both domains were deleted (Nab2^ΔQΔRGG-SV40NLS^-GFP, henceforth called ‘IDR mutant’), condensates could still form, even though the major regions expected to be critical for condensation were removed. In addition, droplets with this Nab2 variant formed even in the absence of RNA (Fig. 4A) and were less dynamic and more solid-like compared to their wild type counterparts, as shown by a reduction in recovery by FRAP (Fig. 4B). We then tested a Nab2 variant that has previously been shown to affect mRNA compaction by impairing Nab2 association with RNA (Nab2^F450A^, ^17^, henceforth called ‘interface mutant’). Nab2 forms an unusual hetero-tetramer, with Zn fingers 5-7 of two Nab2 molecules binding two different RNA strands^17^. The F450A mutation weakens this complex, possibly due to reduced valency. Intriguingly, the Nab2 interface mutant displayed a strong reduction in condensate formation (Fig. 4A). Thus, our results show that Nab2 condensate formation depends on the protein’s ability to interact with itself on RNA. Condensation does not require the IDRs but instead the major disordered regions within the protein, in particular the RGG domain, help to keep Nab2 condensates dynamic.

**4).**
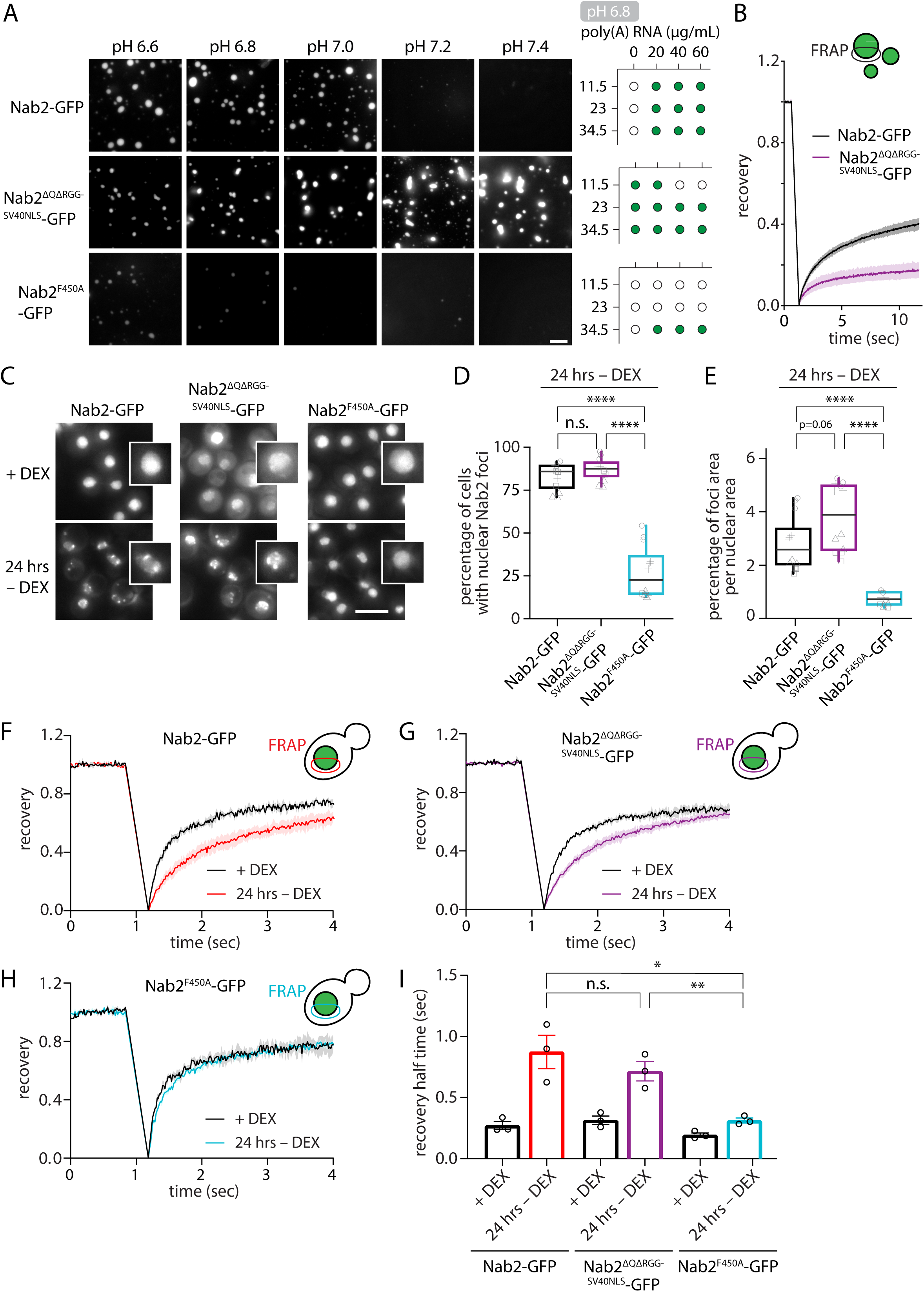
The Nab2 multimerization interface, but not the RGG domain function, drives condensation *in vitro*. A) GFP-tagged recombinant Nab2 wild type or Nab2 mutant droplets (23 µM each) were formed in LSB150 buffer to a final KCl concentration of 94 mM at the indicated pH in the presence of 0.04 mg/mL poly(A) RNA and imaged at 25 °C 30 min after setup. Right side: phase diagram of droplet formation at pH 6.8 with the indicated protein and RNA concentrations. Scale bar 10 µm. B) FRAP of 20 µM Nab2-GFP and Nab2^ΔQΔRGG-SV40NLS^-GFP droplets 30 min after setup with 0.04 mg/mL poly(A) RNA. FRAP curves are normalized to prebleach and postbleach values. Mean curves ± SEM of three biological replicates with ≥14 droplets per replicate are shown. C) Representative images of GFP-tagged Nab2 wild type and mutant cells in +DEX conditions and after acute shift and incubation for 24 hrs in –DEX. Scale bar 5 µm. D) Quantification of C). Percentage of cells with nuclear GFP foci. Shown are box plots of four biological replicates, each grey shape corresponds to individual technical replicates within a biological replicate, n>200 cells per replicate. **** p<0.0001, n.s. p=0.32, Wilcoxon test. E) Quantification of C). Percentage of foci area per nuclear area. Shown are box plots of four biological replicates, each grey shape corresponds to individual technical replicates within a biological replicate, n>200 cells per replicate. **** p<0.0001, Wilcoxon test. F-I) FRAP of cells shown in C). F) Nab2-GFP, G) Nab2^ΔQΔRGG-SV40NLS^-GFP, H) Nab2^F450A^-GFP. FRAP curves are normalized to prebleach and postbleach values. Mean curves ± SEM of three biological replicates with ≥15 cells per replicate are shown. I) Half time of recovery retrieved from fitting individual FRAP curves from F-H) with a single component model. Shown are means ± SEM of three biological replicates. * p=0.0155, ** p=0.0082, ns p=0.3694, unpaired t-test.

### Nab2 condensation properties influence the degree of nuclear mRNA retention

How does the ability of Nab2 to undergo condensation affect nuclear focus formation and RNA retention *in vivo*? Whereas the expression of a Nab2 variant that lacks the Q-rich domain (ΔQ) did not result in any measurable growth disadvantages, deletion of the RGG domain (ΔRGG^SV40NLS^) led to a severe growth impairment (Fig, S5B). Intriguingly, the combined deletion of both the Q and the RGG domain (ΔQ-ΔRGG^SV40NLS^, IDR mutant) rescued the growth defect of the ΔRGG^SV40NLS^ mutant (Fig. S5B), and we therefore continued our experiments with this mutant (Fig. S5C). Consistent with our *in vitro* results, the interface mutant Nab2^F450A^ formed almost no foci in the nucleus upon long-term glucose starvation (Fig. 4C-E) and also did not show a reduction in nuclear mobility (Fig. 4H,I). In contrast, both Nab2^ΔQ^-GFP and Nab2^ΔQΔRGG-SV40NLS^-GFP still formed nuclear foci (Fig. 4C-E and Fig. S5E) coinciding with a reduction in nuclear mobility compared to wild type Nab2-GFP in FRAP (Fig. 4F,G,I and Fig. S5F,G).

The pool of Nab2 within nuclear foci displayed reduced recovery compared to the remaining nuclear Nab2 pool outside of foci (Fig. S5H), indicating that there is a larger immobile fraction within foci. In contrast to the *in vitro* FRAP experiment (Fig. 4B), where the IDR mutant showed a strong reduction in mobility within the droplet compared to wild type Nab2, the *in vivo* recovery kinetics of the Nab2 pool within nuclear foci did not differ between wild type and IDR mutant. This suggests that additional factors can modulate Nab2 mobility *in vivo*, potentially to prevent detrimental solidification. However, compared to wild type the IDR mutant still formed overall larger nuclear foci, while the interface mutant formed fewer and smaller nuclear foci (Fig. 4C-E), suggesting that the degree to which Nab2 mutants form condensates *in vitro* reflects focus formation *in vivo*.

Intriguingly, Nab2’s ability to form foci in stress is also linked to its ability to retain RNA in the nucleus: while the IDR mutant showed an increased nuclear poly(A) RNA signal and a decreased cytoplasm-to-nucleus (C/N) RNA ratio, the interface mutant displayed a decreased nuclear poly(A) RNA signal and an increased C/N RNA ratio upon glucose withdrawal (Fig. 5A-C). This implies that the level of Nab2 condensate formation correlates with the degree of nuclear RNA retention in long-term glucose starvation. Of note, the Nab2 IDR mutant also showed a slightly altered steady-state mRNA localization even in log growing cells (Fig. 5A and S5I,J), suggesting that the RGG domain is important for efficient mRNA export both in unstressed and stressed cells.

**5).**
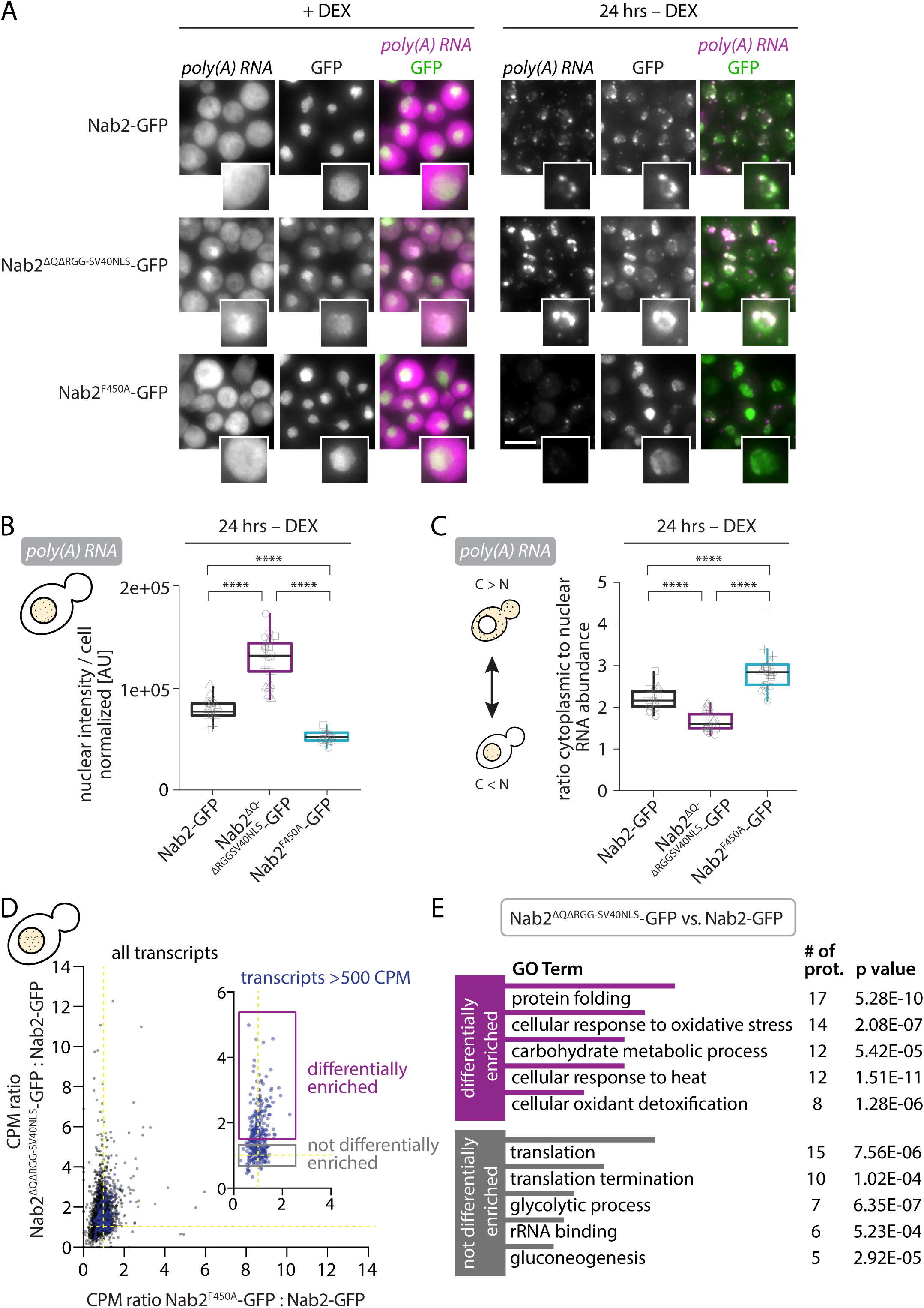
The degree of Nab2 condensation determines the level of nuclear mRNA retention. A) RNA FISH against poly(A) RNA using an oligo(dT)30 probe was performed in fixed cells expressing Nab2-GFP, Nab2^ΔQΔRGG-SV40NLS^-GFP or Nab2^F450A^-GFP. Samples were taken in +DEX or after acute starvation in medium without DEX for 24 hrs. Insets show single nucleus (2.16x magnification). Scale bar 5 µm. B) Quantification of nuclear poly(A) RNA signal intensity in 24 hrs –DEX condition in A). Data is normalized to median RNA value of all replicates. Shown are box plots of four biological replicates (each grey shape corresponds to individual technical replicates within a biological replicate, n>100 cells per replicate). **** p<0.0001, Wilcoxon test. C) Quantification of cytoplasmic to nuclear poly(A) RNA intensity ratios per cell in 24 hrs –DEX condition in A). Data is normalized to median RNA value of all replicates. Shown are box plots of four biological replicates (each grey shape corresponds to individual technical replicates within a biological replicate, n>100 cells per replicate). **** p<0.0001, Wilcoxon test. D) Plot of ratios of nuclear RNA sequencing reads (CPM, counts per million). Single dots represent individual genes. Yellow line indicates CPM ratio of 1:1. Median CPM ratio Nab2^ΔQΔRGG-SV40NLS^- GFP:Nab2-GFP 1.2668; median CPM ratio Nab2^F450A^-GFP:Nab2-GFP 0.899. Inset shows most abundant reads (blue dots, >500 CPM in Nab2-GFP average of 3 biological replicates, representing 42% of the total reads). Boxes highlight differentially enriched genes (Nab2^ΔQΔRGG-SV40NLS^-GFP:Nab2-GFP; purple, fold change ratio >1.5) and not differentially enriched genes (Nab2^ΔQΔRGG-SV40NLS^-GFP:Nab2-GFP; grey, fold change ratio between 0.8 and 1.2). E) Gene ontology (GO) term analysis of most abundant reads (blue dots) from C) using DAVID Functional Annotation Clustering tool ^82,83^.

### In glucose stress Nab2 condensation leads to RNA retention in an RNA dose-dependent manner and affects survival after prolonged stress

In order to identify which mRNAs are retained in nuclear Nab2 foci in stress and to examine if this is altered in the Nab2 mutant backgrounds (Nab2^ΔQΔRGG-SV40NLS^-GFP and Nab2^F450A^-GFP) we purified poly(A) RNA from whole cells cultured in +glucose and 24 hrs –glucose conditions, as well as from nuclear fractions of cells starved for 24 hrs in –glucose (Fig. S5K,L and Supplementary Table 4). As expected, we detected a global rearrangement of the transcriptional landscape between the +glucose and 24 hrs –glucose conditions on a whole cell level (Fig. S6A). This is in agreement with RNA FISH quantifications that revealed a 4-5-fold reduction of total poly(A) RNA abundance in 24 hrs –glucose compared to +glucose (Fig. S5J). However, for a given condition, there was good correlation in RNA abundance across the wild type and different mutants (Fig. S6A) and a stress response was initiated to a similar extent in all strains at the total RNA level (Fig. S6B). By contrast, differences in the nuclear RNA fractions could be observed in 24 hrs –glucose revealing an increase in sequencing reads (CPM, counts per million) for the IDR mutant Nab2^ΔQΔRGG-SV40NLS^ compared to wild type (Fig. 5D and S6C,D, median CPM ratio 1.2668). The IDR mutant displayed nuclear enrichment of certain transcripts whereas others did not show any differential enrichment (Fig. 5D). Gene ontology (GO) term analysis of the most abundant transcripts (CPM >500 in Nab2-GFP average of 3 replicates (318 genes) representing 42% of the total reads; Fig. 5D (blue dots), Fig. 5E (purple) and Fig. S6F,G) revealed that the highest-scoring GO terms for differentially enriched transcripts were stress response-related (protein folding, cellular response to oxidative stress/heat, cellular oxidant detoxification), and carbohydrate metabolic processes. Transcripts that were not differentially enriched (Fig. 5E, grey) showed highest-scoring GO terms in translation, translation termination, glycolytic process/gluconeogenesis and rRNA binding (i.e., ‘pro-proliferation’ mRNAs).

In contrast to the poly(A) *in situ* hybridization experiments, we could not observe a difference in sequencing reads between Nab2 wild type and the interface mutant Nab2^F450A^ (median CPM ratio Nab2^F450A^-GFP:Nab2-GFP 0.899). To investigate this discrepancy, we performed single molecule *in situ* hybridization (smFISH) experiments (Fig. 6A) using probes against individual mRNAs that either belong to the class of differentially enriched mRNAs (*BTN2, HSP42, HXT6*, Fig. 6B-D, Fig. 7A-D and Fig. S6H) or to the not differentially enriched mRNAs (*DED1*, Fig. 6E, Fig. 7A-D and Fig. S6H). The IDR mutant displayed increased nuclear RNA signal, a higher percentage of cells with nuclear RNA accumulation and a stronger degree of Nab2:mRNA co-localization for differentially enriched mRNAs (*BTN2, HSP42, HXT6*), but not for not differentially enriched mRNAs (*DED1*). Conversely, the interface mutant showed decreased nuclear RNA signal, a lower percentage of cells with nuclear mRNA accumulation and a weaker degree of Nab2:mRNA co-localization for *BTN2* and *HSP42*, while mRNA enrichment was similar between wild type and interface mutant for *HXT6*, which could be due to low nuclear mRNA abundance, higher cell-to-cell variability or additional regulatory effects during starvation. Interestingly, we observed differences in the C/N mRNA ratio between wild type and the Nab2 mutants for all four tested mRNAs. The mRNAs displayed a lower C/N ratio in the IDR mutant compared to wild type (Fig. 7A) suggesting a higher degree of mRNA retention in this variant. By contrast, the interface mutant showed a higher C/N mRNA ratio implying that less Nab2 condensation results in mRNA leakage from the nucleus (Fig 7B). The discrepancy between the RNA sequencing and the smFISH results could be due to a lower sensitivity of bulk RNA sequencing compared to single cell single molecule RNA visualization, making it difficult to detect small nuclear abundance differences for low-expressed mRNAs such as *DED1* using RNAseq (Fig.7A). Additionally, as extraction of intact yeast nuclei is inherently difficult^53^, leftover traces of cytoplasm or whole cell mRNA (Fig. S5L) could have affected the sequencing results of the nuclear RNA extracts. Furthermore, the poly(A) selection step in the RNA sequencing protocol might add an additional bias towards capturing mRNAs with long(er) poly(A) tails.

**6).**
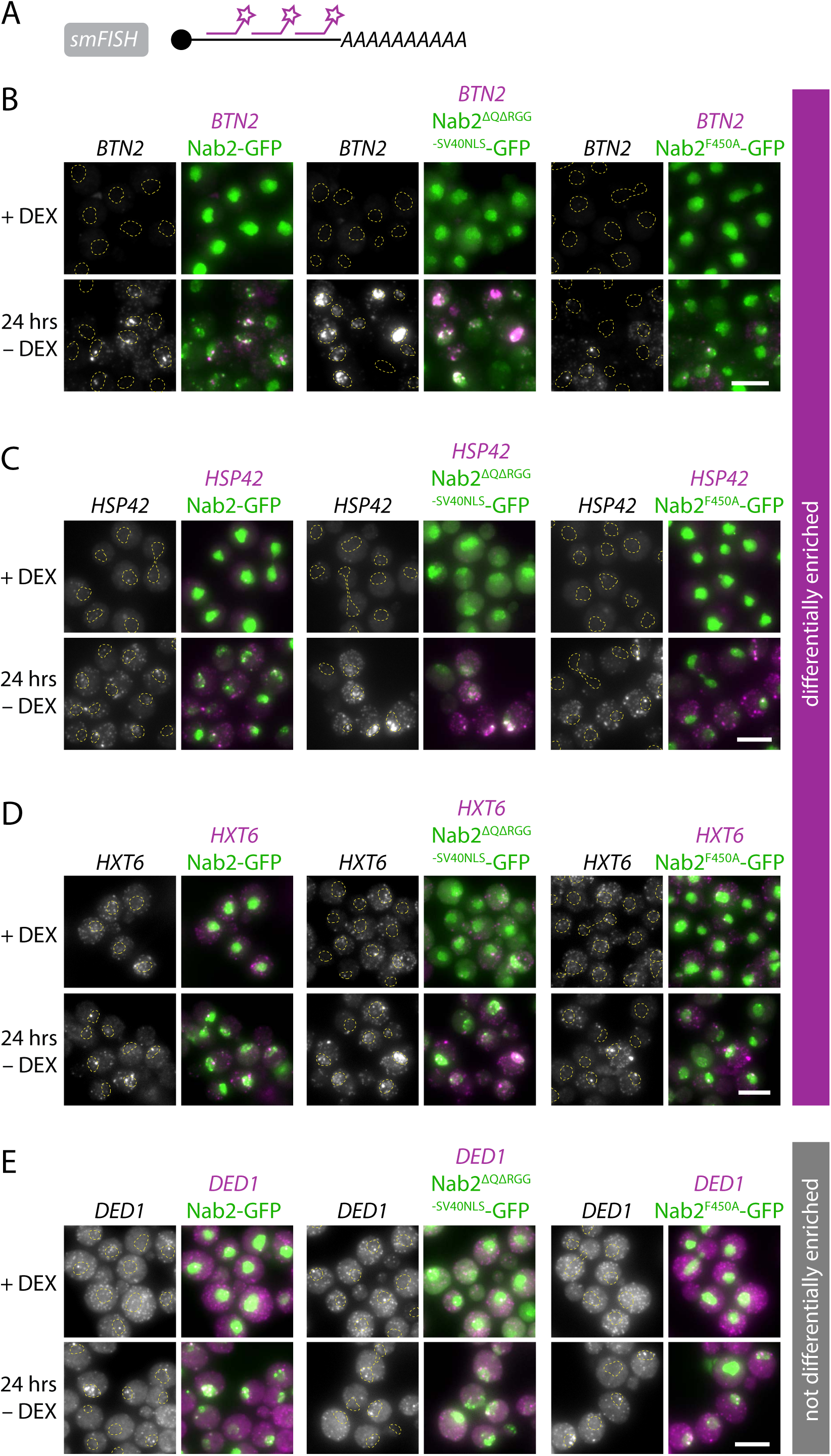
Nab2 condensation mutants affect nuclear mRNA retention *in vivo* during stress. A) Representative images of RNA FISH against *BTN2*, which was performed in fixed cells expressing Nab2-GFP, Nab2^ΔQΔRGG-SV40NLS^-GFP or Nab2^F450A^-GFP. Samples were harvested in +DEX or after acute starvation in medium without DEX for 24 hrs. Dotted yellow line indicates nuclear outline. Scale bar 5 µm. B) Representative images of RNA FISH against *HSP42*, which was performed in fixed cells expressing Nab2-GFP, Nab2^ΔQΔRGG-SV40NLS^-GFP or Nab2^F450A^-GFP. Samples were harvested in +DEX or after acute starvation in medium without DEX for 24 hrs. Dotted yellow line indicates nuclear outline. Scale bar 5 µm. C) Representative images of RNA FISH against *HXT6*, which was performed in fixed cells expressing Nab2-GFP, Nab2^ΔQΔRGG-SV40NLS^-GFP or Nab2^F450A^-GFP. Samples were harvested in +DEX or after acute starvation in medium without DEX for 24 hrs. Dotted yellow line indicates nuclear outline. Scale bar 5 µm. D) Representative images of RNA FISH against *DED1*, which was performed in fixed cells expressing Nab2-GFP, Nab2^ΔQΔRGG-SV40NLS^-GFP or Nab2^F450A^-GFP. Samples were harvested in +DEX or after acute starvation in medium without DEX for 24 hrs. Dotted yellow line indicates nuclear outline. Scale bar 5 µm.

**7).**
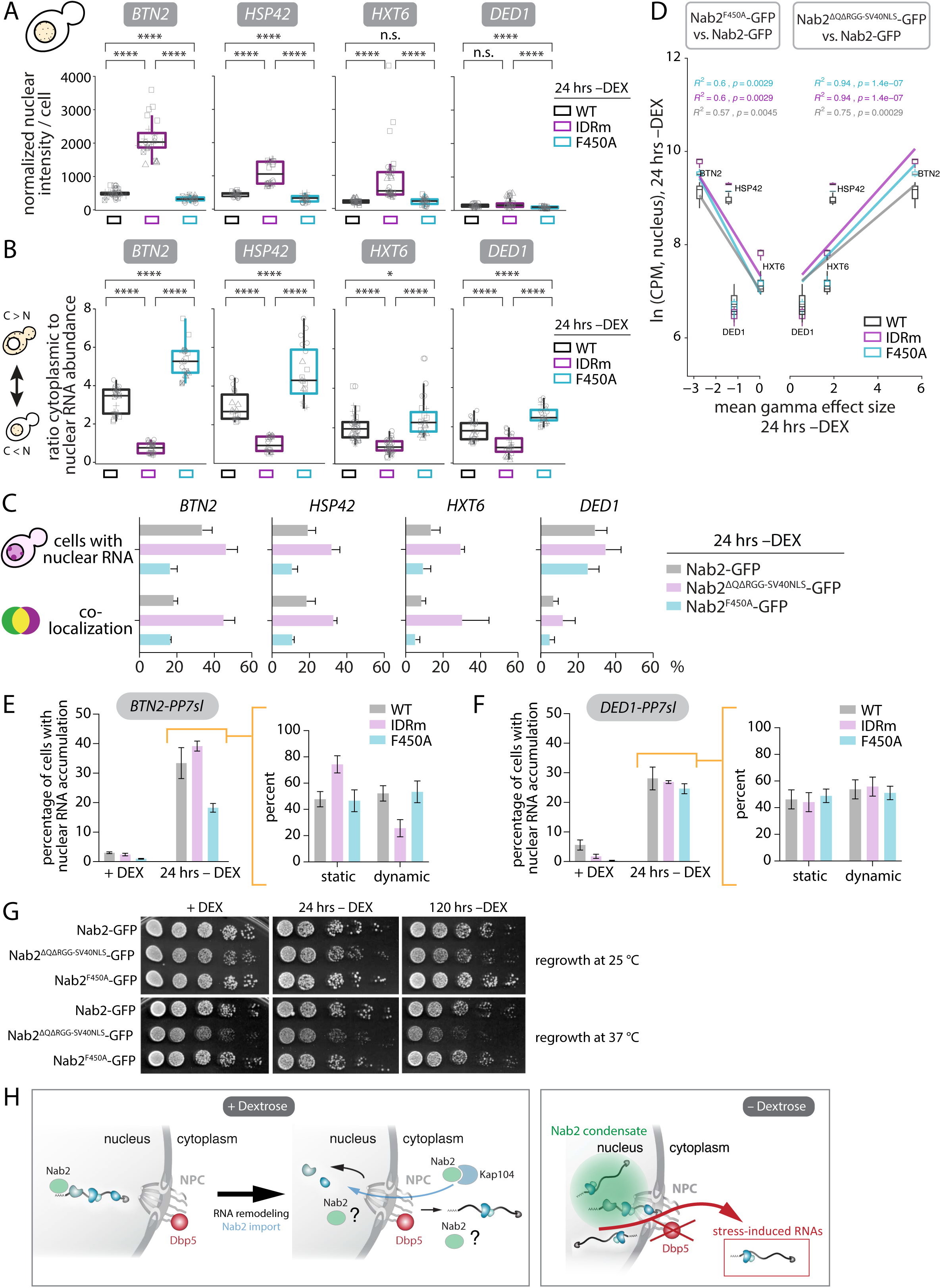
Nab2 condensation confers dose-dependent mRNA retention *in vivo* and correct Nab2 condensation is essential for survival after prolonged stress. A) Quantification of nuclear RNA signal intensity in 24 hrs –DEX condition as shown in Figure 6A-D. Data is normalized to median RNA value of all replicates. Shown are box plots of four (*BTN2*: five; *HXT6*: six) biological replicates, **** p<0.0001, n.s. (*HXT6*) p=0.95, n.s.(*DED1*) p=0.49, Wilcoxon test. B) Quantification of cytoplasmic to nuclear poly(A) RNA intensity ratios per cell as shown in Figure 6B-E). Shown are box plots of four (*BTN2*: five; *HXT6*: six) biological replicates, each grey shape corresponds to individual technical replicates within a biological replicate, n>100 cells per replicate. **** p<0.0001, *(*HXT6*) p=0.038, Wilcoxon test. C) Quantification of 24 hrs –DEX condition in Fig. 6B-E. (Top) percentage of cells with nuclear RNA; (Bottom) co-localization, percentage of nuclear GFP foci co-localizing with indicated RNA (quantified only in cells with nuclear RNA signal). Shown are means ± SEM of four biological replicates, n>100 cells per replicate. D) Correlation between extent of nuclear accumulation ((left) mean gamma effect size Nab2^F450A^- GFP vs. Nab2-GFP, (right) mean gamma effect size Nab2^ΔQΔRGG-SV40NLS^-GFP vs. Nab2-GFP; calculated from data in Figure 7A using nonparametric Cohen’s d-consistent effect size) and nuclear mRNA abundance using RNAseq reads from nuclear extracts shown in Figure 5D (natural logarithm of CPM data), R^2^ and p values calculated by Spearman correlation. E) Live imaging of PCP-mCherry-bound PP7sl-tagged *BTN2* mRNAs in cells expressing Nab2-GFP, Nab2^ΔQΔRGG-SV40NLS^-GFP or Nab2^F450A^-GFP in +DEX and after acute shift and incubation for 24 hrs in –DEX. (Left) nuclear RNA accumulation, percentage of cells where average RNA signal intensity is larger in nucleus than in cytoplasm; (Right) Mobility of nuclear mRNAs in 24 hrs –DEX condition. Shown are means ± SEM of three biological replicates, n>100 cells per replicate. F) Live imaging of PCP-mCherry-bound PP7sl-tagged *DED1* mRNA in cells expressing Nab2-GFP, Nab2^ΔQΔRGG-SV40NLS^-GFP or Nab2^F450A^-GFP in +DEX and after acute shift and incubation for 24 hrs in –DEX. (Left) nuclear RNA accumulation, percentage of cells where average RNA signal intensity is higher in nucleus than in cytoplasm; (Right) Mobility of nuclear mRNAs in 24 hrs –DEX condition. Shown are means ± SEM of three biological replicates, n>100 cells per replicate. G) Growth recovery assay of cells expressing Nab2-GFP, Nab2^ΔQΔRGG-SV40NLS^-GFP or Nab2^F450A^- GFP. Mid-log cultures grown in +DEX or acutely starved in medium without DEX for 24 hrs or 120 hrs were recovered as 1:5 serial dilutions (starting OD_600_ 0.2) on rich medium plates containing glucose at either 25 °C for 2 days or 37 °C for 1 day. H) Model depicting successful export of mRNAs from the nucleus in +DEX conditions (left), where Dbp5 aids in the removal of RNA-binding proteins from the translocating transcript. It is unclear at which point Nab2 is removed from the mRNA, but Nab2 gets re-imported into the nucleus through binding to the karyopherin Kap104. In 24 hrs –DEX (right), nuclear Nab2 condensation leads to bulk mRNA retention in the nucleus and allows transport of stress-induced mRNAs.

The smFISH results also indicated a dose-dependent effect of the Nab2 condensate on nuclear mRNA retention. When we calculated the mean gamma effect size, which reflects the strength of the mutants to increase or decrease nuclear mRNA retention compared to wild type, we observed a dose-dependent mRNA retention effect (Fig. 7D). The more abundant the mRNA in the nucleus (based on the RNAseq data from the nuclear extracts in Fig. 5D), the stronger the nuclear mRNA abundance increase in the IDR mutant (Fig. 7D, right), and the greater the loss of nuclear mRNA signal in the interface mutant (Fig. 7D, left).

Since RNA FISH experiments are inherently static, we also visualized PP7sl-labeled mRNAs in live cells. High-speed single RNA tracking was not possible due to the dim PCP-mCherry label as well as our inability to simultaneously capture single mRNAs and multi-copy mRNA clusters with the highly sensitive EM-CCD camera. We therefore switched to low-speed, low-light long-term imaging to capture general differences in mRNA localization and mobility in early (up to 30 min, Fig. S7A,B) and late –DEX starvation (24 hrs, Fig. 7E,F). We could recapitulate the extent of nuclear RNA accumulation in the wild type and different mutants seen with smFISH (Fig. 7C). After 24 hrs in –DEX, wild type Nab2 cells displayed a roughly equal distribution of static (Video S4, Fig. 7E,F) or dynamic nuclear mRNAs (Video S5, Fig. 7E,F). The interface mutant Nab2^F450A^ differed from wild type Nab2 only in the extent of nuclear *BTN2-PP7sl* accumulation, most likely due to fewer nuclear Nab2 foci (Fig. 4D,E), but not the mobility of remaining nuclear RNAs (Fig. 7E). The IDR mutant Nab2^ΔQΔRGG-SV40NLS^ not only accumulated more *BTN2-PP7sl* mRNAs but these RNAs were also more static compared to wild type (Fig. 7E). *DED1-PP7sl* mRNAs in the different Nab2 backgrounds did not show any difference in the amount of nuclear mRNA accumulation or in mRNA mobility (Fig. 7F). For early time points of –DEX starvation, the difference between wild type Nab2 and the interface mutant Nab2^F450A^ became even more apparent, as most of the PP7sl-labeled *BTN2* and *DED1* mRNAs did not show any nuclear accumulation in the interface mutant (Fig. S7A,B).

In summary, these experiments suggest that nuclear mRNA retention in glucose starvation is affected by Nab2 condensation. Our results imply that the altered condensate state seen in the IDR mutant Nab2^ΔQΔRGG-SV40NLS^ impairs the transport of stress-induced mRNAs, which results in their nuclear accumulation. Conversely, the interface mutant Nab2^F450A^ weakens nuclear Nab2 condensation which leads to a global nuclear mRNA retention defect and results in increased cytoplasmic mRNA localization.

Finally, we wanted to test whether there is a physiological consequence of altered mRNA retention in long-term stress. We therefore washed cells acutely into medium without glucose and incubated them for 24 hrs or 120 hrs before allowing them to recover on rich medium plates containing glucose. Compared to wild type, the IDR mutant Nab2^ΔQΔRGGSV40NLS^ displayed a slight growth disadvantage on rich medium plates in log growing cells, which was exacerbated after prolonged glucose withdrawal (Fig. 7G and S7E,F) and when stress is maintained after dextrose recovery by incubating cells at different temperatures (Fig. 7G and S7C). Furthermore, gradual glucose depletion also affected the recovery of the IDR mutant as cells that had recovered from early stationary phase (24 hrs after mid-log phase) showed a similar growth impairment as after acute glucose withdrawal (Fig. S7D). The slow growth of the IDR mutant is caused by a delayed recovery after stress resulting in a prolonged lag phase, as all strains display almost identical growth kinetics during exponential growth (Fig. S7G).

Together, these results suggest that aberrant mRNA localization caused by altered Nab2 condensation can affect recovery from glucose stress conditions.

## Discussion

Here we show that the mRNA-binding factor Nab2 can form RNA-dependent condensate-like foci *in vitro* and *in vivo*, and that nuclear focus formation *in vivo* is counteracted by active mRNA export mediated by the DEAD-box ATPase Dbp5. Our results also provide evidence that global retention of bulk mRNAs in the nucleus in physiologically relevant stress conditions coincides with nuclear RNA-dependent Nab2 condensation. Furthermore, our data suggest that the degree of Nab2 condensation influences the degree of mRNA retention, and this in turn is affected by the nuclear abundance of a given mRNA (Supplementary Table 5). Based on these results, we propose the following model (Fig. 7H): in +DEX conditions, mRNAs traverse the NPC and get remodeled at the NPC by Dbp5 for successful nuclear exit. It is still unclear at which step and in which compartment Nab2 is being removed from the mRNP. Unlike several other nuclear mRNP processing factors, it shuttles between nucleus and cytoplasm^21^, and a study has shown a transient interaction between Nab2 and Dpb5 at the cytoplasmic side of the NPC^28^. Therefore, Nab2 release from the mRNA might occur when the mRNA has already partially or completely transited to the cytoplasm. Upon –DEX starvation, reduced abundance of Dbp5 at the NPC leads to a global mRNA export block, probably by inhibiting mRNP remodeling and removal of factors such as Nab2 from export-competent mRNPs by Dbp5. Over time, as the export block persists mRNPs accumulate in the nucleus in distinct foci that contain Nab2 and the cap-binding complex, suggesting that the mRNA cap and poly(A) tail are still present (Fig. 3C,D and Fig. S4A), possibly to protect the mRNAs from nuclear degradation (as has been shown for Nab2-bound transcripts in acute Mex67 depletion^54^) and/or to store them for later use either during prolonged stress or after stress release. However, stress-related transcripts still need to escape nuclear retention to elicit a timely cellular stress response, suggesting that cells could use confinement in or release from nuclear condensates to re-wire mRNA export during stress. Storing transcripts in the nucleus during glucose starvation might be a strategy for rapidly re-initiating cell growth after stress release, allowing cells to quickly re-start protein production in a transcription-independent fashion and minimize energy consumption at early stages of re-growth.

Although our data do not rule out selectivity towards certain mRNA classes, Nab2 condensate retention potential seems to scale with nuclear mRNA abundance (Fig. 7D). The more abundant a given mRNA in the nucleus, the more it gets retained in the IDR mutant with increased Nab2 condensation, and the more it is lost from the nucleus in the interface mutant with weakened nuclear Nab2 condensation. All four tested mRNAs from either the differentially enriched class of mRNAs or the not differentially enriched mRNAs (based on the RNAseq classification, Fig. 5D,E) displayed altered steady-state localization in both Nab2 mutants (Fig. 6B-E and Fig. 7A,B). This highlights differences in sensitivity between bulk RNA sequencing and single molecule RNA FISH and emphasizes the necessity to confirm RNA sequencing results with orthogonal approaches. RNA dosage dependency could be caused by differences in mRNA induction strength in stress, suggesting that uninduced mRNAs such as *DED1* are less affected by the IDR mutant as they are not being actively transcribed in stress conditions, and are perhaps therefore more readily lost from the nucleus in the interface mutant. Alternatively or additionally, differences in the primary sequence or secondary structure of the mRNA could lead to the observed phenotypes. Indeed, we see differences in mRNA length (Fig. S6I) and the amount of mRNAs stemming from intron-containing genes (Fig. S6J) between the differentially enriched and the not differentially enriched groups of transcripts, although it is unclear if those mRNAs still contain introns in nuclear Nab2 condensates. Nab2 has been implicated in splicing regulation and artificial depletion of Nab2 from the nucleus leads to transcriptional read-through which produces chimeric transcripts with retained introns^18,55^. RNA length has been reported to correlate with enrichment in cytoplasmic stress granules and poor translation efficiency^56^, suggesting that short transcripts, such as mRNAs responsible for the cellular stress response, could evade both nuclear and cytoplasmic condensates to allow for their preferential translation in stress conditions. Furthermore, nuclear mRNA retention has previously been reported in mammalian systems, where it serves to rapidly modify the neuronal transcriptome upon stimulation^57^, or to buffer gene expression noise^58^. Buffering mechanisms could also be at play in yeast, which needs to be investigated in the future.

Our model does not rule out the possibility that other factors, such as the major export adaptor Mex67-Mtr2, are involved in regulating Nab2 condensation, or influence nuclear mRNA retention in a Nab2-independent manner. Indeed, Tudek and colleagues have reported nuclear RNA focus formation following acute anchor-away of both Mex67 and Mtr2, which like acute Dbp5 depletion also causes a global mRNA export block^54^. However, the authors did not link RNA focus formation to Nab2’s role in protecting RNAs from nuclear degradation. We show that Mex67 localization is unaffected in 24 hrs – DEX starvation (Fig. S3C) and we have previously reported that acute Dbp5 depletion affects neither Mex67’s localization to nor its mobility at the NPC^22^. We therefore suspect that Mex67-Mtr2 possibly indirectly affects Nab2 condensation by promoting the export of a Nab2-bound mRNP.

It has previously been reported that treatment with the glucose analog 2-deoxyglucose in combination with the respiratory chain inhibitor antimycin A, which causes a global shutdown of the cell’s energy production, restricts the NPC diameter of fission yeast *S. pombe* and leads to an mRNA export block^59^. Therefore, 24 hrs glucose withdrawal used in our experiments could lead to physical changes within the NPC structure, which could restrict mRNA export, which in turn could lead to the nuclear accumulation of export-competent mRNPs in Nab2-positive nuclear condensates. However, it remains to be determined if the NPC diameter and/or its biophysical properties are indeed affected in physiological stress conditions. Our data reveal that Nab2 affects nuclear RNA retention in glucose stress as modifying the state of the Nab2 condensate results in altered nuclear RNA retention (Fig. 5A-C), but it remains to be determined which additional mechanisms modulate mRNA transport in stress.

The other tested mRNA-binding proteins, Yra1 and Npl3, do not enrich in Nab2 condensates (Fig. S2H,I), and this selective exclusion could hint towards nuclear remodeling of mRNPs in stress conditions, which could in turn facilitate bulk nuclear mRNA storage. Since their nuclear mobility does not change in unstressed versus stressed cells (Fig. S2J-M), Yra1 and Npl3 could still dynamically associate with a subset of mRNAs that are excluded from the Nab2 condensate in stress, e.g., stress-induced mRNAs, which could be a requirement for their swift and efficient nuclear export. Future research is needed to understand mRNP composition and its functional consequence in long-term glucose stress.

We show that forcing Dbp5 to NPCs via fusion to ^ΔN^Nup159 prevents stress-induced Nab2 mobility reduction and nuclear focus formation (Fig. 2I-K). Previous reports have demonstrated a transient interaction between Nab2 and Dbp5 at the NPC using a splitVenus system^28^, suggesting that Nab2 could be removed from the mRNP by Dbp5 at late stages of NPC transit. We speculate that the DEAD-box ATPase Dbp5 could generally act on RNPs to remodel them or to dissolve RNP-containing condensates. Consistent with this, overexpression of the mammalian ortholog of Dbp5, DDX19, prevents stress granule formation in an RNA binding-dependent manner ^60,61^. However, local ATPase activation through its interaction partner Gle1, which resides exclusively at the NPC, could target Dbp5’s activity largely towards specific RNPs such as Nab2-bound mRNAs that pass through the NPC during export. In line with this, Dbp5-^ΔN^Nup159 localization has been shown to be sufficient for viability^28^, suggesting that the essential role of Dbp5 is restricted to the NPC.

Nuclear Nab2 condensation depends on several domains that modulate the degree of condensation and the material properties of the condensates, thereby influencing the degree of nuclear RNA retention (Fig. 5A-C). Surprisingly, when we deleted the major regions predicted to be causal for condensation (Fig. S4F), the Q-rich domain and the RGG domain, we observed a stronger Nab2 condensation phenotype, indicating that these domains act together to function as ‘liquefiers’ of condensates or to block condensation altogether. Similar observations have been made for the stress granule component Ded1, where deletion of evolutionarily preserved residues in its N-terminal intrinsically disordered region (IDR) lead to constitutive SG formation^62^. How these two domains act together to facilitate proper Nab2 condensation is unclear. Their relative position within the protein structure might be important, and the presence of an RGG domain could provide a survival benefit when the protein also contains a Q-rich domain, as deletion of the Nab2 RGG domain alone is lethal, while co-deleting the Q-rich domain restores growth (Fig. S5B). Spatial domain organization has previously been shown to affect condensate formation, as placing a Q-rich region close to the RRM of G3BP1 restored SG formation^63^.

Conversely, condensation of Nab2 (and nuclear focus formation/RNA retention) depends on its ability to form a hetero-tetramer with RNA (Nab2^F450A^, interface mutant, Fig. 4A,C-E and 5A). Reduced valency, due to weakened RNA binding^17^, could therefore lead to reduced condensation. The interface mutation lies within the zinc fingers (ZnF) essential for RNA binding, highlighting the multivalent binding mode of Nab2 as well as the importance of non-IDR regions for condensate formation. Indeed, it was previously shown that increasing the number of ZnFs promotes cytoplasmic condensation of the SG component G3BP1, highlighting how altering RNA binding strength and potentially the valency of interaction affects condensate formation^63^.

Glucose starvation is accompanied by an intracellular pH drop from pH 7.5 to approximately pH 6-6.5 ^64–67^. Intriguingly, altered condensation of the IDR mutant Nab2^ΔQΔRGG-SV40NLS^ resulted in pH insensitivity *in vitro* (Fig. 4A) and an altered steady-state localization of poly(A) RNA even in +glucose conditions (Fig. 5A and Fig. S5I), suggesting that Nab2 condensation could also play a role in mRNA export in exponentially growing cultures. Future research is needed to determine the extent to which mRNP (micro)condensation is also involved in mRNA export in unstressed cells, and if the essential function Dbp5 in mRNA export is to locally prevent or modulate mRNP condensation.

### Limitations of the study

We are only beginning to understand the formation and regulation of biomolecular condensates. This study shows that Nab2 forms RNA-dependent condensates *in vitro* and *in vivo*, yet the precise mechanism by which RNA retention is modulated by Nab2 condensation during long-term glucose starvation, and the upstream signaling cues that drive Nab2 condensation, still need to be elucidated.

We also lack an understanding of RNA sequence composition and turnover in long-term glucose starvation, and our RNA FISH experiments might have missed changes in poly(A) tail length and/or RNA decay intermediates. This could lead to altered FISH labeling, which in turn would introduce a bias in our readout for RNA localization. The alternative approach of PP7sl labeling confirmed our RNA FISH results but also has the inherent problem of adding substantial mass to the endogenous transcript, which could lead to aberrant RNA localization and decay^68,69^. Furthermore, microscopy limitations did not allow us to follow individual RNAs at high temporal resolution to investigate the purpose of nuclear retention and the fate of the nuclear retained RNAs during or after stress.

A general bottleneck in the field is a lack of tools with which to study biomolecular condensation in cells. In our FRAP experiments, for example, limited spatial resolution and the small size of the Nab2 foci prevented us from performing half-structure bleaching experiments in glucose starvation, which would have been informative of recovery dynamics within the condensate.

Lastly, still very little is known about the regulation of mRNA retention and export in different organisms. While yeast Nab2 is conserved and its human homolog, ZC3H14, has been shown to localize to nuclear speckles^70^, it is unclear if nuclear condensation coupled with selective RNA retention are also features of other biological systems.

## Materials and Methods

### Lead contact

Further information and requests for resources and reagents should be directed to the lead contact, Karsten Weis.

**Supplemental Table 1.**
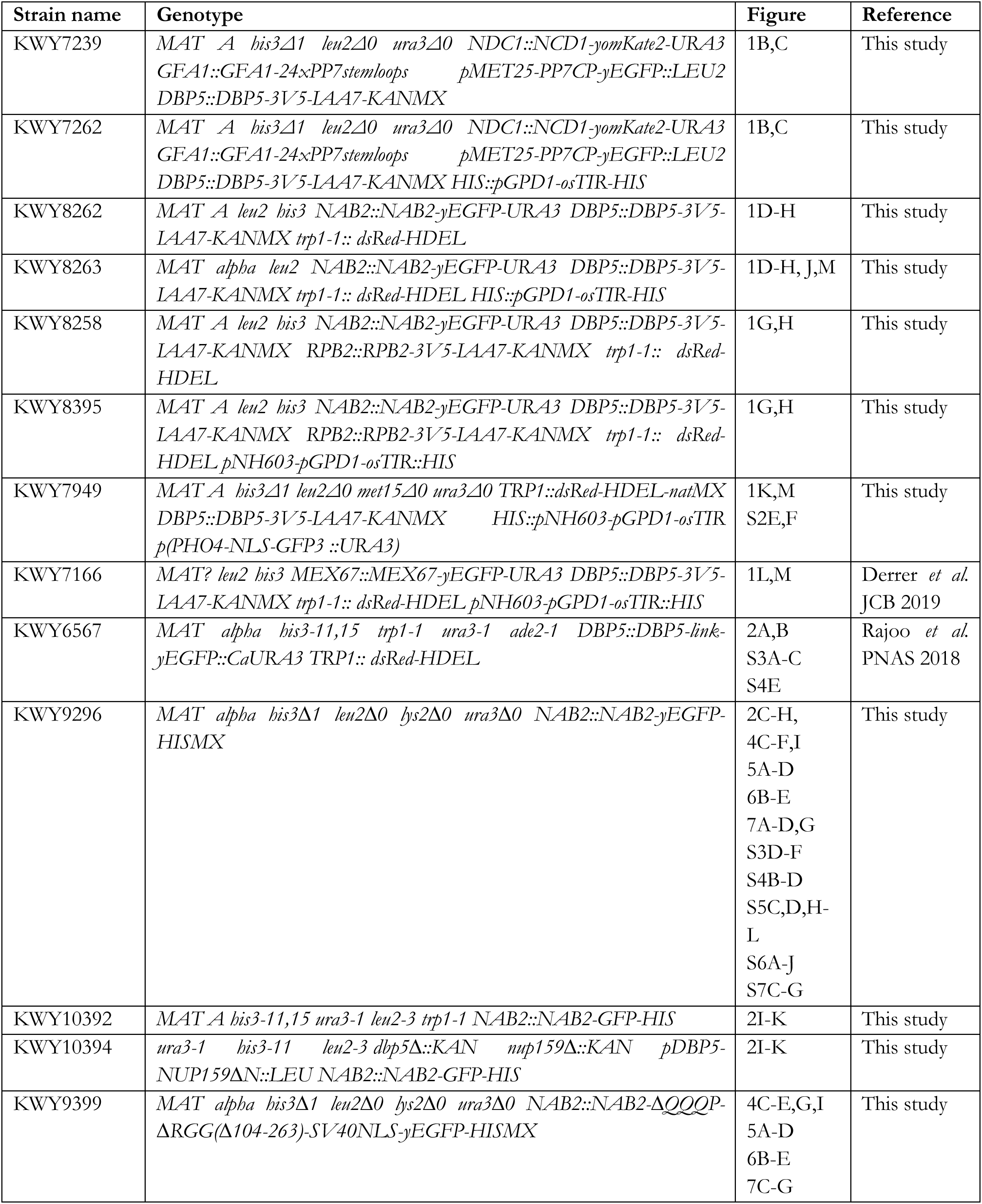
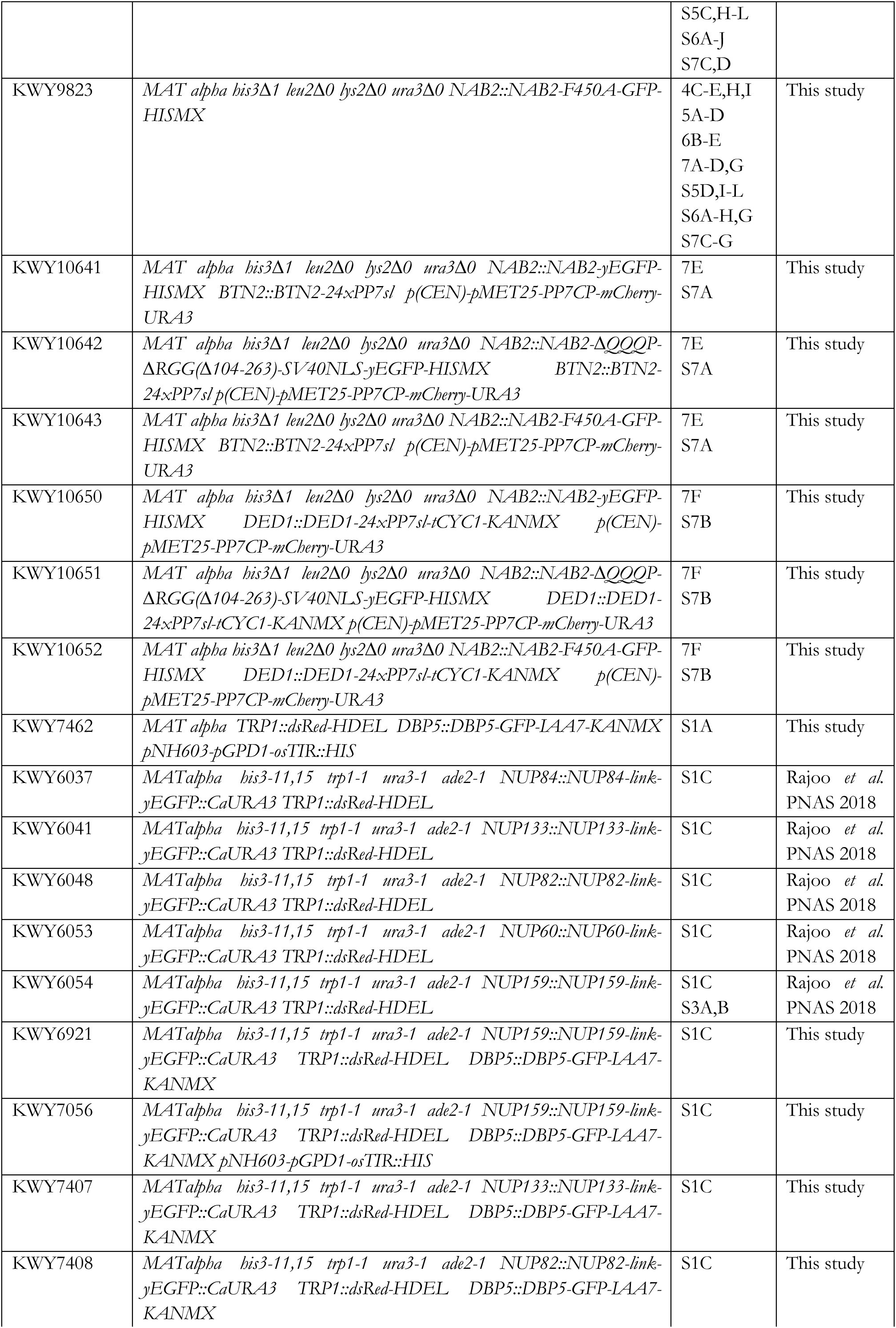
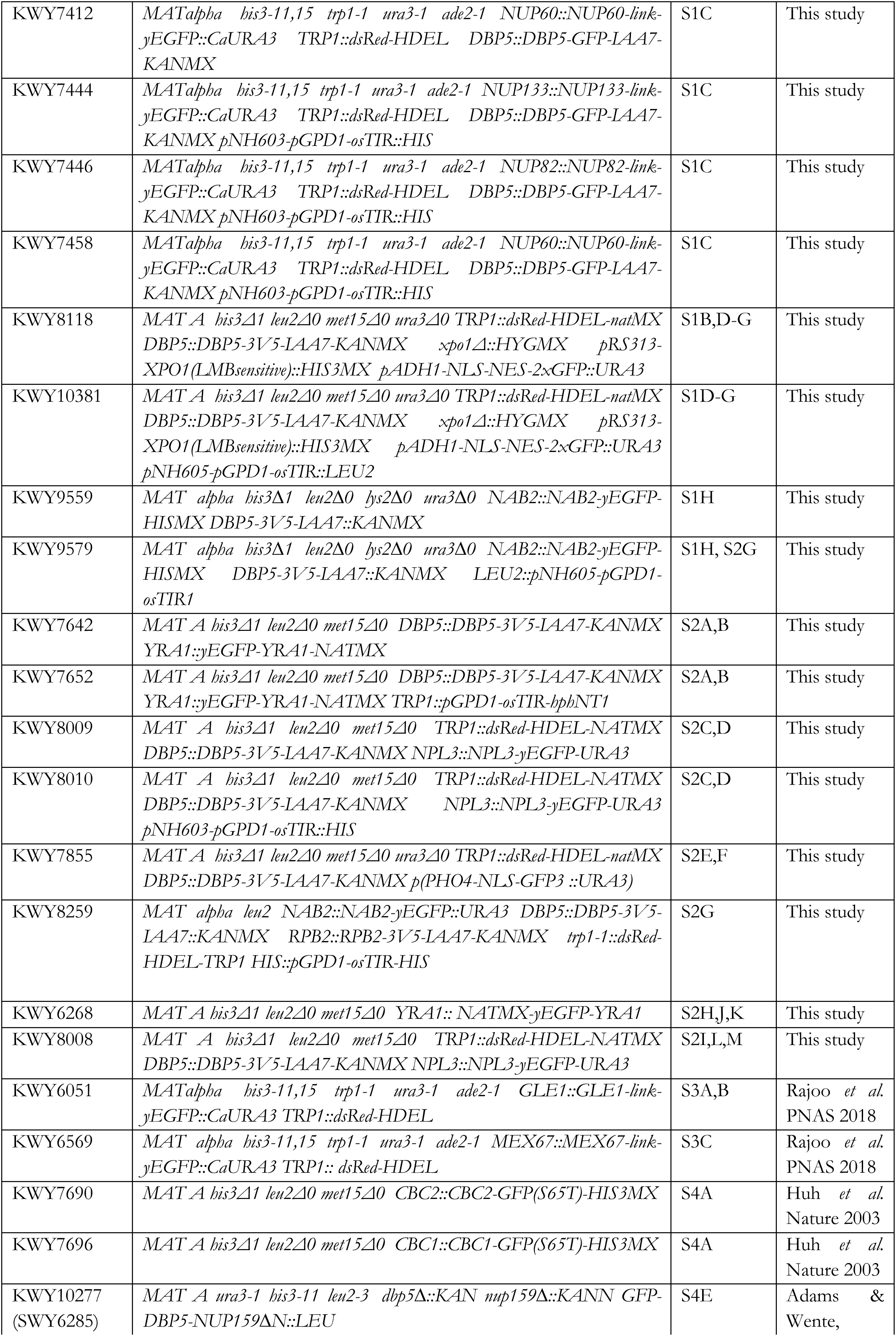
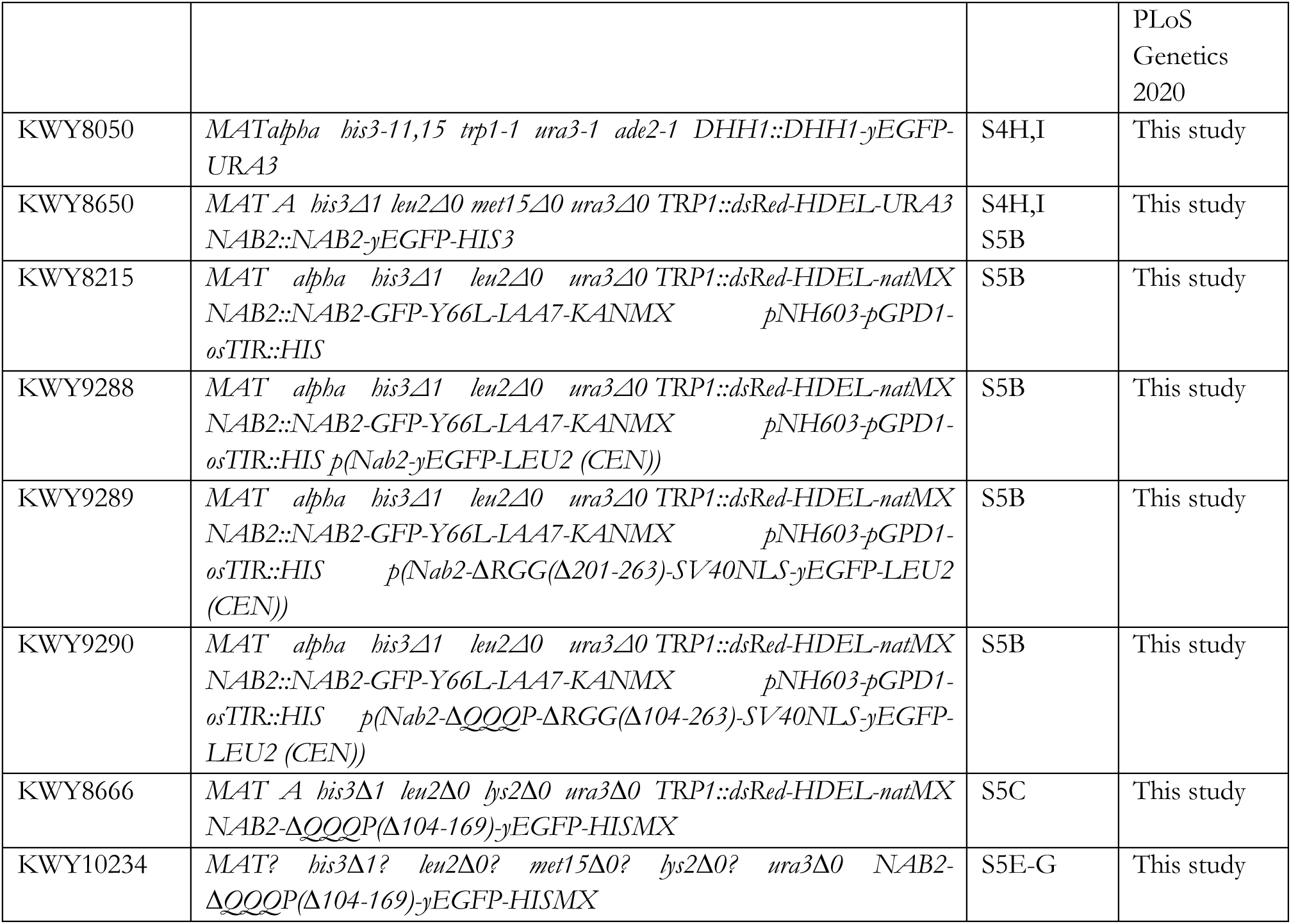
(Strain list)

**Supplemental Table 2.**
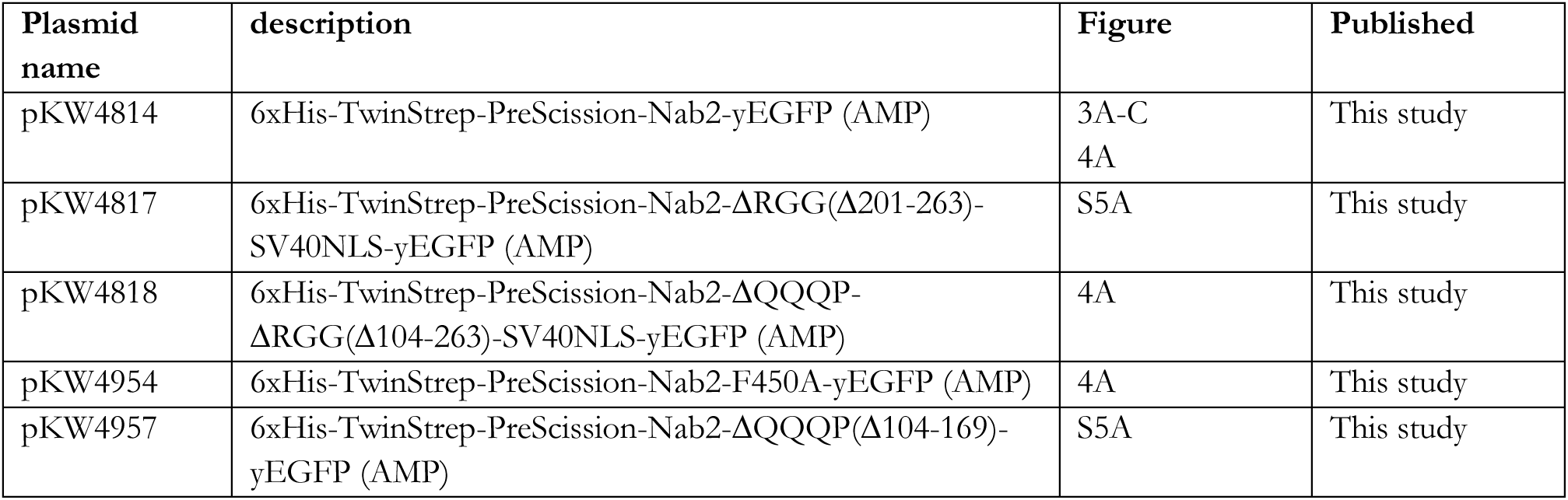
(Plasmid List)

**Supplemental Table 3.**
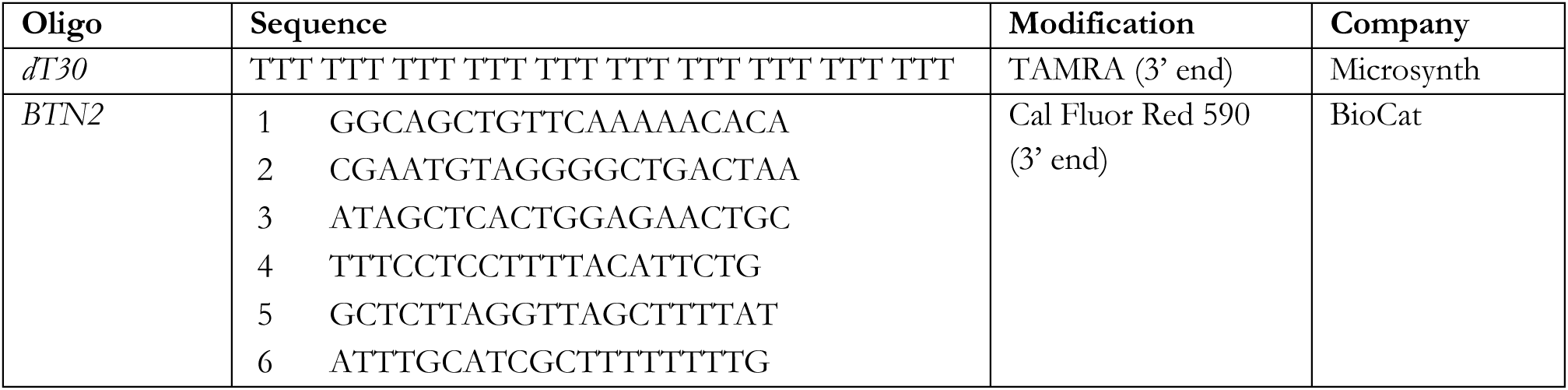
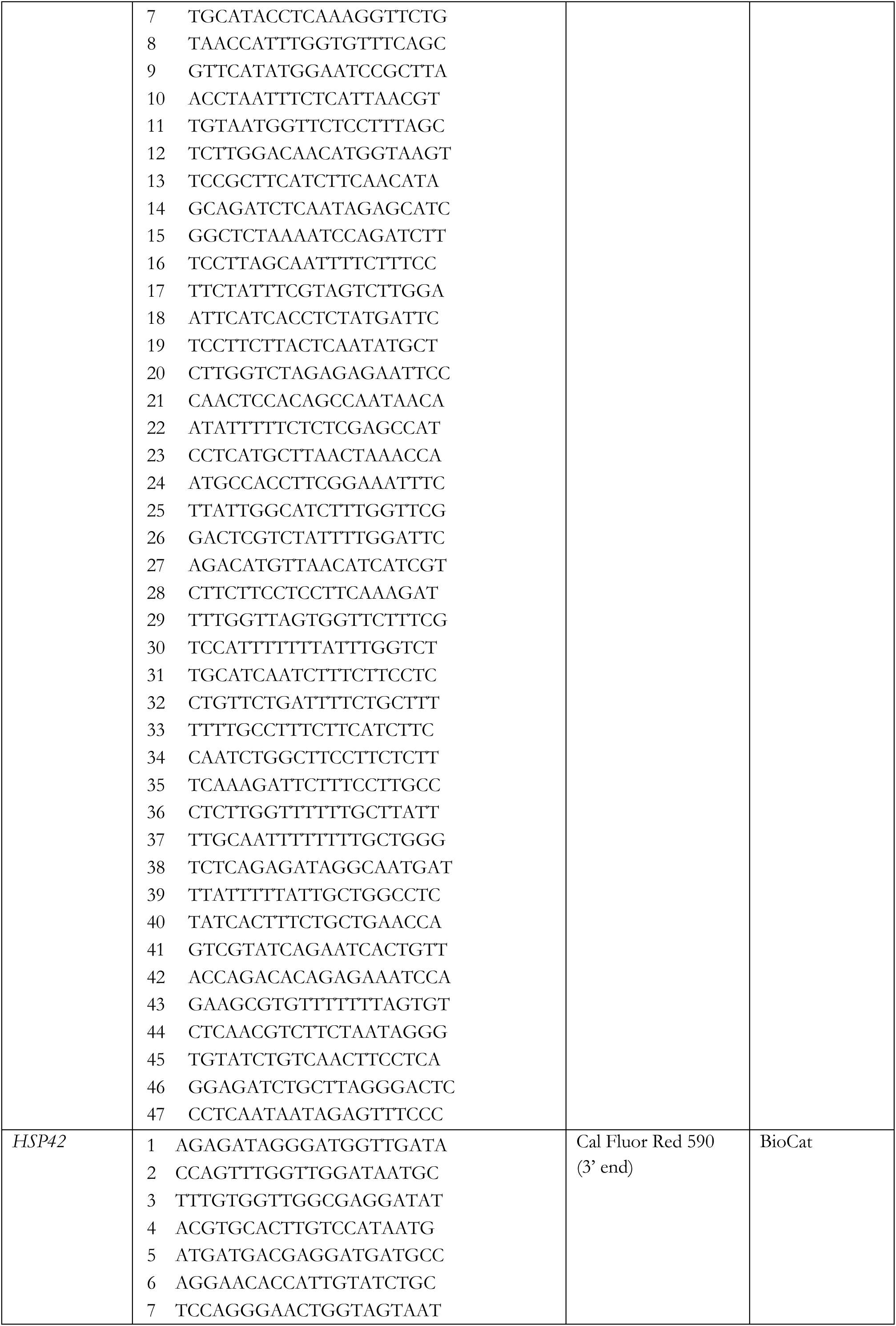
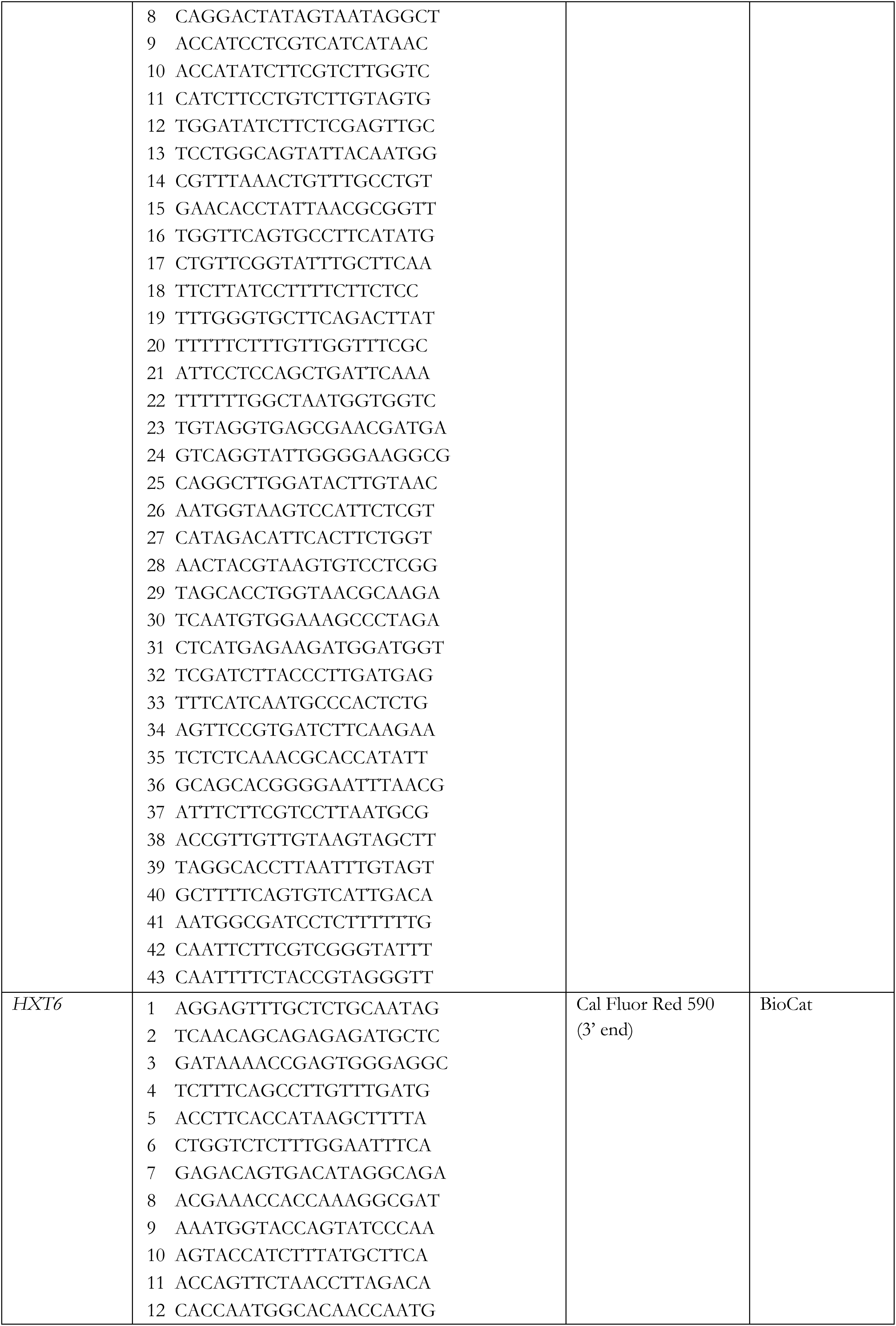
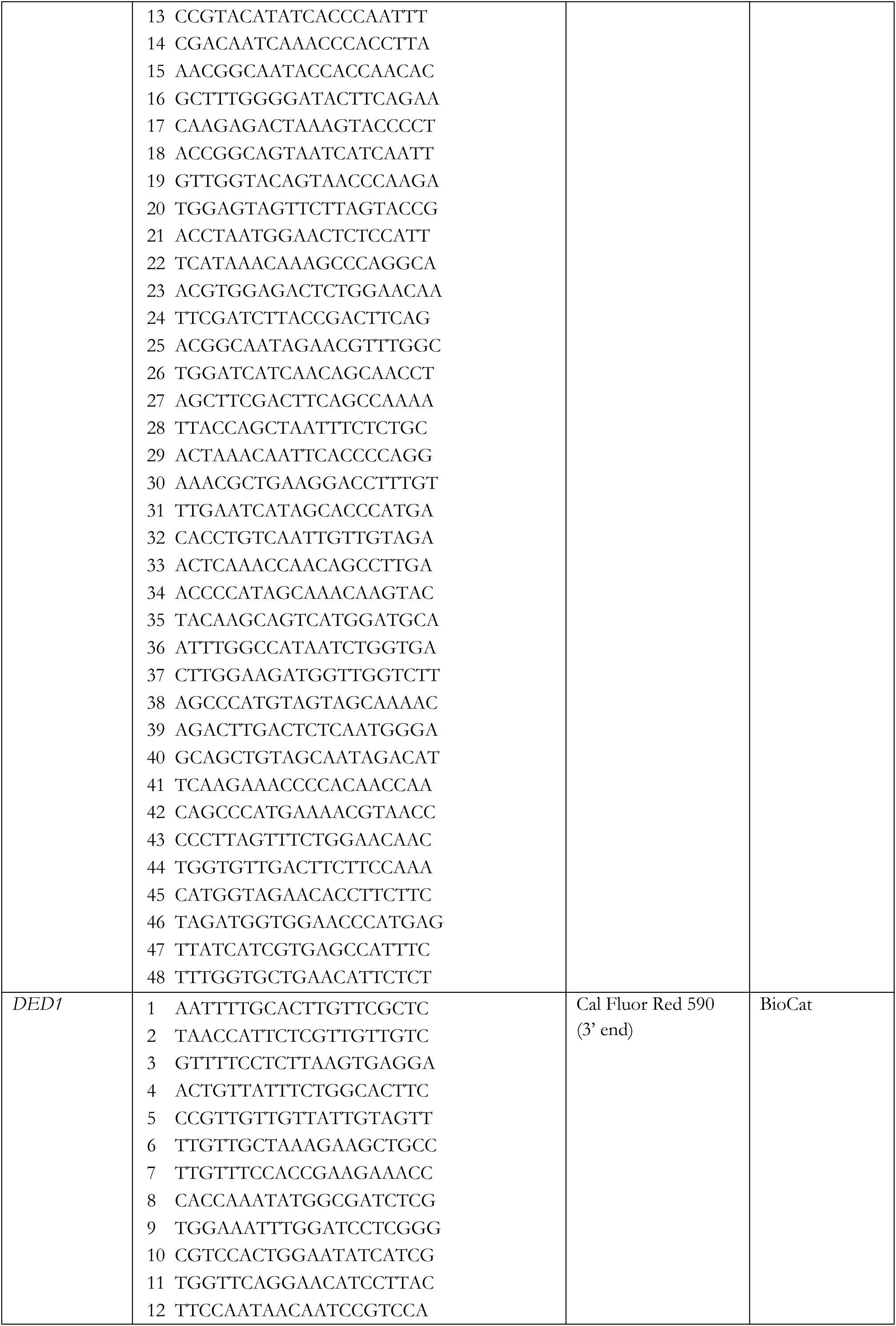
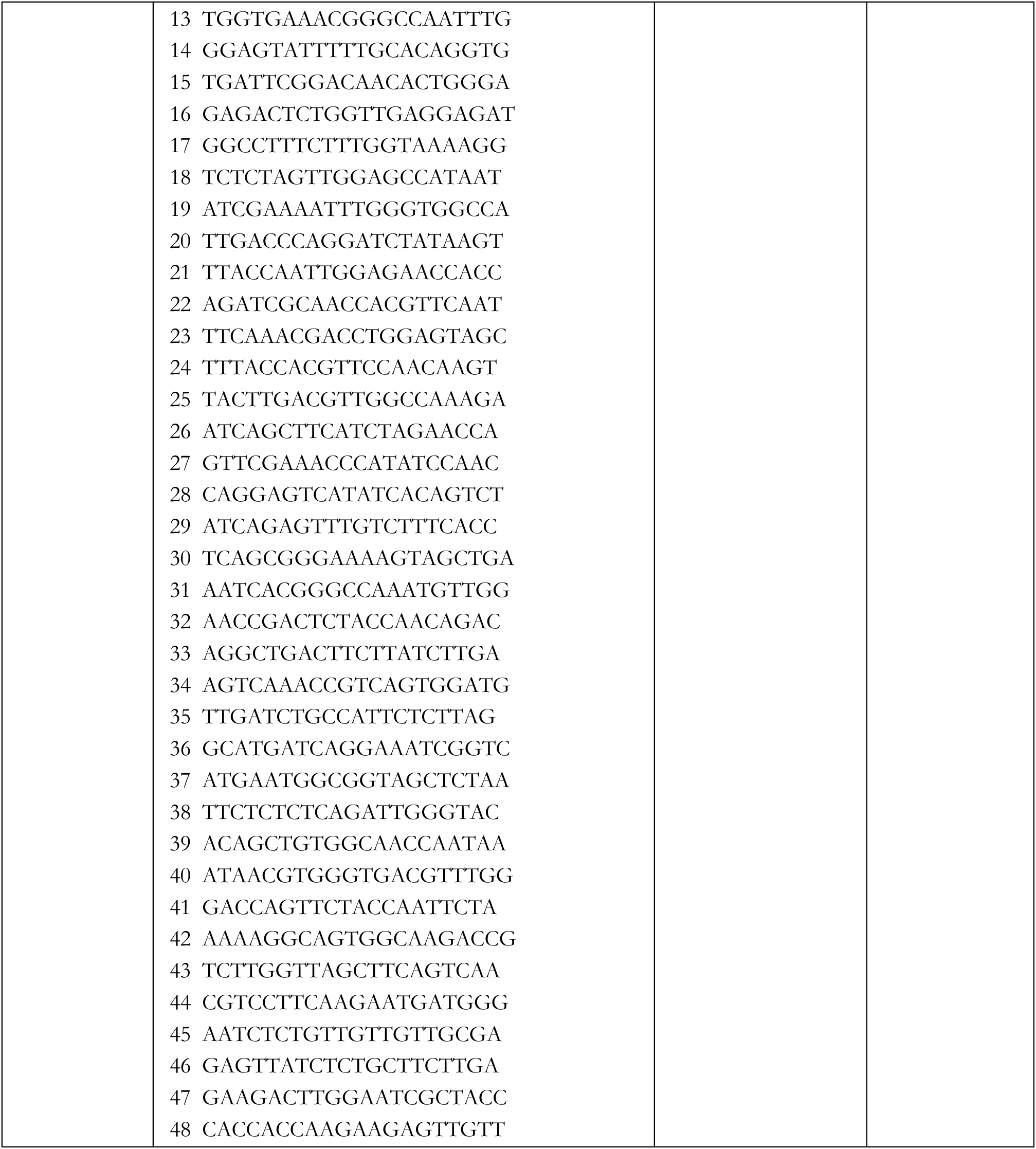
(Oligo list (smFISH))

### Yeast strain construction

*Saccharomyces cerevisiae* strains were constructed using standard yeast genetic techniques either by transformation of a CEN plasmid, a linearized plasmid, or of a PCR amplification product with homology to the target site ^71^. All yeast strains expressing a GFP-tagged protein were tagged with a yeast-optimized monomeric EGFP version (A206K).

Nab2^ΔQ^ and Nab2^ΔQ-ΔRGG-SV40NLS^ mutant strains were constructed using a CRISPR/Cas9 strategy ^72^. In brief, gRNA sites were determined using Benchling (settings: single guide of 20 nt length and an NGG PAM site (SpCas9, 3’ side)). For both Nab2^ΔQ^ and Nab2^ΔQ-ΔRGG-SV40NLS^ the cut site lies within the Q-rich region (gRNA regions amplified off pML104^72^: (A) ‘upstream’ gRNA: GTCCCTAGTTGTGGCTGAAGgttttagagctagaaatagcaagttaaaataaggctagtccg + cggaaacgctatacctccacc; (B) ‘downstream’ gRNA: gcgaggaagcggaagagcg + ctagctctaaaacCTTCAGCCACAACTAGGGACgatcatttatctttcactgcggagaag; KWY9296 was transformed with stitched gRNA regions, SwaI-digested pML104 and PCR amplification products with homology to the target site (ΔQ: amplified homology product from genomic DNA excluding DNA region coding for aa104-169; ΔQ-ΔRGG-SV40NLS: amplified homology product from plasmid pKW4774 where DNA region coding for aa104-263 was replaced by the SV40NLS sequence (AAAGGCCTAAGAAGAAGAGGAAAGTATTA)).

### Yeast cell culturing

Cells were cultured in synthetic complete medium containing 2% glucose (Formedium, CSC0205) (for microscopy, RNA FISH and nuclear extraction assay). Cells were inoculated from saturated overnight cultures into fresh medium and grown at 25°C to OD_600_ 0.6–0.8. For acute glucose withdrawal, cells were then washed 3 times in synthetic complete medium without glucose, resuspended in the same medium and cultured for the indicated time (30 min or 24 hours, shaking at 150 rpm). For long-term incubation in synthetic complete medium without glucose (up to 5 days), cultures were covered with an O_2_- permeable membrane to prevent evaporation.

Acute degron depletion was performed by addition of auxin (500 μM indole-3-acetic acid (IAA); 500 mM stock in ethanol) for the indicated time points (typically 1 or 2 hours). As a solvent control, cells were treated with ethanol.

### RNA Fluorescence *in situ* hybridization (poly(A) RNA FISH and single molecule FISH (smFISH))

Probe design: DNA probes coupled to CAL Fluor Red 590 (Stellaris Biosearch Technologies, synthesized by BioCat) were used for smFISH, targeting gene-specific moieties for single molecule visualization. dT30 DNA probes coupled to TAMRA (synthesized by Microsynth AG) targeting poly(A) stretches were used for poly(A) RNA visualization.

Methodology: Samples were processed as described in ^22^. Briefly, cultures were fixed for 30 min with 4% paraformaldehyde (Electron Microscopy Sciences, 15714). Cells were washed once with buffer B (1.2 M sorbitol and 100 mM KHPO4, pH 7.5, at 4°C), then spheroplasted for 20 min using 1% 20T zymolyase in 1.2 M sorbitol, 100 mM KHPO4, pH 7.5, 20 mM vanadyl ribonuclease complex (New England Biolabs, S1402S), and 20 μM β-mercaptoethanol. To stop the spheroplasting reaction, cells were washed once with buffer B, and then transferred into pre-hybridization buffer (10% formamide (Merck Millipore, S4117), 2x SSC (saline-sodium citrate buffer; Life Technologies, AM9624). Separately, 0.5 μL of probe mix (stock 25 μM) was mixed with 2 μL of salmon-sperm DNA (10 mg/ml; Life Technologies, 15632-011) and 2 μL yeast transfer RNA (10mg/ml; Life Technologies, AM7119). The probe mix was denatured in 50 μL of hybridization buffer F (20% formamide and 10mM NaHPO4, pH7.0) for 3 min at 95 °C, and then mixed with 50 μL hybridization buffer H (4x SSC, 4 mg/ml acetylated BSA (Sigma Fluka Aldrich, B8894-1.5mL), 20 mM vanadyl ribonuclease complex). This final hybridization mixture was added to the pre-hybridized cell pellet (which corresponds to approx. 3 OD_600_) and incubated overnight at 37 °C. The next day, cells were washed in 1x PBS/DAPI and incubated for 3 min, washed again in 1x PBS and then stored at 4 °C.

### Microscopy for live cell imaging and RNA FISH

For live cell imaging and NuRIM fluorescence intensity quantification experiments, cells were transferred to a 384-well plate (Brooks, Matrical MGB101-1-2-LG-L) treated with Concanavalin A (stock solution of 1 mg/ml, air-dried), and spun down for 1 min at 100 rcf.

For RNA FISH, approx. 1 OD_600_ of fixed cells were spread onto a glass slide using a 12mm cover slip (Huber, 10.0360.52) to obtain a single layer of cells prior to imaging. Alternatively, fixed cells were transferred to a 384-well plate (Brooks, Matrical MGB101-1-2-LG-L) treated with Concanavalin A (stock solution of 1 mg/ml, air-dried), spun down for 1 min at 100 rcf, then washed twice with 1x PBS to obtain a single layer of cells.

All imaging experiments were performed at 25 °C. Microscopy was performed using an inverted epi-fluoresence microscope (Nikon Ti) equipped with a Spectra X LED light source and a Hamamatsu Flash sCMOS camera using a 100X Plan-Apo objective NA 1.4 and NIS Elements software.

### Live single molecule RNA analysis

PP7sl-RNA tracking in Fig. 1C was performed as follows:

1. Image denoising: Noise2Void^73^ was used to denoise the optical microscopy data of the RNA particles. The algorithm has a deep learning approach based on noisy images without the need for clean target images. The U-Net was trained with a 3×3 kernel size over 100 epochs with 100 steps per epoch. The batch size was 128.
2. Single molecule tracking analysis: The localization of the single RNA transcripts was done using TrackMate, an open-source ImageJ plugin^74^. Sub-pixel localization was achieved with the Laplacian of Gaussian (LoG) detector with a transcript radius of 1.5 pixels and a threshold of 190. The radius was chosen to be slightly smaller than the expected transcript size. The intensity threshold for the localizations was defined to minimize false localizations. The resulting single-molecule data including the transcript position, mean intensity and particle quality was imported into Python.
3. Single molecule tracking was done with Trackpy^75^, a Python implementation of the Crocker-Grier particle tracking algorithm ^76^. The maximum displacement per frame was set to 5px and the memory parameter was set to 2 frames (a single transcript could disappear for two frames and still maintain its ID).
4. The nuclear envelope data was segmented using the MATLAB functions *adaptthresh* and *imbinarize* from the Image Processing Toolbox. The segmentation was then refined using the watershed transform.
5. MSD analysis: the detected single molecules were separated into cytoplasmic and nuclear compartments based on the above nuclear envelope segmentation. The mean squared displacement (MSD) analysis of the nuclear RNA was done using *msdanalyzer* ^77^.

PP7sl-RNAs in Fig. 7E,F were classified as static if the majority of RNA signal within a single nucleus showed a mobility of <105.6 nm^2^ (=5×5 pixels with 67nm per pixel) within the full 60 seconds of imaging. If particles showed mobility larger than 105.6nm^2^ they were classified as dynamic.

### Live cell imaging and RNA FISH image analysis

Images were processed using FIJI software and plots were generated in GraphPad Prism 8 (GraphPad). Nab2 focus quantification and mRNA quantification of RNA FISH experiments were performed using the FIJI adaptiveThr function (ImageJ plugin by Quingzong Tseng) with a weighted mean and strict thresholds to remove background and to isolate individual mRNAs (smFISH) and GFP foci. Statistical analysis in Figure 4D,E and 7A,B and D was performed with the statistical language R (version 4.3.1), and functions from the stats package or the tidyverse package^78^ were used. Raw intensity per cell was calculated as the ratio of the integrated density obtained from Fiji (Int_den) and the total cell count per image. The nuclear signal was determined by subtracting the cytoplasmic signal from the whole cell signal. The raw intensity per cell for each replicate was median normalized, with a distribution median corresponding to the median RNA signal over all biological replicates for a given FISH probe. The data did not follow a normal distribution, therefore non-parametrical Wilcoxon test was performed to compare the normalized intensity per cell and the cytoplasmic to nuclear (C/N) RNA ratio. The nonparametric Cohen’s d-consistent effect size was calculated using the *gammaEffectSize()* function in the cfaTools package (https://centerforassessment.github.io/cfaTools/) over the quantile range [0.3, 0.7]; the mean value was used for further analysis. The relationship between RNA count (from nuclear RNAseq analysis in Fig. 5D) and effect size was evaluated using Spearman’s rank correlation.

For NuRIM quantification, images were processed using NuRIM code^38,79^ in MATLAB. Briefly, this method uses the HDEL-DsRed signal to generate a mask that segments the nuclear envelope. This mask is then transferred to the GFP channel to extract average intensity information from this mask.

### Fluorescence recovery after photobleaching (FRAP)

A) Yeast cells: 200 µL of a culture with OD 0.6-0.8 was added to a Concanavalin A-treated imaging well (IBIDI, µ- Slide 8-well, 80827) and spun down for 1 min at 100 rcf. Microscopy was performed on a Leica TCS SP8 STED/FLIM/FCS or a Leica TCS SP8-AOBS microscope using a 63x 1.4NA Oil HC PL APO CS2 objective. A bidirectional scanner at a speed of 8,000 Hz, NF488/561/633, an Airy unit of 1.5, and a FRAP booster for bleaching were applied for every FRAP experiment using a PMT detector (500–551 nm). An image size of 504 x 50 pixel and a zoom of 4 were used together with line accumulation of three, yielding a frame rate of 74.07 frames/s with a pixel dwell time of 73 ns. 50 prebleach and 300 postbleach frames were acquired. A 488-nm argon laser line was used at 30% power. Typically, an elliptical region covering approximately one half of the cell nucleus was bleached, except for Figure 1J-L, where, in addition, elliptical regions covering approximately one third or one quarter of the cell nucleus were bleached. The bleaching was performed using 80% argon laser power for 195 ms. Imaging was conducted with 1.5% laser intensity (with a gain of 570 for Nab2-GFP and GFP-Yra1, 560 for Npl3-GFP, 650 for NLS-3xGFP and Mex67-GFP) to illuminate the GFP during prebleach and postbleach acquisition.
B) *In vitro* droplets: Protein droplets were set up according to ^34^. In brief, reactions were pipetted in 384-well microscopy plates (Brooks, Matrical MGB101-1-2-LG-L). 3 µL recombinant GFP-tagged protein (stock 150 µM to a final concentration of 20 µM) was mixed at room temperature with 20 µL of a master mix (14.5 µL LSB-100 (100 mM KCl, 30 mM HEPES-KOH pH 7.4, 2 mM MgCl_2_), 0.9 µL CRM reconstitution mix (40 mM ATP, 40 mM MgCl_2_, 200 mM creatine phosphate, 70 U ml^−1^ creatine kinase), 1.8 µL ATP/MgCl2 (100 mM), 0.9 µL BSA (10 mg mL^−1^), 0.9 µL poly(A) RNA (1 mg mL^-1^), 0.9 µL 1M HEPES pH 6.8, 0.1 µL RNasin Plus (Promega). Microscopy was performed on a Leica TCS SP8 STED/FLIM/FCS or a Leica TCS SP8-AOBS microscope using a 63x 1.4NA Oil HC PL APO CS2 objective. A bidirectional scanner at a speed of 8,000 Hz, NF488/561/633, an Airy unit of 1.5, and a FRAP booster for bleaching were applied for every FRAP experiment using a PMT detector (500– 551 nm). An image size of 504 x 50 pixel and a zoom of 4 were used together with line accumulation of three, yielding a frame rate of 74.07 frames/s with a pixel dwell time of 73 ns. 50 prebleach and 800 postbleach frames were acquired. A 488-nm argon laser line was used at 30% power. Imaging was conducted with 1.5% laser intensity (with a gain of 570) to illuminate the GFP during prebleach and postbleach acquisition. The bleach was performed using 80% argon laser power for 195 ms. The bleaching was performed by defining an elliptical region covering approximately one half of the droplet.
C) Image analysis:

1) yeast cells: Fluorescence recovery in the bleached region was evaluated by subtracting the extracellular background from the intensity of the bleached region, and the values were then bleach-corrected by normalizing for total cell intensity ^80^.
2) droplets: Fluorescence recovery in the bleached region was evaluated by subtracting the background signal outside the droplet from the intensity of the bleached region, and the values were then bleach-corrected by normalizing for total droplet intensity. All curves were fitted with an exponential function in a custom-written MATLAB script. Plots were generated in GraphPad Prism 8 (GraphPad).

### Nucleus extraction

The protocol was adapted from ^81^ and ^53^. Cells were inoculated in a 10 mL pre-culture of synthetic complete media containing 2% glucose (SCD, Formedium, CSC0205), and cultured for 8-10 hrs at 25 °C. Cells were then back-diluted in 500 mL SCD to OD_600_ 0.006, grown for 16-18 hrs at 25 °C to OD_600_ 0.8 and harvested onto a cellulose ester membrane filter (Whatman 10400921) using a vacuum filtration pump. Cells were then washed from the membrane with 40 mL ddH_2_O into a 50 mL Falcon tube, spun down for 2 min at 845 rcf, resuspended in 25 mL buffer 1 (0.1M Tris-HCl pH7.4, 10mM DTT) and incubated shaking at 30 °C for 20 min in a horizontal shaker at 150 rpm. Cells were then spun down for 2 min at 845 rcf, washed in 10 mL buffer 2 (20mM KPi pH 7.4, 1.2M Sorbitol), and spun down again for 2 min at 845 rcf. For cell wall digestion (spheroplasting), cells were resuspended in 50 mL buffer 3 (20mM KPi pH 7.4, 1.2M Sorbitol, 10 mM ribonucleoside vanadyl complex (NEB S1402S, pre-warmed at 65 °C), 0.08mg/ml Zymolyase-20T (Amsbio 326921)) and incubated for 60 min at 30 °C in a rotation wheel. Spheroplasts were spun at 3000 rcf for 2 min, washed in 10 mL buffer 4 (20mM KPi pH 6.4, 1.2M Sorbitol), and spun again at 3000 rcf for 2 min. To release nuclei, spheroplasts were carefully resuspended in 20 mL buffer F (20% Ficoll, 20mM KPi pH 6.4, 5 mM RVC, 0.005mg/ml Pepstatin A, 1x protease inhibitor cocktail (PIC, Sigma P8215-5mL), 250 µg/mL DAPI) and put on ice. To mechanically assist nuclei release, the lysate was then processed with 5 slow strokes in a dounce homogenizer on ice (this step is not essential as most of the nuclei are already released upon resuspension in buffer F). To separate nuclei from remaining cells/spheroplasts, the lysate was spun for 5 min at 3460 rcf at 4 °C. Afterwards, the supernatant was carefully removed with a 25 mL pipet until ca. 2 mL was left (to avoid pipetting the cell/spheroplast pellet) and transferred to a fresh 15 mL Falcon tube on ice. For nucleus enrichment and removal of other organelles, a sucrose gradient was prepared as follows: ice-cold 5 mL 2.5 M sucrose was added to the bottom of an open-top plastic tube (Beckman-Coulter 344058), carefully topped up with ice-cold 5 mL 1.875 M sucrose, which in turn was carefully topped up with ice-cold 5 mL 1.7 M sucrose. Lastly, the lysate supernatant was carefully added on top of the 1.7 M sucrose layer, and the full tube was transferred to a pre-cooled metal holder of a SW32_Ti ultracentrifuge rotor (Beckman-Coulter). Tubes were spun for 30 min at 103.000 rcf at 4 °C, which resulted in ‘cloudy’ layer formation between the 1.7 M and 1.875 M sucrose interface (containing mostly cytoplasmic organelles, fraction 1) and between the 1.875 M and 2.5 M sucrose layer (containing nuclei, fraction 2). The enriched nucleus fraction 2 (typically around 4 mL) was then harvested by carefully pipetting away the top layers. To reduce the sucrose molarity, 4 mL of ice-cold buffer 5 (20mM KHPO4 pH 6.45) was added to the nucleus fraction 2, mixed and spun for 4 min at 3460 rcf at 4 °C. The supernatant was carefully removed with a 10 mL pipet until ca. 2 mL was left to separate any remaining cell debris from the nuclear fraction. 2×500 µL aliquots were taken for DNA content determination as a proxy for the number of harvested nuclei. Enriched nuclei were then immediately processed for RNA extraction (see below).

### RNA extraction from nuclear sucrose gradient fraction

The enriched nucleus fraction was transferred to a fresh open-top plastic tube (Beckman-Coulter 344058) and spun for 30 min at 20.000 rcf for 30 min in SW32Ti rotor. The pellet was resuspended in 500 µL RNase-free TES buffer (10 mM TrisHCl pH 7.5, 10 mM EDTA, 0.5% SDS) and transferred to screw-cap tubes. 500 µL acid-saturated phenol was added, the mixture was vortexed and incubated for 30 min in a thermo block at 65 °C (shaking at 150 rpm). Afterwards, the tube was put on ice for 5 min and then spun for 10 min at 13.000 rcf at 4 °C. The upper aqueous phase was transferred to a fresh RNase-free tube, mixed with 500 µL acid-saturated phenol, vortexed and spun for 5 min at 13.000 rcf at 4 °C. The upper aqueous phase was transferred to a chloroform-resistant screw-cap tube, mixed with 500 µL chloroform, vortexed and spun for 5 min at 13.000 rcf at 4 °C. The upper aqueous phase was transferred to a fresh RNase-free tube, mixed with 60 µL 3M NaOAc pH 5.2 and 700 µL isopropanol, and incubated overnight at -20 °C. The next morning, the mixture was spun for 40 min at 13.000 rcf at 4 °C, the supernatant was decanted and 300 µL 75% EtOH was added, followed by centrifugation for 5 min at 13.000 rcf at 4 °C. The supernatant was removed and the pellet was air-dried for 10 min on ice, resuspended in nuclease-free H2O and stored at -80 °C.

### RNA extraction from whole cell lysates

Cells were inoculated in a 10 mL pre-culture of synthetic complete media containing 2% glucose (SCD, Formedium, CSC0205), and cultured for 8-10 hrs at 25 °C. Cells were then back-diluted in 400 mL SCD to OD_600_ 0.006, grown for 16-18 hrs at 25 °C to OD_600_ 0.8 and harvested onto a cellulose ester membrane filter (Whatman 10400921) using a vacuum filtration pump. Cells were scraped off the membrane with a metal rod and snap-frozen in a 50 mL Falcon tube containing liquid nitrogen. To lyse the cells, the cell pellet was resuspended in 400 uL TES buffer (10mM TrisHCl pH7.5, 10mM EDTA, 0.5% SDS), followed by addition of 400 uL acid-saturated phenol. The samples were incubated at 65°C for 1 hour with constant vortexing. Afterwards, the tube was put on ice for 5 min and then spun for 10 min at 20.000 rcf at 4 °C. The upper aqueous phase was transferred to a fresh RNase-free tube, mixed with 500 µL acid-saturated phenol, vortexed and spun for 5 min at 20.000 rcf at 4 °C. The upper aqueous phase was transferred to a fresh RNase-free tube, mixed with 400 µL chloroform, vortexed and spun for 10 min at 20.000 rcf at 4 °C. The upper aqueous phase was transferred to a fresh RNase-free tube, and the RNA was precipitated by adding 55uL 3M NaOAc pH5.2 and 550 uL isopropanol, and incubated at -20 °C for at least 2 hours. To pellet the RNA, the sample was spun for 30 min at 20.000 rcf at 4 °C, the supernatant was removed and the pellet was washed with 800 µL 80% EtOH, followed by centrifugation for 10 min at 20.000 rcf at 4 °C. The EtOH was completely removed, and the RNA pellet was resuspended in nuclease-free water.

### RNA seq library preparation and sequencing

For poly(A) selection, 150 ug of total RNA in 200 uL binding buffer (10 mM Tris-HCl pH 7.5, 0.5 M LiCl, 6.7 mM EDTA) was first denatured at 80°C for 2 min, and immediately incubated on ice. 150 μL of resuspended Dynabeads® Oligo(dT)_25_ (Invitrogen, 61002) were first washed twice with binding buffer, then mixed with the total RNA samples and incubated for 5 min at room temperature. The beads were then washed twice with washing buffer (10 mM Tris-HCl pH 7.5, 0.15 M LiCl, 1 mM EDTA), then resuspended in 40 μL 10 mM Tris pH 7 followed by incubation at 80°C for 2 min to elute the poly(A) RNA. The eluted poly(A) RNA was used to generate strand-specific sequencing libraries using the NEXTflex Rapid Directional Illumina RNA-Seq Library Prep Kit (BioO) according to the manufacturer’s instructions. The libraries were sequenced on an NextSeq2000 sequencer. Reads were adaptor-trimmed using cutadapt and mapped using Tophat2 following rRNA read subtraction using Bowtie2, and quantified using featureCounts. The read counts per gene were normalised to the total number of reads per sample.

RNA extracted from the nuclear sucrose gradient fraction was processed the same way with the following adjustments: 3 ng of exogenous poly(A)-containing *in vitro* transcribed RNAs (‘spike-ins’ (*E. coli trpS, oppF, holB, mrsP*)) were added to 50 ug of total RNA extracted from the wild type nuclear sucrose gradient fraction. To adjust for the number of harvested nuclei within a replicate, an adjusted amount of the spike-ins was added to the 50 µg of RNA extracted from each of the Nab2 mutant nuclear sucrose gradient fraction, based on the DNA content determination of the sucrose-fractionated harvested nuclei relative to the Nab2-GFP wild type. After sequencing, spike-in reads were summed per sample and normalized to the number of total reads. Sample normalization within a replicate was performed by calculating the spike-in ratios between Nab2-GFP (set to ‘1’), Nab2-GFP:Nab2^ΔQΔRGG-SV40NLS^-GFP and Nab2-GFP:Nab2^F450A^-GFP using the normalized spike-in reads. Spike-in correction of the raw counts-per-million (CPM) values was then performed by multiplying the spike-in ratio value of each sample with the CPM value of each gene. To reliably detect mRNAs in all cells in the RNA FISH confirmation experiments, a cutoff of reads >500 CPM (in the Nab2-GFP average of the 3 biological replicates) was introduced. These most abundant transcripts were then further processed in the gene ontology (GO) term analysis using DAVID (Database for Annotation, Visualization and Integrated Discovery) functional annotation tool with medium stringency, 2021 update ^82,83^.

### Protein expression and purification

Protein expression and purification was performed according to ^84^. All recombinantly expressed Nab2 proteins were amplified from yeast genomic DNA of the yeast strains used in this study and cloned into pETMCN-based expression vector with a cleavable N-terminal 6×His-TwinStrep tag plus a C-terminal mGFPA206K tag. Plasmids are listed in Supplementary Table 3. The plasmids were transformed into chemically competent E. coli BL21* under the selection of 0.1 mg/mL ampicillin. Pre-cultures were grown in LB at 37 °C overnight, and diluted 1:100 into rich medium the next morning. Cells were grown at 37 °C to an OD_600_ of 0.6 and induced with final 200 mM IPTG. Cells were then grown overnight at 18 °C, collected and resuspended in 30 ml lysis buffer (500 mM NaCl, 25 mM Tris-HCl pH 7.5, 10 mM imidazole, 10% glycerol, protease inhibitors) per cell pellet from 2L of culture. Cells were lysed by EmulsiFlex (Avestin). To extract the 6×His-tagged proteins from the lysate, the lysate was incubated with ∼5mL (bead volume) Ni^2+^ sepharose in gravity flow columns for 30 minutes, washed 3 times with lysis buffer (gravity flow) and eluted in 2mL fractions. Fractions containing the protein were supplemented with PreScission (3C) protease for cleavage of the 6xHis-TwinStrep tag, and dialysed using a 14 kDa MWCO membrane into storage buffer (200 mM NaCl, 25 mM Tris-HCl pH 7.5, 10 mM imidazole, 5% glycerol) overnight at 4°C. The next day, protein solutions were further concentrated to ∼ 100 µM and purified by size exclusion with a Superdex 200 column (SD200 16/60) on an AEKTA purifier (GE Life Sciences) in the final storage buffer (200 mM NaCl, 25 mM Tris-HCl pH 7.5, 10 mM imidazole, 5% glycerol). From the gel filtration run, clean fractions of the monomeric peak were collected, pooled, concentrated to approximately 1000 μM using Millipore Amicon Centrifugation units (30 kDa MWCO), snap frozen as 20 μL aliquots in siliconized tubes in liquid nitrogen and stored at −80 °C. Protein expression levels, His eluates and gel filtration fractions were analysed by SDS–PAGE. Dbp5ΔN constructs were purified according to the same protocol, using TEV instead of PreScission (3C) protease.

### *In vitro* droplet formation assay

A) Nab2-GFP wild type and mutant experimental setup: Protein droplets were set up according to ^34^. In brief, recombinant GFP-tagged protein solutions were freshly thawed from -80 °C, and spun down at 14.000 rpm for 2 minutes to precipitate and remove aggregates. Stock solutions were diluted to a 195 µM working stock solution with storage buffer and kept on ice. 15 µL of a master mix contained 10.875 µL low-salt buffer (LSB; variable: LSB-100 (100 mM KCl, 30 mM HEPES-KOH pH 7.4, 2 mM MgCl_2_) or LSB-150 (150 mM KCl, 30 mM HEPES-KOH pH 7.4, 2 mM MgCl_2_), as indicated in figure legend), 0.675 µL CKM reconstitution mix (40 mM ATP, 40 mM MgCl_2_, 200 mM creatine phosphate, 70 U ml^−1^ creatine kinase), 1.35 µL ATP/MgCl2 (100 mM, dissolved in 1M HEPES pH 6.8), 0.675 µL BSA (10 mg mL^−1^ in ddH_2_O), 0.675 µL poly(A) RNA (1 mg mL^-1^ in nuclease-free H_2_O), 0.675 µL 1M HEPES (variable: pH 6.6/6.8/7.0/7.2/7.4, as indicated in the figure legend), 0.075 µL RNasin Plus (Promega). Reactions were pipetted in 384-well microscopy plates (Brooks, Matrical MGB101-1-2-LG-L). For a final GFP-protein concentration of 23 µM, 2 μL of the 195 µM working stock was mixed with 15 µL of the master mix by first pipetting the protein into the well plate, then adding the master mix and slowly pipetting up and down 2-3 times to mix. With the last pipetting, the mixture was transferred to the glass bottom of the well. For a final concentration of 11.5 µM, 1 µL Nab2-GFP (from 195 µM stock) was mixed with 16 µL of the aforementioned master mix. For a final concentration of 34.5 µM, 3 µL Nab2-GFP (from 195 µM stock) was mixed with 14 µL of the aforementioned master mix. Droplets were imaged 20-30 min after setup at 25 °C. Microscopy was performed using an inverted epi-fluorescence microscope (Nikon Ti) equipped with a Spectra X LED light source and a Hamamatsu Flash 4.0 sCMOS camera using a 100X Plan-Apo objective NA 1.4 and NIS Elements software. Images were processed using FIJI software.
B) Nab2+Dbp5 experimental setup: 2 µl of a 100 µM Nab2 solution was mixed with a total of 3 µl Dbp5ΔN protein (WT or mutant) mixed with Dbp5 storage buffer (200 mM NaCl, 25 mM Tris pH 7.5, 5 % glycerol) to achieve the final concentration as indicated. The final volume of 5 µl protein solution was mixed with 20 µL master mix (16 µL LSB100 buffer (100 mM KCl, 30 mM HEPES-KOH pH 7.4, 2 mM MgCl_2_), 1 µL 10 mg/ml BSA, 1 µL 100 mM ATP / 100 mM MgCl_2_, 1 µl HEPES-KOH pH 6.2, 1 µL poly(A) 1 mg/mL). Reactions were incubated for 10 minutes at room temperature and images recorded as described above.

## Acknowledgements

We thank S. Wente and A. Corbett for providing yeast strains and plasmids, ScopeM and J. Kusch for microscopy support, and the Weis lab for critical comments on the manuscript. This work was supported by an EMBO long-term fellowship to S.H. and M.H. (ALTF 290-2014, EMBOCOFUND2012, GA-2012-600394 to S.H.; ALTF 870-2014 to M.H.). M.H. was supported by a Human Frontier Science Program (HFSP) postdoctoral fellowship (LT000914/2015) and an ETH postdoctoral fellowship (FEL-37-14-2). This work was supported by the Swiss National Science Foundation (SNF 31003A_179275 and CRSII5_193740 to K.W.).

## Competing Interest Statement

The authors have declared no competing interest.

## Author contributions

Conceptualization, S.H., M.H. and K.W.; Methodology, S.H., M.H., D.G. and K.W.; Investigation, S.H., M.H., C.P.D., D.M., M.Z., R.M.; Formal analysis, S.H., M.Z., A.O., L.M., F.U.; Writing – Original Draft, S.H.; Writing – Review & Editing, S.H., M.H., D.M., M.Z., R.M., A.O., F.U., S.K. and K.W.; Funding Acquisition, S.H., M.H. and K.W.; Resources, K.W.; Supervision, S.H. and K.W.

## Supplemental Figures

**1).**
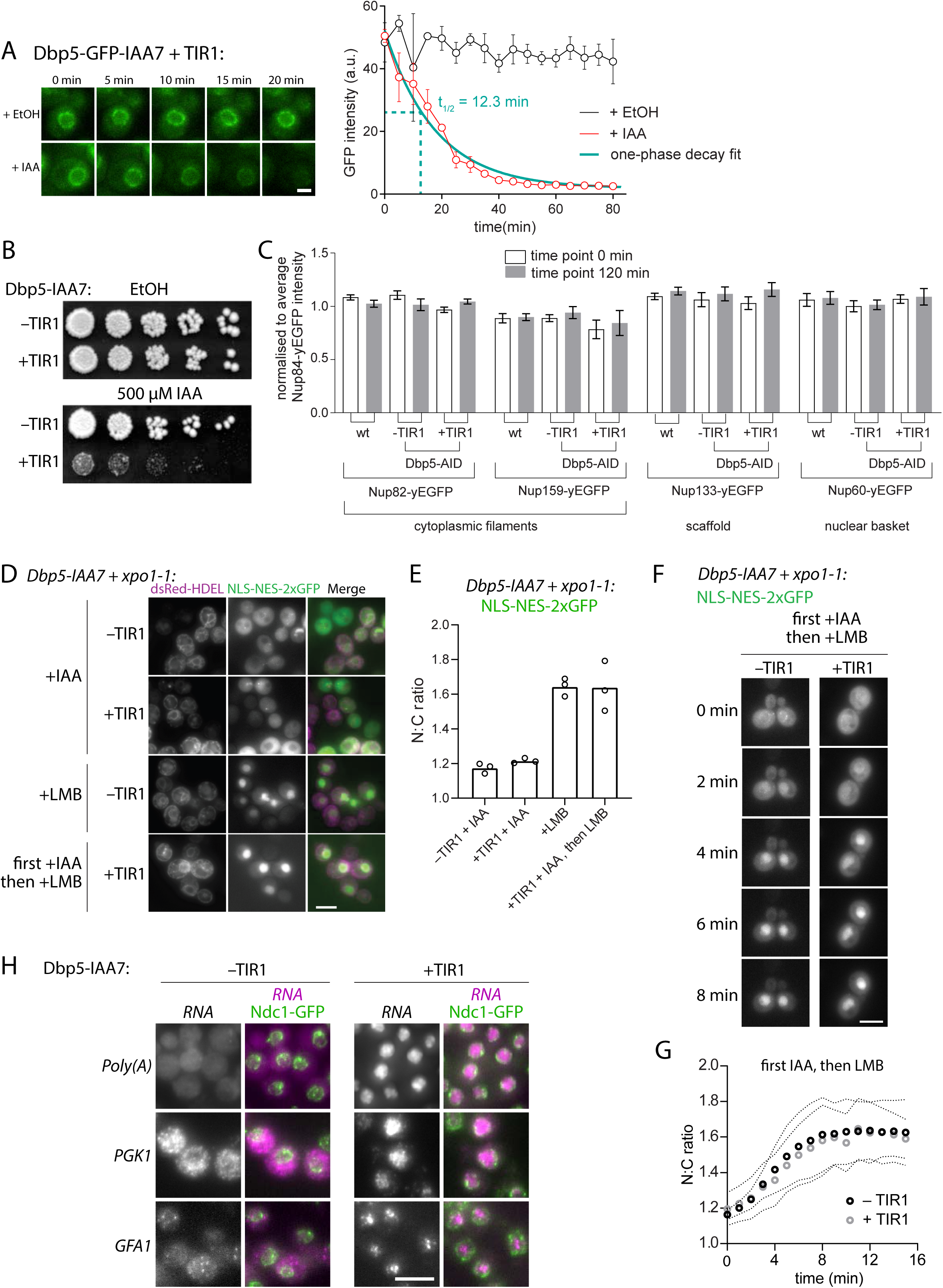
Acute depletion of Dbp5 does not affect NPC composition or NPC transport function; related to Figure 1. A) Left: Time lapse imaging of Dbp5-IAA7-GFP strain expressing TIR1 protein after addition of either 500 µM IAA or solvent control (EtOH). Scale bar 2 µm. Right: GFP intensity quantification of >200 cells per condition, green: one-phase decay fit, half-life = 12.3 min. B) Growth assay on rich medium plates containing 500 µM IAA or solvent control (EtOH). Cells of a Dbp5-IAA7 strain with or without expression of TIR1 protein were spotted as 1:5 serial dilutions (starting OD_600_ 0.2) and incubated at 25 °C for 2 days. C) Quantification of GFP-tagged nucleoporin (NUP) intensity at the nuclear rim in wild type *DBP5* background (wt), and in the presence (-TIR1) or absence (+TIR1) of the Dbp5 degron (Dbp5-AID) after incubation with 500 µM IAA for 0 min or 120 min. Shown are means ± SEM of three biological replicates, n>400 cells. D) Mid-log cultures of Dbp5-IAA7 *xpo1-1* cells expressing an NLS-NES-2xGFP reporter and the nuclear rim marker dsRed-HDEL with or without TIR1 were treated for 1 hr with either 500 µM IAA, 200 µM leptomycin B (LMB) or 1 hr 500 µM IAA followed by 1 hr 200 µM LMB before imaging. Scale bar 5 µm. E) Quantification of cells in B). Plot shows nuclear to cytoplasmic (N:C) GFP intensity ratio of NLS-NES-2xGFP reporter. Shown are means of three biological replicates, n>100 cells per replicate. F) Time lapse imaging of cells in D). Scale bar 5 µm. G) Quantification of time lapse imaging in F) shows nuclear to cytoplasmic (N:C) GFP intensity ratio of NLS-NES-2xGFP reporter. Means ± SEM (dotted lines) of three biological replicates, n>100 cells per replicate. H) RNA FISH against the indicated RNAs was performed in cells expressing Dbp5-IAA7 + Ndc1-GFP as a nuclear rim marker with or without TIR1. Samples were harvested after 2 hrs incubation with 500 µM IAA. Scale bar 5 µm.

**2).**
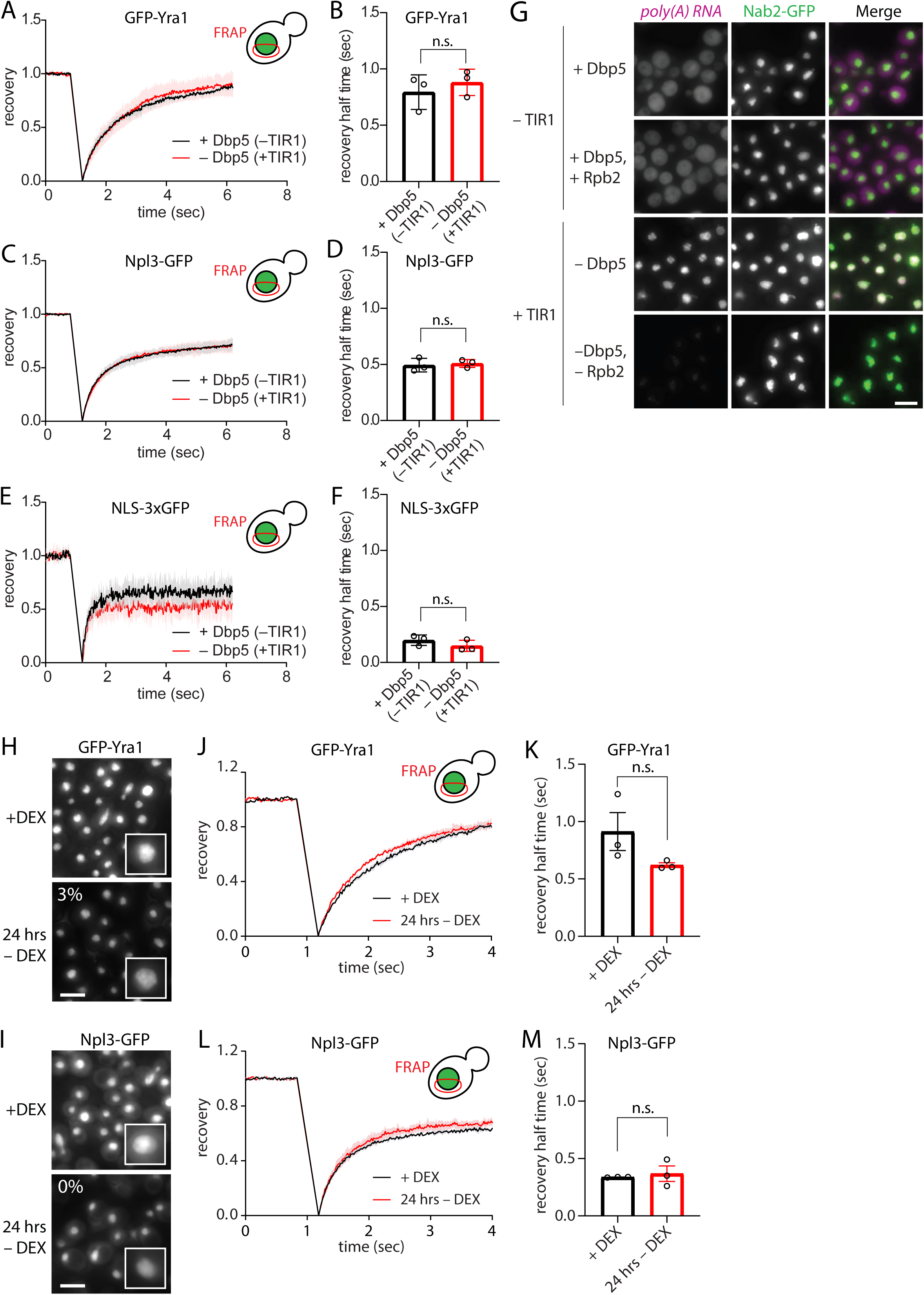
Acute depletion of Dbp5 or glucose stress does not alter mobility of Yra1, Npl3 or a nuclear GFP reporter; related to Figure 1 and 2. A) FRAP of GFP-Yra1 cells expressing the Dbp5-degron with or without TIR1. FRAP curves are normalized to prebleach and postbleach values. Mean curves ± SEM of three biological replicates with ≥13 cells per replicate are shown. B) Half time of recovery retrieved from fitting individual FRAP curves from A) with a single component model. Shown are means ± SEM of three biological replicates. ns p=0.4818, unpaired t-test. C) FRAP of Npl3-GFP cells expressing the Dbp5-degron with or without TIR1. FRAP curves are normalized to prebleach and postbleach values. Mean curves ± SEM of three biological replicates with ≥12 cells per replicate are shown. D) Half time of recovery retrieved from fitting individual FRAP curves from C) with a single component model. Shown are means ± SEM of three biological replicates. ns p=0.7217, unpaired t-test. E) FRAP of NLS-3xGFP cells expressing the Dbp5-degron with or without TIR1. FRAP curves are normalized to prebleach and postbleach values. Mean curves ± SEM of three biological replicates with ≥12 cells per replicate are shown. F) Half time of recovery retrieved from fitting individual FRAP curves from E) with a single component model. Shown are means ± SEM of three biological replicates. ns p=0.2642, unpaired t-test. G) RNA FISH against poly(A) RNA using an oligo(dT)30 probe was performed in cells expressing Dbp5-IAA7, or Dbp5-IAA7 and Rpb2-IAA7, with or without TIR1 protein. Samples were harvested after 2 hrs incubation with 500 µM IAA. Scale bar 5 µm. H) Representative images of GFP-Yra1 cells in +DEX or 24 hrs –DEX. Percentage indicates cells with microscopically visible GFP foci. Scale bar 5 µm. I) Representative images of Npl3-GFP cells in +DEX or 24 hrs –DEX. Percentage indicates cells with microscopically visible GFP foci. Scale bar 5 µm. J) FRAP of GFP-Yra1 cells in +DEX or 24 hrs –DEX. FRAP curves are normalized to prebleach and postbleach values. Mean curves ± SEM of three biological replicates with ≥15 cells per replicate are shown. K) Half time of recovery retrieved from fitting individual FRAP curves from E) with a single component model. Shown are means ± SEM of three biological replicates. ns p=0.154, unpaired t-test. L) FRAP of Npl3-GFP cells in +DEX or 24 hrs –DEX. FRAP curves are normalized to prebleach and postbleach values. Mean curves ± SEM of three biological replicates with ≥14 cells per replicate are shown. M) Half time of recovery retrieved from fitting individual FRAP curves from G) with a single component model. Shown are means ± SEM of three biological replicates. ns p=0.679, unpaired t-test.

**3).**
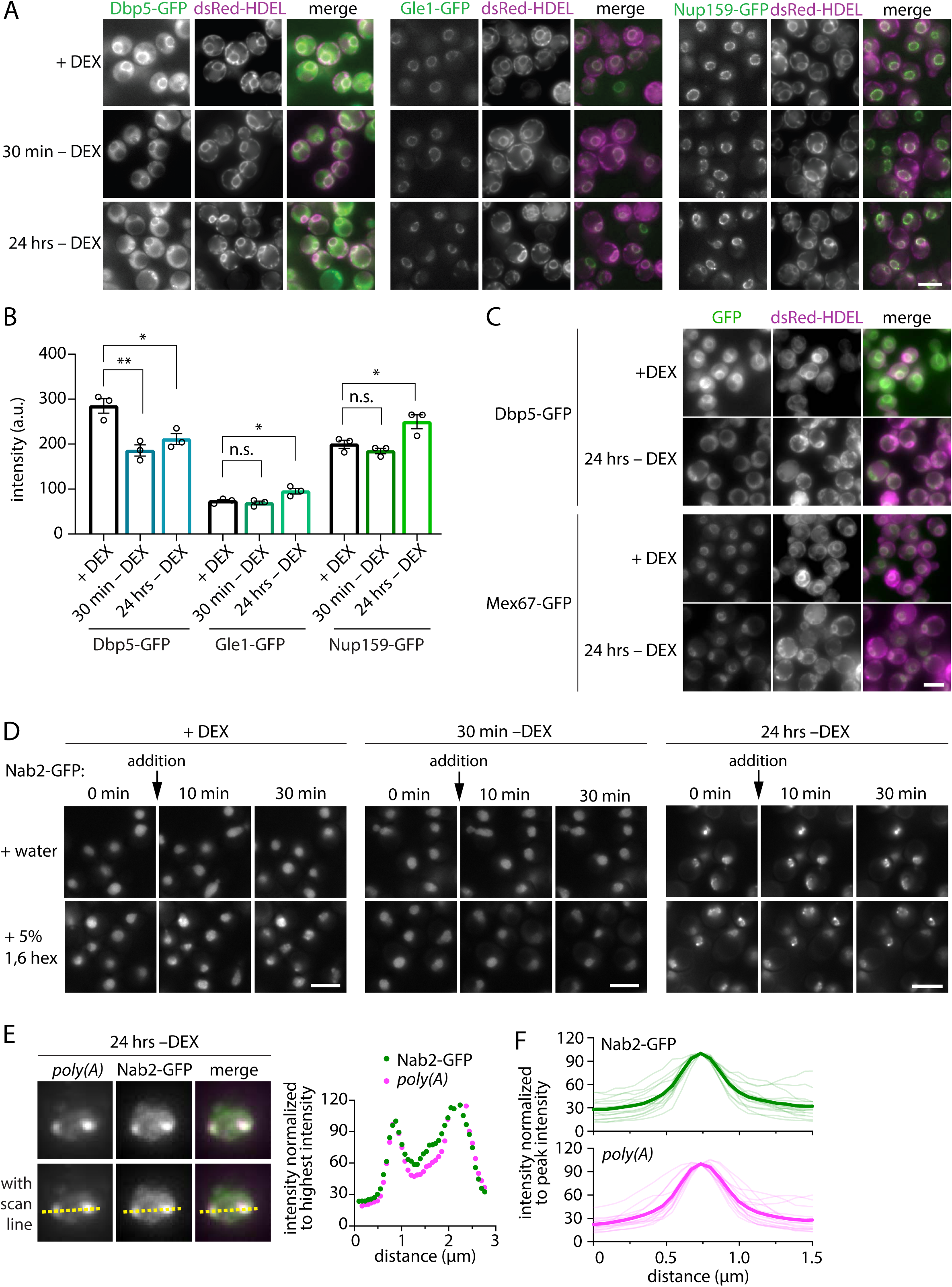
Glucose stress does not affect Gle1, Nup159 or Mex67 localization; Nab2 focus formation is mildly affected upon addition of 1,6 hexanediol and coincides with RNA focus formation *in vivo*; related to Figure 2. A) Cells expressing Dbp5-GFP, Gle1-GFP or Nup159-GFP, and dsRed-HDEL were grown in medium + DEX or acutely shifted for 30 min or 24 hrs into medium without DEX. Scale bar 5 µm. Dbp5-GFP data is also shown in Fig. 3A. B) Quantification of Dbp5-GFP, Gle1-GFP or Nup159-GFP intensity at the nuclear envelope by NuRIM in the conditions shown in A). Shown are means ± SEM of three biological replicates. At least 350 cells were analyzed per condition and replicate. Dbp5-GFP data also shown in Fig. 3A. Dbp5: ** p=0.0082 (+DEX vs. 30 min –DEX), * p=0.022 (+DEX vs. 24 hrs –DEX); Gle1: n.s. p=0.4342 (+DEX vs. 30 min –DEX), * p=0.0368 (+DEX vs. 24 hrs –DEX); Nup159: n.s. p=0.2453 (+DEX vs. 30 min –DEX), * p=0.0499 (+DEX vs. 24 hrs –DEX); unpaired t-test. C) Cells expressing Dbp5-GFP or Mex67-GFP, and dsRed-HDEL were grown in medium + DEX or acutely shifted into medium without DEX and incubated for 24 hrs. Scale bar 5 µm. D) Time lapse imaging of Nab2-GFP cells that were treated with water or 5% 1,6-hexanediol in +DEX (left), 30 min –DEX (middle) or 24 hrs –DEX (right). Scale bar 5 µm. E) Line scan of representative Nab2-GFP nucleus with nuclear poly(A) accumulation. Right: Intensity profile represents intensity along yellow dotted line, GFP and poly(A) signal normalized to maximum intensity. F) Line scans of 15 randomly chosen Nab2-GFP nuclei with nuclear poly(A) accumulation. GFP and poly(A) signal normalized to maximum intensity. Bold line represents average intensity.

**4).**
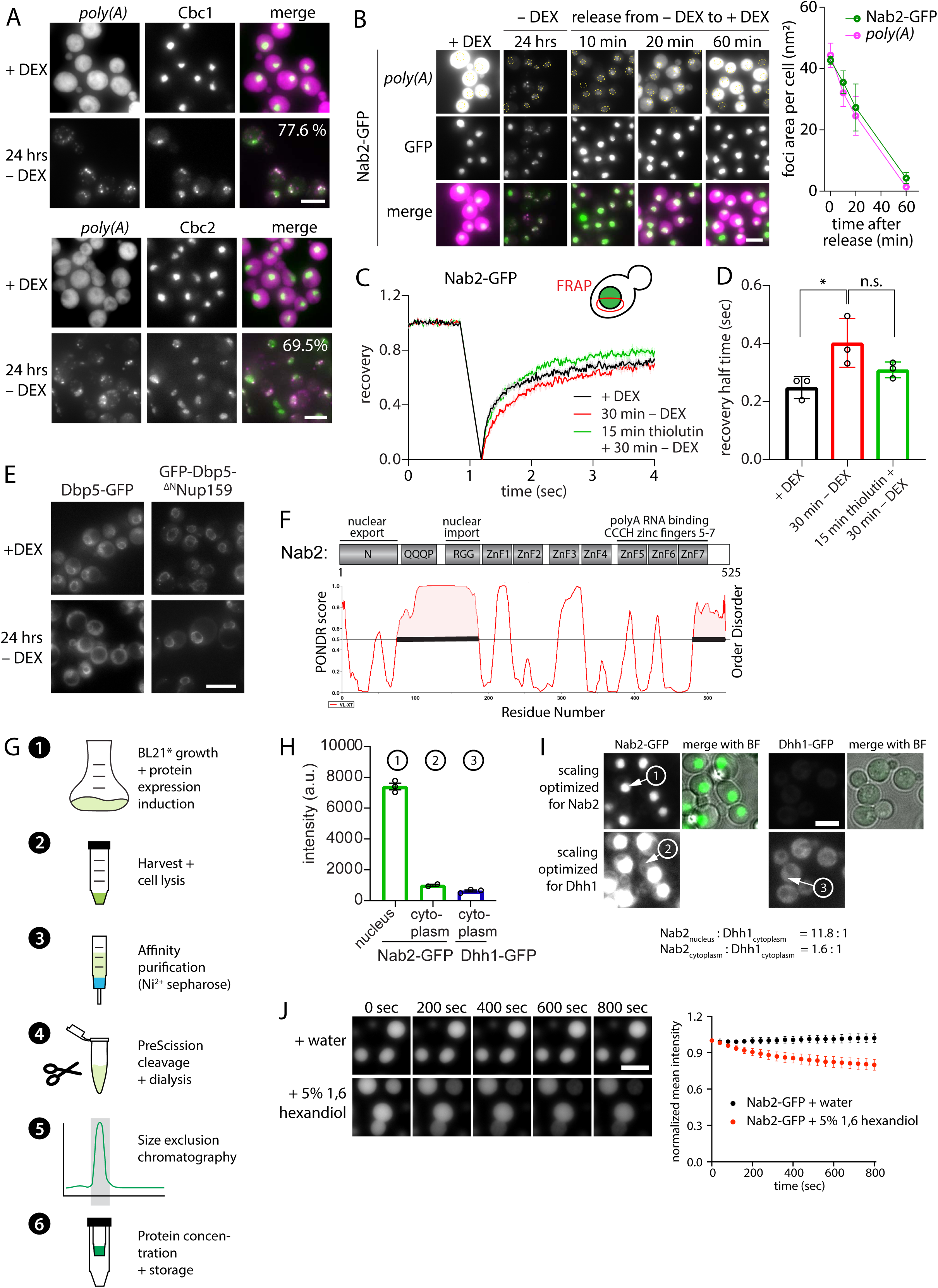
In glucose stress, CBC components co-localize with *poly(A)* RNA foci, and Nab2-GFP focus formation is reversible; *in vitro* Nab2-GFP condensates are mildly sensitive towards addition of 1,6 hexanediol; related to Figure 2 and 3. A) RNA FISH against poly(A) RNA using an oligo(dT)30 probe was performed in fixed cells expressing Cbc1-GFP or Cbc2-GFP. Samples were harvested in +DEX or after acute starvation in medium without DEX for 24 hrs. Percentages indicate co-localization of microscopically visible nuclear GFP foci with microscopically visible poly(A) RNA foci. Scale bar 5 µm. B) RNA FISH against poly(A) RNA using an oligo(dT)30 probe was performed in fixed cells expressing Nab2-GFP. Samples were taken in +DEX conditions, after acute shift and incubation for 24 hrs in –DEX, and after release from –DEX starvation. Dotted yellow line indicates nuclear outline. Right: Quantification of Nab2-GFP foci and RNA foci area per cell. Scale bar 5 µm. C) FRAP of Nab2-GFP cells in +DEX, 30 min –DEX or pre-treated for 15 min with 3 µg/mL thiolutin before starving cells for 30 min in –DEX. FRAP curves are normalized to prebleach and postbleach values. Mean curves ± SEM of three biological replicates with ≥14 cells per replicate are shown. D) Half time of recovery retrieved from fitting individual FRAP curves from C) with a single component model. Shown are means ± SEM of three biological replicates. * p=0.0454, ns p=0.1437, unpaired t-test. E) Representative images of wild type (Dbp5-GFP) or GFP-Dbp5-^ΔN^Nup159 (‘GFP-Dbp5 tether’) cells in +DEX or 24 hrs –DEX. Scale bar 5 µm. F) Domain architecture of Nab2 (top), together with PONDR prediction of naturally disordered regions (bottom) (VL-XT predictor, trained on *V*ariously characterized *L*ong disordered regions and two trained on *X*-ray characterized *T*erminal disordered regions). Black bars indicate predicted disordered regions. G) Schematic of protein expression and purification procedure. (1) *E. coli* BL21* containing pETMCN-based expression vector were grown at 37 °C to OD_600_ 0.6, protein expression was induced with 200 mM IPTG_final_, and the culture was then grown overnight at 18 °C, (2) cells were harvested, resuspended in lysis buffer and lysed with the EmulsiFlex cell homogenizer, (3) 6xHis tagged proteins were extracted using a Ni^2+^ sepharose column, (4) the 6xHis-TwinStrep tag was cleaved with PreScission(3C) protease, and protein was dialysed with a 14 kDa MWCO membrane into storage buffer, (5) size exclusion chromatography was performed with a Superdex 200 column using an AEKTA purifier, and the monomeric peak fractions (grey) were collected, (6) the purified protein was concentrated to approx. 1000 µM with Amicron Centrifugation units (30 kDa MWCO), and 20 µL aliquots were snap-frozen in liquid N_2_ and stored at -80 °C. H) Concentration determination of nuclear and cytoplasmic Nab2-GFP relative to Dhh1-GFP (approx. cellular concentration of 2 µM^1^) as reference for *in vitro* condensate setup. Shown are means ± SEM of three biological replicates. I) Representative images depicting the nuclear and cytoplasmic compartment that was measured, respectively, and the calculated intensity ratios of Nab2-GFP vs. Dhh1-GFP, suggesting that Nab2-GFP has a nuclear concentration of approx. 24 µM. Scale bar 5 µm. J) Recombinant Nab2-GFP droplets were formed in LSB100 buffer to a final KCl concentration of 63 mM in the presence or absence of 0.04 mg/mL polyA RNA. Time lapse imaging at 25 °C was started 60 min after setup upon addition of water or 5% 1,6-hexanediol. Scale bar 5 µm. (Bottom) quantification of mean droplet intensity. Shown are means ± SD of GFP droplet intensity of two biological replicates.

**5).**
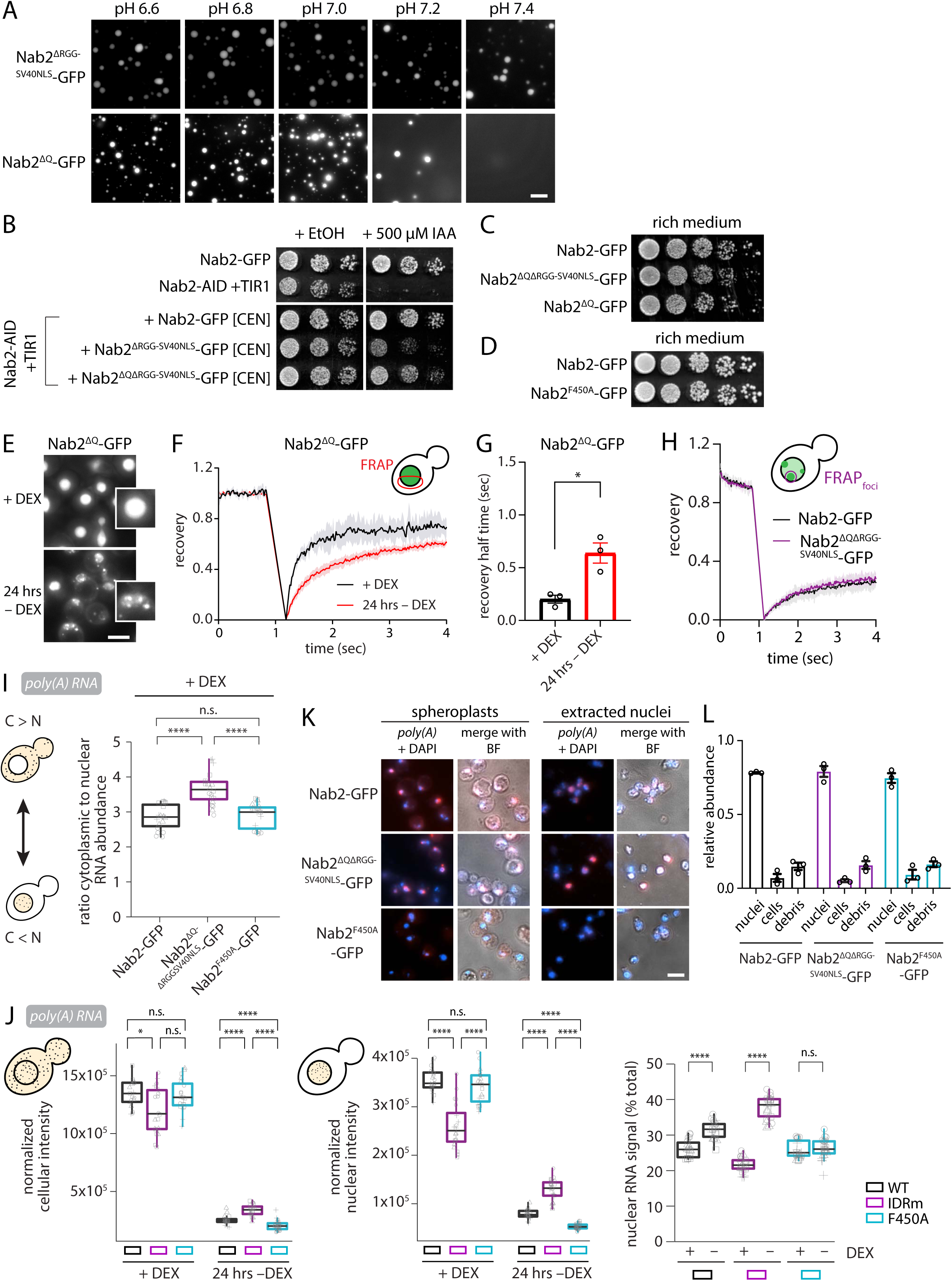
Characterization of Nab2 mutants *in vivo* and *in vitro*, and nucleus isolation efficiency; related to Figure 4 and 5. A) GFP-tagged recombinant Nab2^ΔRGG-SV40NLS^ or Nab2^ΔQ^ mutant droplets were formed in LSB150 buffer at the indicated pH in the presence of 0.04 mg/mL poly(A) RNA and imaged at 25 °C 30 min after setup. Scale bar 10 µm. B) Growth assay on rich medium plates containing 500 µM IAA or solvent control (EtOH). The indicated strains were spotted as 1:5 serial dilutions (starting OD_600_ 0.2) and incubated at 25 °C for 2 days. C-D) Growth assay on rich medium plates. The indicated strains were spotted as 1:5 serial dilutions (starting OD_600_ 0.2) and incubated at 25 °C for 2 days. E) Representative images of Nab2^ΔQ^-GFP cells in +DEX or 24 hrs –DEX. Scale bar 5 µm. F) FRAP of Nab2^ΔQ^-GFP cells in +DEX or 24 hrs –DEX. FRAP curves are normalized to prebleach and postbleach values. Mean curves + SEM of three biological replicates with ≥14 cells per replicate are shown. G) Half time of recovery retrieved from fitting individual FRAP curves from F) with a single component model. Shown are means ± SEM of three biological replicates. * p=0.0132, unpaired t-test. H) 24 hrs –DEX data from Fig. 4F) and G) re-analyzed, including only cells with visible nuclear GFP foci. Bleach region was narrowed over individual GFP foci. FRAP curves are normalized to prebleach and postbleach values. Mean curves ± SEM of three biological replicates with ≥6 cells per replicate are shown. I) Quantification of the ratio of cytoplasmic poly(A) RNA signal intensity versus nuclear poly(A) RNA signal intensity in +DEX from RNA FISH experiment shown in Figure 5A. Data is normalized to median RNA value of all biological replicates. Shown are box plots of four biological replicates (each grey shape corresponds to individual technical replicates within a biological replicate, n>100 cells per replicate). **** p<0.0001, n.s. p=0.47, Wilcoxon test. J) Quantification of the cellular (left) and nuclear (middle) poly(A) RNA signal intensity, as well as the percentage of nuclear poly(A) RNA signal (right), in +DEX and 24 hrs –DEX from RNA FISH experiment shown in Figure 5A. Data is normalized to median RNA value of all biological replicates. Shown are box plots of four biological replicates (each grey shape corresponds to individual technical replicates within a biological replicate, n>100 cells per replicate). **** p<0.0001, * p=0.024 (Nab2-GFP vs. Nab2^ΔQΔRGG-SV40NLS^-GFP, +DEX), n.s. p=0.6 (Nab2-GFP vs. Nab2^F450A^-GFP, +DEX), n.s. p=0.07 (Nab2^ΔQΔRGG-SV40NLS^-GFP vs. Nab2^F450A^-GFP, +DEX), n.s. p=0.23 (Nab2-GFP vs. Nab2^F450A^-GFP, +DEX, nucleus), n.s. p=0.44 (Nab2^F450A^-GFP, +DEX vs. 24 hrs –DEX, percentage nuclear RNA), Wilcoxon test. K) RNA FISH against poly(A) RNA using an oligo(dT)30 probe was performed on spheroplasts or extracted nuclei from indicated strains harvested after 24 hrs starvation in –DEX. BF, bright field. Scale bar 5 µm. L) Quantification of extracted nuclei fraction in K). Shown are means ± SEM of 3 biological replicates.

**6).**
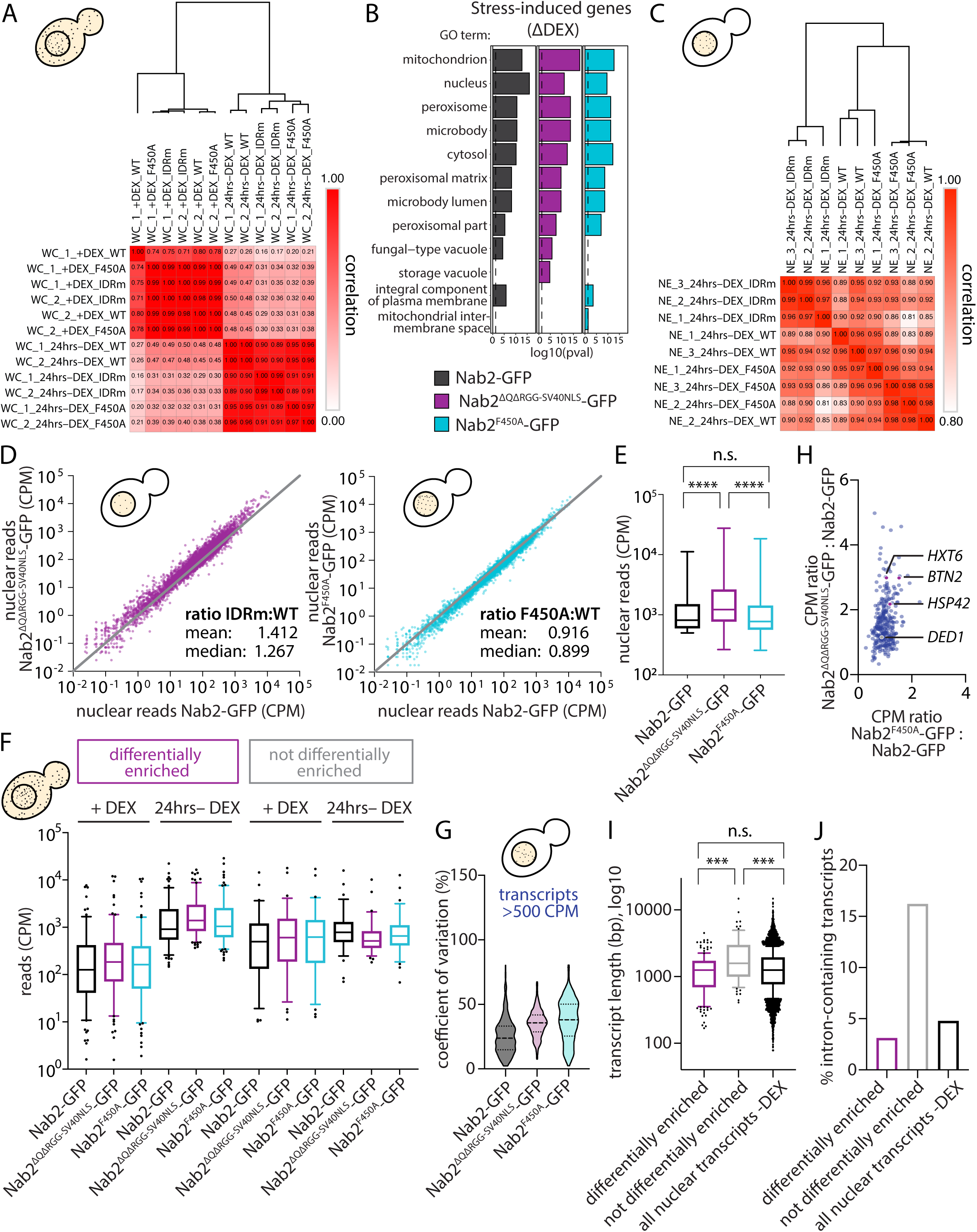
RNA sequencing analysis of whole cell extracts and nuclear extracts; related to Figure 5 and 6. A) Similarity matrix of individual replicates of whole cell RNA sequencing reads, and hierarchical clustering (one minus Pearson correlation) using Morpheus software (Broad Institute, https://software.broadinstitute.org/morpheus). B) GO term analysis of 204 genes upregulated in 24 hrs –DEX compared to +DEX (pval< 0.05 and log2 Fold Change> 3) in the indicated strains. C) Similarity matrix of individual replicates of nuclear RNA sequencing reads, and hierarchical clustering (one minus Pearson correlation) using Morpheus software (Broad Institute, https://software.broadinstitute.org/morpheus). D) Comparison of nuclear RNAseq reads (CPM, counts per million), left: Nab2-GFP vs. Nab2^ΔQΔRGG-SV40NLS^-GFP, right: Nab2-GFP vs. Nab2^F450A^-GFP. Each dot represents one gene. Grey line represents a 1:1 ratio. Shown are average CPMs of three biological replicates. E) Distribution of average reads of most abundant genes (>500 CPM in Nab2-GFP, 318 genes) from D). Box-and-whiskers plot, min to max. **** p<0.001, ns p=0.0578, unpaired t-test. F) Comparison of whole cell RNAseq reads (CPM, counts per million) of differentially enriched vs. not differentially enriched genes of most abundant genes (>500 CPM in Nab2-GFP, 318 genes). Box-and-whisker plot, 10^th^ to 90^th^ percentile. G) Coefficient of variation of most abundant genes (>500 CPM in Nab2-GFP average, 318 genes) determined from RNA sequencing reads of 3 biological replicates. H) Most abundant reads (blue dots, >500 CPM in Nab2-GFP average of 3 biological replicates, representing 42% of the total reads), from Figure 5D, highlighting the 4 mRNAs analyzed with smFISH in Figure 6 and 7. I) Comparison of transcript length (log10) of differentially enriched (161 genes) vs. not differentially enriched (74 genes) vs. all transcripts (5506). Box-and-whiskers plot, 10^th^ to 90^th^ percentile. *** p<0.008, ns p=0.3182, Mann Whitney test. J) Comparison of intron-containing transcripts of differentially enriched (3 out of 149 genes) vs. not differentially enriched (14 out of 100 genes) vs. all transcripts (263 out of 5506).

**7).**
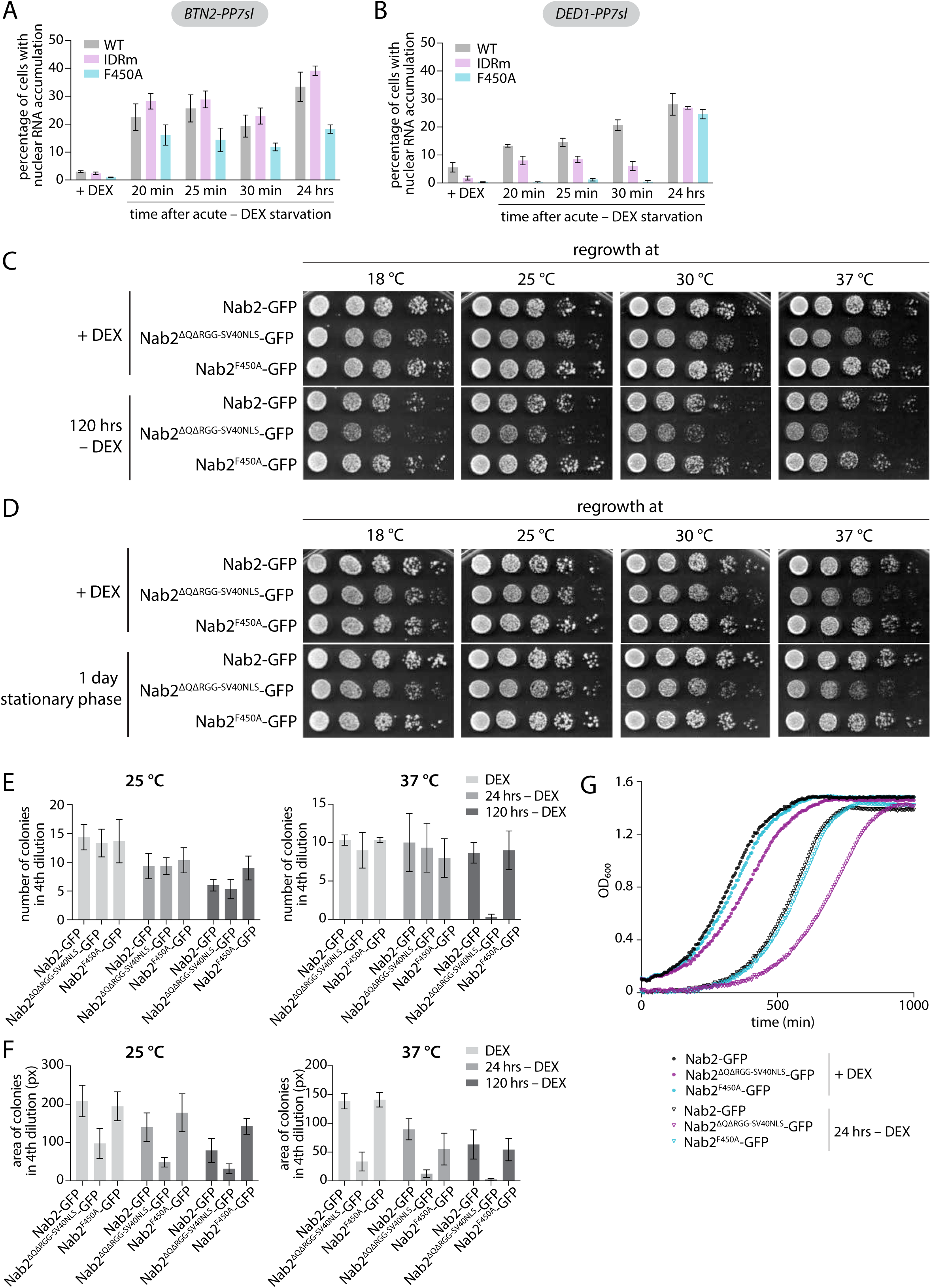
Correct Nab2 condensation is essential for survival after glucose stress, both acutely and in stationary phase; related to Figure 7. A) Live imaging of PP7sl-tagged *BTN2* mRNA in cells expressing Nab2-GFP, Nab2^ΔQΔRGG-SV40NLS^-GFP or Nab2^F450A^-GFP in +DEX and after acute shift into –DEX. Quantification of nuclear RNA accumulation, i.e. percentage of cells where average RNA signal intensity is larger in nucleus than in cytoplasm. Shown are means ± SEM of three biological replicates, n>100 cells per replicate. +DEX and 24 hrs –DEX data are also shown in Figure 7E. B) Live imaging of PP7sl-tagged *DED1* mRNA in cells expressing Nab2-GFP, Nab2^ΔQΔRGG-SV40NLS^-GFP or Nab2^F450A^-GFP in +DEX and after acute shift into –DEX. Quantification of nuclear RNA accumulation, i.e. percentage of cells where average RNA signal intensity is larger in nucleus than in cytoplasm. Shown are means ± SEM of three biological replicates, n>100 cells per replicate. +DEX and 24 hrs –DEX data are also shown in Figure 7F. C) Growth recovery assay of cells expressing Nab2-GFP, Nab2^ΔQΔRGG-SV40NLS^-GFP or Nab2^F450A^-GFP. Mid-log cultures grown at 25 °C in +DEX or acutely starved in medium without DEX for 120 hrs were recovered as 1:5 serial dilutions (starting OD_600_ 0.2) on rich medium plates containing glucose at either 18 °C for 4 days, 25 °C for 2 days, 30 °C for 1 day or 37 °C for 1 day. D) Growth recovery assay of cells expressing Nab2-GFP, Nab2^ΔQΔRGG-SV40NLS^-GFP or Nab2^F450A^-GFP. Cultures in +DEX were grown from mid-log into stationary phase for 1 day at 25 °C and subsequently recovered as 1:5 serial dilutions (starting OD_600_ 0.2) on rich medium plates containing glucose at either 18 °C for 4 days, 25 °C for 2 days, 30 °C for 1 day or 37 °C for 1 day. E) Quantification of growth recovery assay shown in Figure 7G. Number of individual colonies counted in 4^th^ dilution; left: 25 °C, right: 37 °C. Shown are means ± SEM of three biological replicates. F) Quantification of growth recovery assay shown in Figure 7G. Area of all colonies determined in 4^th^ dilution; left: 25 °C, right: 37 °C. Shown are means ± SEM of three biological replicates. G) Growth recovery assay in liquid medium of cells expressing Nab2-GFP, Nab2^ΔQΔRGG-SV40NLS^-GFP or Nab2^F450A^-GFP. Mid-log cultures grown at 25 °C in +DEX or acutely starved in medium without DEX 24 hrs were back-diluted in synthetic complete medium containing glucose and culture density (OD_600_) was measured every 5 min for 16.7 hrs. The starting OD_600_ of +DEX cultures was 5x higher than for the –DEX cultures. Shown are representative growth curves from 1 of 3 biological replicates.

**Video S1** Cells expressing GFA1-PP7sl, PP7CP-GFP, the nuclear rim marker Ndc1and the Dbp5-degron without TIR1 are treated for 1 hr with Auxin (IAA) before imaging using a custom-built microscope^2^. Each frame is the average projection of 10 consecutive images. Scale bar 5 µm. Related to Figure 1.

**Video S2** Cells expressing GFA1-PP7sl, PP7CP-GFP, the nuclear rim marker Ndc1and the Dbp5-degron with TIR1 are treated for 1 hr with Auxin (IAA) before imaging using a custom-built microscope^2^. Each frame is the average projection of 10 consecutive images. Scale bar 5 µm. Related to Figure 1.

**Video S3** Recombinant Nab2-GFP droplets were formed in LSB100 buffer to a final KCl concentration of 63 mM in the presence of 0.04 mg/mL poly(A) RNA and imaged every 2 min for 2 hrs at 25 °C directly after setup. Scale bar 10 µm. Related to Figure 2.

**Video S4** Live imaging experiment shown in Figure 7E of PP7sl-tagged *BTN2* mRNA in cells expressing Nab2-GFP after acute shift and incubation for 24 hrs in –DEX. Representative movies show 2 cells with dynamic *BTN2-PP7sl* mRNAs. Scale bar 2 µm. Related to Figure 7.

**Video S5** Live imaging experiment shown in Figure 7E of PP7sl-tagged *BTN2* mRNA in cells expressing Nab2-GFP after acute shift and incubation for 24 hrs in –DEX. Representative movies show 2 cells with static *BTN2-PP7sl* mRNAs. Scale bar 2 µm. Related to Figure 7.

## Notes

### Summary of Updates

Title changed Main text revised Figure 5, 6 and 7 revised

## References

1. Ashkenazy-Titelman, A., Shav-Tal, Y., and Kehlenbach, R.H. (2020). Into the basket and be-yond: the journey of mRNA through the nuclear pore complex. Biochem. J. 477, 23–44. 10.1042/BCJ20190132.

2. De Magistris, P. (2021). The Great Escape: mRNA Export through the Nuclear Pore Complex. Int. J. Mol. Sci. 22, 11767. 10.3390/ijms222111767.

3. Herzel, L., Ottoz, D.S.M., Alpert, T., and Neugebauer, K.M. (2017). Splicing and transcription touch base: co-transcriptional spliceosome assembly and function. Nat. Rev. Mol. Cell Biol. 18, 637– 650. 10.1038/nrm.2017.63.

4. Rambout, X., and Maquat, L.E. (2020). The nuclear cap-binding complex as choreographer of gene transcription and pre-mRNA processing. Genes Dev. 34, 1113–1127. 10.1101/gad.339986.120.

5. Shi, Y., and Manley, J.L. (2015). The end of the message: multiple protein–RNA interactions define the mRNA polyadenylation site. Genes Dev. 29, 889–897. 10.1101/gad.261974.115.

6. Singh, G., Pratt, G., Yeo, G.W., and Moore, M.J. (2015). The Clothes Make the mRNA: Past and Present Trends in mRNP Fashion. Annu. Rev. Biochem. 84, 325–354. 10.1146/annurev-biochem-080111-092106.

7. Izaurralde, E., Lewis, J., McGuigan, C., Jankowska, M., Darzynkiewicz, E., and Mattaj, I.W. (1994). A nuclear cap binding protein complex involved in pre-mRNA splicing. Cell 78, 657–668. 10.1016/0092-8674(94)90530-4.

8. Katahira, J. (2012). mRNA export and the TREX complex. Biochim. Biophys. Acta BBA - Gene Regul. Mech. 1819, 507–513. 10.1016/j.bbagrm.2011.12.001.

9. Pühringer, T., Hohmann, U., Fin, L., Pacheco-Fiallos, B., Schellhaas, U., Brennecke, J., and Plaschka, C. (2020). Structure of the human core transcription-export complex reveals a hub for multi-valent interactions. eLife 9, e61503. 10.7554/eLife.61503.

10. Sträßer, K., Masuda, S., Mason, P., Pfannstiel, J., Oppizzi, M., Rodriguez-Navarro, S., Rondón, A.G., Aguilera, A., Struhl, K., Reed, R., et al. (2002). TREX is a conserved complex coupling transcrip-tion with messenger RNA export. Nature 417, 304–308. 10.1038/nature746.

11. Sträßer, K., and Hurt, E. (2001). Splicing factor Sub2p is required for nuclear mRNA export through its interaction with Yra1p. Nature 413, 648–652. 10.1038/35098113.

12. Wegener, M., and Müller-McNicoll, M. (2019). View from an mRNP: The Roles of SR Proteins in Assembly, Maturation and Turnover. In The Biology of mRNA: Structure and Function Advances in Experimental Medicine and Biology., M. Oeffinger and D. Zenklusen, eds. (Springer International Publishing), pp. 83–112. 10.1007/978-3-030-31434-7_3.

13. Anderson, J.T., Wilson, S.M., Datar, K.V., and Swanson, M.S. (1993). NAB2: a yeast nuclear polyadenylated RNA-binding protein essential for cell viability. Mol. Cell. Biol. 13, 2730–2741. 10.1128/mcb.13.5.2730.

14. Hector, R.E., Nykamp, K.R., Dheur, S., Anderson, J.T., Non, P.J., Urbinati, C.R., Wilson, S.M., Minvielle-Sebastia, L., and Swanson, M.S. (2002). Dual requirement for yeast hnRNP Nab2p in mRNA poly(A) tail length control and nuclear export. EMBO J. 21, 1800–1810. 10.1093/emboj/21.7.1800.

15. Schmid, M., Poulsen, M.B., Olszewski, P., Pelechano, V., Saguez, C., Gupta, I., Steinmetz, L.M., Moore, C., and Jensen, T.H. (2012). Rrp6p Controls mRNA Poly(A) Tail Length and Its Decoration with Poly(A) Binding Proteins. Mol. Cell 47, 267–280. 10.1016/j.molcel.2012.05.005.

16. Schmid, M., Olszewski, P., Pelechano, V., Gupta, I., Steinmetz, L.M., and Jensen, T.H. (2015). The Nuclear PolyA-Binding Protein Nab2p Is Essential for mRNA Production. Cell Rep. 47, 267–280. 10.1016/j.celrep.2015.06.008.

17. Aibara, S., Gordon, J.M.B., Riesterer, A.S., McLaughlin, S.H., and Stewart, M. (2017). Structural basis for the dimerization of Nab2 generated by RNA binding provides insight into its contribution to both poly(A) tail length determination and transcript compaction in Saccharomyces cerevisiae. Nucleic Acids Res. 45, 1529–1538. 10.1093/nar/gkw1224.

18. Alpert, T., Straube, K., Oesterreich, F.C., and Neugebauer, K.M. (2020). Widespread Transcrip-tional Readthrough Caused by Nab2 Depletion Leads to Chimeric Transcripts with Retained Introns. Cell Rep. 33, 108324. 10.1016/j.celrep.2020.108324.

19. Fasken, M.B., Corbett, A.H., and Stewart, M. (2019). Structure–function relationships in the Nab2 polyadenosine-RNA binding Zn finger protein family. Protein Sci. 28, 513–523. 10.1002/pro.3565.

20. Green, D.M., Johnson, C.P., Hagan, H., and Corbett, A.H. (2003). The C-terminal domain of myosin-like protein 1 (Mlp1p) is a docking site for heterogeneous nuclear ribonucleoproteins that are required for mRNA export. Proc. Natl. Acad. Sci. 100, 1010–1015. 10.1073/pnas.0336594100.

21. Aitchison, J.D., Blobel, G., and Rout, M.P. (1996). Kap104p: A Karyopherin Involved in the Nuclear Transport of Messenger RNA Binding Proteins. Science 274, 624–627. 10.1126/sci-ence.274.5287.624.

22. Derrer, C.P., Mancini, R., Vallotton, P., Huet, S., Weis, K., and Dultz, E. (2019). The RNA ex-port factor Mex67 functions as a mobile nucleoporin. J. Cell Biol. 218 (*12*), 3967–3976. 10.1083/jcb.201909028.

23. Alcázar-Román, A.R., Tran, E.J., Guo, S., and Wente, S.R. (2006). Inositol hexakisphosphate and Gle1 activate the DEAD-box protein Dbp5 for nuclear mRNA export. Nat. Cell Biol. 8, 711–716. 10.1038/ncb1427.

24. Tran, E.J., Zhou, Y., Corbett, A.H., and Wente, S.R. (2007). The DEAD-Box Protein Dbp5 Controls mRNA Export by Triggering Specific RNA:Protein Remodeling Events. Mol. Cell 28, 850– 859. 10.1016/j.molcel.2007.09.019.

25. Weirich, C.S., Erzberger, J.P., Flick, J.S., Berger, J.M., Thorner, J., and Weis, K. (2006). Activa-tion of the DExD/H-box protein Dbp5 by the nuclear-pore protein Gle1 and its coactivator InsP6 is required for mRNA export. Nat. Cell Biol. 8, 668–676. 10.1038/ncb1424.

26. Montpetit, B., Thomsen, N.D., Helmke, K.J., Seeliger, M.A., Berger, J.M., and Weis, K. (2011). A conserved mechanism of DEAD-box ATPase activation by nucleoporins and InsP6 in mRNA ex-port. Nature 472, 238–242. 10.1038/nature09862.

27. Noble, K.N., Tran, E.J., Alcázar-Román, A.R., Hodge, C.A., Cole, C.N., and Wente, S.R. (2011). The Dbp5 cycle at the nuclear pore complex during mRNA export II: nucleotide cycling and mRNP remodeling by Dbp5 are controlled by Nup159 and Gle1. Genes Dev. 25, 1065–1077. 10.1101/gad.2040611.

28. Adams, R.L., and Wente, S.R. (2020). Dbp5 associates with RNA-bound Mex67 and Nab2 and its localization at the nuclear pore complex is sufficient for mRNP export and cell viability. PLOS Ge-net. 16, e1009033. 10.1371/journal.pgen.1009033.

29. Kaminski, T., Siebrasse, J.P., and Kubitscheck, U. (2013). A single molecule view on Dbp5 and mRNA at the nuclear pore. Nucleus 4, 8–13. 10.4161/nucl.23386.

30. Zhao, J., Jin, S.-B., Björkroth, B., Wieslander, L., and Daneholt, B. (2002). The mRNA export factor Dbp5 is associated with Balbiani ring mRNP from gene to cytoplasm. EMBO J. 21, 1177–1187. 10.1093/emboj/21.5.1177.

31. Banani, S.F., Lee, H.O., Hyman, A.A., and Rosen, M.K. (2017). Biomolecular condensates: or-ganizers of cellular biochemistry. Nat. Rev. Mol. Cell Biol. 18, 285–298. 10.1038/nrm.2017.7.

32. Shin, Y., and Brangwynne, C.P. (2017). Liquid phase condensation in cell physiology and dis-ease. Science 357, eaaf4382. 10.1126/science.aaf4382.

33. Roden, C., and Gladfelter, A.S. (2021). RNA contributions to the form and function of bio-molecular condensates. Nat. Rev. Mol. Cell Biol. 22, 183–195. 10.1038/s41580-020-0264-6.

34. Hondele, M., Sachdev, R., Heinrich, S., Wang, J., Vallotton, P., Fontoura, B.M.A., and Weis, K. (2019). DEAD-box ATPases are global regulators of phase-separated organelles. Nature, 573, 144–148. 10.1038/s41586-019-1502-y.

35. Riback, J.A., Katanski, C.D., Kear-Scott, J.L., Pilipenko, E.V., Rojek, A.E., Sosnick, T.R., and Drummond, D.A. (2017). Stress-Triggered Phase Separation Is an Adaptive, Evolutionarily Tuned Re-sponse. Cell 168, 1028–1040.e19. 10.1016/j.cell.2017.02.027.

36. Kroschwald, S., Maharana, S., Mateju, D., Malinovska, L., Nüske, E., Poser, I., Richter, D., and Alberti, S. (2015). Promiscuous interactions and protein disaggregases determine the material state of stress-inducible RNP granules. eLife 4, e06807. 10.7554/eLife.06807.

37. Sheth, U., and Parker, R. (2003). Decapping and Decay of Messenger RNA Occur in Cytoplas-mic Processing Bodies. Science 300, 805–808. 10.1126/science.1082320.

38. Rajoo, S., Vallotton, P., Onischenko, E., and Weis, K. (2018). Stoichiometry and compositional plasticity of the yeast nuclear pore complex revealed by quantitative fluorescence microscopy. Proc. Natl. Acad. Sci., 201719398. 10.1073/pnas.1719398115.

39. Neville, M., and Rosbash, M. (1999). The NES-Crm1p export pathway is not a major mRNA export route in Saccharomyces cerevisiae. EMBO J. 18, 3746–3756. 10.1093/emboj/18.13.3746.

40. Hodge, C.A., Colot, H.V., Stafford, P., and Cole, C.N. (1999). Rat8p/Dbp5p is a shuttling transport factor that interacts with Rat7p/Nup159p and Gle1p and suppresses the mRNA export de-fect of xpo1-1 cells. EMBO J. 18, 5778–5788. 10.1093/emboj/18.20.5778.

41. Snay-Hodge, C.A., Colot, H.V., Goldstein, A.L., and Cole, C.N. (1998). Dbp5p/Rat8p is a yeast nuclear pore-associated DEAD-box protein essential for RNA export. EMBO J. 17, 2663–2676. 10.1093/emboj/17.9.2663.

42. Larson, D.R., Zenklusen, D., Wu, B., Chao, J.A., and Singer, R.H. (2011). Real-Time Observa-tion of Transcription Initiation and Elongation on an Endogenous Yeast Gene. Science 332, 475–478. 10.1126/science.1202142.

43. Smith, C., Lari, A., Derrer, C.P., Ouwehand, A., Rossouw, A., Huisman, M., Dange, T., Hop-man, M., Joseph, A., Zenklusen, D., et al. (2015). In vivo single-particle imaging of nuclear mRNA ex-port in budding yeast demonstrates an essential role for Mex67p. J. Cell Biol. 211, 1121–1130. 10.1083/jcb.201503135.

44. McSwiggen, D.T., Mir, M., Darzacq, X., and Tjian, R. (2019). Evaluating phase separation in live cells: diagnosis, caveats, and functional consequences. Genes Dev. 33, 1619–1634. 10.1101/gad.331520.119.

45. Izawa, S., Takemura, R., Ikeda, K., Fukuda, K., Wakai, Y., and Inoue, Y. (2005). Characteriza-tion of Rat8 localization and mRNA export in Saccharomyces cerevisiae during the brewing of Japa-nese sake. Appl. Microbiol. Biotechnol. 69, 86–91. 10.1007/s00253-005-1954-x.

46. Takemura, R., Inoue, Y., and Izawa, S. (2004). Stress response in yeast mRNA export factor: reversible changes in Rat8p localization are caused by ethanol stress but not heat shock. J. Cell Sci. 117, 4189–4197. 10.1242/jcs.01296.

47. Tipper, D.J. (1973). Inhibition of Yeast Ribonucleic Acid Polymerases by Thiolutin. J. Bacteriol. 116(*1*), 245–56. 10.1128/jb.116.1.245-256.1973.

48. Marfatia, K.A., Crafton, E.B., Green, D.M., and Corbett, A.H. (2003). Domain Analysis of the Saccharomyces cerevisiae Heterogeneous Nuclear Ribonucleoprotein, Nab2p Dissecting The Require-ments For Nab2p-Facilitated Poly(A) RNA Export. J. Biol. Chem. 278, 6731–6740. 10.1074/jbc.M207571200.

49. Soniat, M., Sampathkumar, P., Collett, G., Gizzi, A.S., Banu, R.N., Bhosle, R.C., Chamala, S., Chowdhury, S., Fiser, A., Glenn, A.S., et al. (2013). Crystal structure of human Karyopherin β2 bound to the PY-NLS of Saccharomyces cerevisiae Nab2. J. Struct. Funct. Genomics 14, 31–35. 10.1007/s10969-013-9150-1.

50. Li, X., Romero, P., Rani, M., Dunker, A.K., and Obradovic, Z. (1999). Predicting Protein Dis-order for N-, C-, and Internal Regions. Genome Inform. Workshop Genome Inform. 10, 30–40.

51. Romero, P., Obradovic, Z., and Dunker, A.K. (1997). Sequence Data Analysis for Long Disor-dered Regions Prediction in the Calcineurin Family. Genome Inform. Workshop Genome Inform. 8, 110–124.

52. Tuck, A.C., and Tollervey, D. (2013). A Transcriptome-wide Atlas of RNP Composition Re-veals Diverse Classes of mRNAs and lncRNAs. Cell 154, 996–1009. 10.1016/j.cell.2013.07.047.

53. Niepel, M., Farr, J.C., Rout, M.P., and Strambio-De-Castillia, C. (2017). Rapid isolation of func-tionally intact nuclei from the yeast Saccharomyces. bioRxiv, 162388. 10.1101/162388.

54. Tudek, A., Schmid, M., Makaras, M., Barrass, J.D., Beggs, J.D., and Jensen, T.H. (2018). A Nu-clear Export Block Triggers the Decay of Newly Synthesized Polyadenylated RNA. Cell Rep. 24, 2457–2467.e7. 10.1016/j.celrep.2018.07.103.

55. Soucek, S., Zeng, Y., Bellur, D.L., Bergkessel, M., Morris, K.J., Deng, Q., Duong, D., Seyfried, N.T., Guthrie, C., Staley, J.P., et al. (2016). Evolutionarily Conserved Polyadenosine RNA Binding Pro-tein Nab2 Cooperates with Splicing Machinery To Regulate the Fate of Pre-mRNA. Mol. Cell. Biol. 36, 2697–2714. 10.1128/MCB.00402-16.

56. Khong, A., Matheny, T., Jain, S., Mitchell, S.F., Wheeler, J.R., and Parker, R. (2017). The Stress Granule Transcriptome Reveals Principles of mRNA Accumulation in Stress Granules. Mol. Cell 68, 808–820.e5. 10.1016/j.molcel.2017.10.015.

57. Mauger, O., Lemoine, F., and Scheiffele, P. (2016). Targeted Intron Retention and Excision for Rapid Gene Regulation in Response to Neuronal Activity. Neuron 92, 1266–1278. 10.1016/j.neu-ron.2016.11.032.

58. Bahar Halpern, K., Caspi, I., Lemze, D., Levy, M., Landen, S., Elinav, E., Ulitsky, I., and Itz-kovitz, S. (2015). Nuclear Retention of mRNA in Mammalian Tissues. Cell Rep. 13, 2653–2662. 10.1016/j.celrep.2015.11.036.

59. Zimmerli, C.E., Allegretti, M., Rantos, V., Goetz, S.K., Obarska-Kosinska, A., Zagoriy, I., Hala-vatyi, A., Hummer, G., Mahamid, J., Kosinski, J., et al. (2021). Nuclear pores dilate and constrict in cel-lulo. Science 374, eabd9776. 10.1126/science.abd9776.

60. Hochberg-Laufer, H., Schwed-Gross, A., Neugebauer, K.M., and Shav-Tal, Y. (2019). Uncou-pling of nucleo-cytoplasmic RNA export and localization during stress. Nucleic Acids Res. 47, 4778– 4797. 10.1093/nar/gkz168.

61. Tauber, D., Tauber, G., Khong, A., Treeck, B.V., Pelletier, J., and Parker, R. (2020). Modulation of RNA Condensation by the DEAD-Box Protein eIF4A. Cell 180(*3*), 411–426. 10.1016/j.cell.2019.12.031.

62. Iserman, C., Desroches Altamirano, C., Jegers, C., Friedrich, U., Zarin, T., Fritsch, A.W., Mitta-sch, M., Domingues, A., Hersemann, L., Jahnel, M., et al. (2020). Condensation of Ded1p Promotes a Translational Switch from Housekeeping to Stress Protein Production. Cell 181, 818–831.e19. 10.1016/j.cell.2020.04.009.

63. Yang, P., Mathieu, C., Kolaitis, R.-M., Zhang, P., Messing, J., Yurtsever, U., Yang, Z., Wu, J., Li, Y., Pan, Q., et al. (2020). G3BP1 Is a Tunable Switch that Triggers Phase Separation to Assemble Stress Granules. Cell 181, 325–345.e28. 10.1016/j.cell.2020.03.046.

64. Dechant, R., Saad, S., Ibáñez, A.J., and Peter, M. (2014). Cytosolic pH Regulates Cell Growth through Distinct GTPases, Arf1 and Gtr1, to Promote Ras/PKA and TORC1 Activity. Mol. Cell 55, 409–421. 10.1016/j.molcel.2014.06.002.

65. Joyner, R.P., Tang, J.H., Helenius, J., Dultz, E., Brune, C., Holt, L.J., Huet, S., Müller, D.J., and Weis, K. (2016). A glucose-starvation response regulates the diffusion of macromolecules. eLife 5, e09376. 10.7554/eLife.09376.

66. Martínez-Muñoz, G.A., and Kane, P. (2008). Vacuolar and Plasma Membrane Proton Pumps Collaborate to Achieve Cytosolic pH Homeostasis in Yeast. J. Biol. Chem. 283, 20309–20319. 10.1074/jbc.M710470200.

67. Orij, R., Postmus, J., Beek, A.T., Brul, S., and Smits, G.J. (2009). In vivo measurement of cyto-solic and mitochondrial pH using a pH-sensitive GFP derivative in Saccharomyces cerevisiae reveals a relation between intracellular pH and growth. Microbiology 155, 268–278. 10.1099/mic.0.022038-0.

68. Garcia, J.F., and Parker, R. (2015). MS2 coat protein bound to yeast mRNAs block 5′ to 3′ deg-radation and trap mRNA decay products: implications for the localization of mRNAs by MS2-MCP system. RNA 21,1–3. 10.1261/rna.051797.115.

69. Heinrich, S., Sidler, C.L., Azzalin, C.M., and Weis, K. (2017). Stem–loop RNA labeling can af-fect nuclear and cytoplasmic mRNA processing. RNA 23, 134–141. 10.1261/rna.057786.116.

70. Leung, S.W., Apponi, L.H., Cornejo, O.E., Kitchen, C.M., Valentini, S.R., Pavlath, G.K., Dun-ham, C.M., and Corbett, A.H. (2009). Splice variants of the human ZC3H14 gene generate multiple isoforms of a zinc finger polyadenosine RNA binding protein. Gene 439, 71–78. 10.1016/j.gene.2009.02.022.

71. Longtine, M.S., Mckenzie III, A., Demarini, D.J., Shah, N.G., Wach, A., Brachat, A., Philippsen, P., and Pringle, J.R. (1998). Additional modules for versatile and economical PCR-based gene deletion and modification in Saccharomyces cerevisiae. Yeast 14, 953–961. 10.1002/(SICI)1097-0061(199807)14:10<953::AID-YEA293>3.0.CO;2-U.

72. Laughery, M.F., Hunter, T., Brown, A., Hoopes, J., Ostbye, T., Shumaker, T., and Wyrick, J.J. (2015). New vectors for simple and streamlined CRISPR-Cas9 genome editing in Saccharomyces cere-visiae. Yeast Chichester Engl. 32, 711–720. 10.1002/yea.3098.

73. Krull, A., Buchholz, T.-O., and Jug, F. (2018). Noise2Void - Learning Denoising from Single Noisy Images. ArXiv181110980 Cs.

74. Tinevez, J.-Y., Perry, N., Schindelin, J., Hoopes, G.M., Reynolds, G.D., Laplantine, E., Bedna-rek, S.Y., Shorte, S.L., and Eliceiri, K.W. (2017). TrackMate: An open and extensible platform for sin-gle-particle tracking. Methods San Diego Calif 115, 80–90. 10.1016/j.ymeth.2016.09.016.

75. Allan, D.B., Caswell, T., Keim, N.C., and van der Wel, C.M. (2018). trackpy: Trackpy v0.4.1. 10.5281/zenodo.1226458.

76. Crocker, J.C., and Grier, D.G. (1996). Methods of Digital Video Microscopy for Colloidal Stud-ies. J. Colloid Interface Sci. 179, 298–310. 10.1006/jcis.1996.0217.

77. Tarantino, N., Tinevez, J.-Y., Crowell, E.F., Boisson, B., Henriques, R., Mhlanga, M., Agou, F., Israël, A., and Laplantine, E. (2014). TNF and IL-1 exhibit distinct ubiquitin requirements for inducing NEMO-IKK supramolecular structures. J. Cell Biol. 204, 231–245. 10.1083/jcb.201307172.

78. Wickham, H., Averick, M., Bryan, J., Chang, W., McGowan, L.D., François, R., Grolemund, G., Hayes, A., Henry, L., Hester, J., et al. (2019). Welcome to the Tidyverse. J. Open Source Softw. 4, 1686. 10.21105/joss.01686.

79. Vallotton, P., Rajoo, S., Wojtynek, M., Onischenko, E., Kralt, A., Derrer, C.P., and Weis, K. (2019). Mapping the native organization of the yeast nuclear pore complex using nuclear radial intensity measurements. Proc. Natl. Acad. Sci. 116, 14606–14613. 10.1073/pnas.1903764116.

80. Bancaud, A., Huet, S., Rabut, G., and Ellenberg, J. (2010). Fluorescence Perturbation Tech-niques to Study Mobility and Molecular Dynamics of Proteins in Live Cells: FRAP, Photoactivation, Photoconversion, and FLIP. Cold Spring Harb. Protoc. 2010, pdb.top90. 10.1101/pdb.top90.

81. Aris, J.P., and Blobel, G. (1991). [53] Isolation of yeast nuclei. In Methods in Enzymology Guide to Yeast Genetics and Molecular Biology. (Academic Press), pp. 735–749. 10.1016/0076-6879(91)94056-I.

82. Huang, D.W., Sherman, B.T., and Lempicki, R.A. (2009). Bioinformatics enrichment tools: paths toward the comprehensive functional analysis of large gene lists. Nucleic Acids Res. 37, 1–13. 10.1093/nar/gkn923.

83. Huang, D.W., Sherman, B.T., and Lempicki, R.A. (2009). Systematic and integrative analysis of large gene lists using DAVID bioinformatics resources. Nat. Protoc. 4, 44–57. 10.1038/nprot.2008.211.

84. Heinrich, S., and Hondele, M. (2022). Probing Liquid–Liquid Phase Separation of RNA-Bind-ing Proteins In Vitro and In Vivo. In Alternative Splicing: Methods and Protocols Methods in Molecu-lar Biology., P. Scheiffele and O. Mauger, eds. (Springer US), pp. 307–333. 10.1007/978-1-0716-2521-7_18.

85. Nishimura, K., Fukagawa, T., Takisawa, H., Kakimoto, T., and Kanemaki, M. (2009). An auxin-based degron system for the rapid depletion of proteins in nonplant cells. Nat. Methods 6, 917–922. 10.1038/nmeth.1401.

## References

1. Ho, B., Baryshnikova, A., and Brown, G.W. (2018). Unification of Protein Abundance Datasets Yields a Quantitative Saccharomyces cerevisiae Proteome. Cell Systems 6, 192–205 10.1016/j.cels.2017.12.004.

2. Smith, C., Lari, A., Derrer, C.P., Ouwehand, A., Rossouw, A., Huisman, M., Dange, T., Hopman, M., Joseph, A., Zenklusen, D., et al. (2015). In vivo single-particle imaging of nuclear mRNA export in budding yeast demonstrates an essential role for Mex67p. J Cell Biol 211, 1121–1130. 10.1083/jcb.201503135.

